# Haplotype-resolved centromeric chromatin organization from a complete diploid human genome

**DOI:** 10.64898/2026.03.27.714900

**Authors:** Yuan Xu, Hailey Loucks, Julian Menendez, Fedor Ryabov, Julian K. Lucas, Monika Cechova, Luke Morina, Emily Xu, Danilo Dubocanin, Cy Chittenden, Mobin Asri, Ivo Violich, Christian Ortiz, Joshua M.V. Gardner, Todd Hillaker, Sara O’Rourke, Brandy McNulty, Tamara A Potapova, Matthew W. Mitchell, Jacob P. Schwartz, Aaron F. Straight, Jennifer L. Gerton, Winston Timp, Ivan A. Alexandrov, Nicolas Altemose, Karen H. Miga

## Abstract

Centromeres ensure proper chromosome segregation during cell division, yet the organization and regulation of centromeric chromatin within satellite DNA arrays remain incompletely understood. Here, we leverage the complete diploid human genome benchmark (T2T-HG002) to provide a detailed study of centromeric sequence and chromatin architecture on individual haplotypes. Using adaptive-sampling-enriched, ultra-long-read DiMeLo-seq, we achieve single-molecule chromatin profiling across all centromeres, revealing that along single chromatin fibers, CENP-A, the histone variant specifying centromere identity, forms multiple discrete subdomains within hypomethylated centromere dip regions (CDRs) that are flanked by H3K9me3-enriched heterochromatin. Despite underlying sequence variation, CDRs localize to sequence-homogeneous domains and maintain relatively balanced CENP-A dosage and aggregate length across all chromosomes and between haplotypes. Further, we show that bidirectional changes to centromeric and pericentromeric DNA methylation are accompanied by changes to centromeric chromatin architecture. In passaged cells with centromeric hypomethylation, subdomain boundaries are eroded, and adjacent CENP-A domains tend to merge and expand. Conversely, in pluripotent stem cells with centromeric hypermethylation, CDRs are fundamentally reorganized, such that discrete hypomethylated domains are frequently consolidated into broader contiguous tracts. These methylation-associated CDR restructuring events suggest that DNA methylation acts as a principal regulator of human centromere organization, with implications for understanding centromere plasticity, epigenetic inheritance, and chromosomal instability in development and disease.

## Introduction

Centromeres are essential genomic loci that ensure accurate chromosome segregation, yet fundamental questions about their chromatin architecture remain unresolved ^1^. Centromere protein A (CENP-A), a histone H3 variant, epigenetically specifies centromere identity and directs kinetochore assembly ^2,3^. Rather than forming a single contiguous domain, CENP-A chromatin is organized into multiple discrete subdomains interspersed with canonical histone H3 ^4^, a pattern proposed to support the higher-order three-dimensional organization that underlies kinetochore structure and function ^5,6^. CENP-A overexpression, often observed in cancer, can alter domain boundaries and destabilize centromere function ^7–10^. Complete and highly accurate telomere-to-telomere (T2T) genome assemblies now enable detailed studies of the genetic and epigenetic organization of centromeres ^11–13^. Prior studies in the T2T-CHM13 genome revealed that CENP-A chromatin coincides with centromere dip regions (CDRs), or localized hypomethylated domains within otherwise heavily methylated alpha-satellite (αSat) higher-order repeat (HOR) arrays ^14–18^. CDRs show internal structure composed of multiple discrete hypomethylated “sub-CDRs” separated by methylated intervals, and they tend to localize to the youngest, most homogeneous repeats within each αSat array ^15,19^. Complementing these assembly-based analyses, Directed Methylation with Long-read sequencing (DiMeLo-seq), an antibody-directed protein–DNA interaction mapping approach combined with long-read sequencing, has enabled initial maps of CENP-A enrichment within satellite DNA ^20,21^. While these early findings established the association between DNA methylation and CENP-A positioning, fundamental questions remain: What determines CENP-A subdomain boundaries? How does this multi-domain architecture vary between homologous chromosomes in diploid genomes, and how does it vary between different cell types from the same donor?

The HG002 genome provides a strong foundation for beginning to answer these questions, by combining a complete, high-accuracy diploid assembly ^22^ with a well-characterized and widely available biological resource. HG002 is a Genome in a Bottle benchmark sample with a stable diploid XY karyotype, extensive publicly available sequencing data, and broad donor consent for research use ^23–25^. Multiple matched cell resources are available, including the reference lymphoblastoid cell line (LCL), induced pluripotent stem cell (iPSC) lines derived from lymphoblastoid and peripheral blood cells, and parental cell lines, enabling integrated haplotype-resolved and cross–cell type analyses.

Here, we pair detailed centromere satellite characterization within the fully phased diploid T2T assembly ^22^ with ultra-long-read, single-molecule chromatin profiling to define centromeric chromatin organization. To understand the underlying centromere satellite sequence, we generated haplotype-resolved satellite annotations and multi-scale maps of centromeric repeat structure, revealing extensive haplotype-specific variation in array size, internal repeat unit organization, and evolutionary layering of related satellite repeats within arrays. We then applied ultra-long-read DiMeLo-seq profiling of CENP-A and H3K9me3, a histone H3 lysine 9 trimethylation mark associated with pericentromeric heterochromatin ^26,27^, to quantify centromeric chromatin architecture at single-molecule resolution. We show that CENP-A domains form multiple ordered subdomains whose aggregate size and protein dosage are broadly constrained across centromeres and between homologs. Finally, by comparing HG002 iPSCs and extended passaged HG002 LCLs, we show that changes in DNA methylation levels are accompanied by remodeling of subdomain boundaries and CENP-A domain spans, establishing methylation as a likely regulator of core centromeric chromatin architecture. Together, these results provide a diploid, single-molecule framework for studying centromere organization and its epigenetic plasticity across human genomes and cell states.

## Results

### Haplotype-resolved annotation reveals multi-scale variation in human centromeres

The complete and highly accurate phased haplotype assemblies in the T2T-HG002 diploid human genome benchmark^22^ enable comparative analysis of centromeric and pericentromeric regions across chromosomes and haplotypes. To support genetic and epigenetic studies of human centromeres, we implemented an automated tool suite to predict satellite DNA annotations across the diploid HG002 assembly (**Methods**). Building on prior expert-curated satellite annotation tracks developed for the T2T CHM13 centromere assemblies, we applied an automated method to generate centromeric satellite (‘censat’) annotations in the diploid HG002 genome, enabling genome-wide classification of satellite families and systematic definition of array organization (**Fig 1a**, **SFig 1**, **STable 1, Methods**).

**Figure 1.**
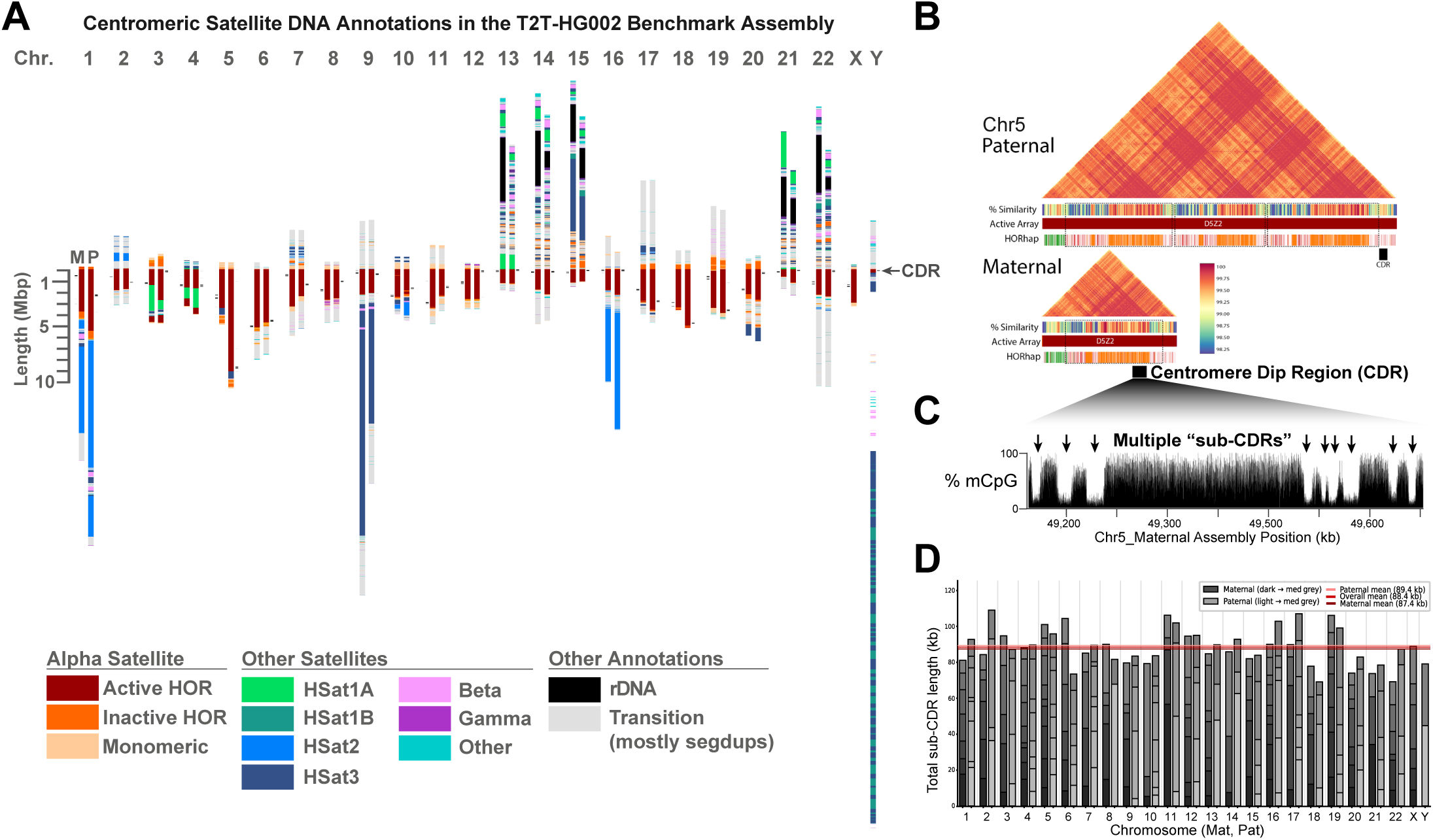
Haplotype-resolved centromere annotation reveals multi-scale variation in the diploid HG002 genome. (A) Centromeric satellite (censat) annotations for maternal (’M’) and paternal (’P’) chromosomes. Annotations are centered at the p-arm proximal boundary of the active array. Acrocentric short-arms (13p, 14p, 15p, 21p, and 22p) are included due to their enrichment of satellite classes common to centromeric regions. The locations of centromere dip regions (CDRs) are indicated in black. (B) Chromosome 5 paternal and maternal haplotypes illustrating the largest homolog-specific size difference observed among active αSat arrays (9.49 Mb paternal vs. 3.59 Mb maternal), with higher-order repeat haplotypes (HORhap) and a 1D modplot heatmap^28^ and 1D projected ModDotPlot tracks. Repeat annotations highlight a triplication in the chromosome 5 paternal active array. CDRs are shown in black within the q-arm proximal region. (C) CpG methylation track across the chromosome 5 maternal active αSat array, illustrating nine discrete sub-CDR hypomethylation dips. (D) Stacked bar plot of sub-CDR lengths for all chromosomes, ordered by genomic position, with mean aggregate sub-CDR lengths indicated separately for maternal (dark red), paternal (light red), and all chromosomes combined (red). Despite extensive variation in active array length, total, or aggregate sub-CDR lengths per chromosome appears similar between homologs and consistent between chromosomes genome-wide (mean ∼90 kb, range 69–109 kb).

The diploid HG002 assembly contains ∼336 Mb of annotated satellite sequence, with alpha satellite (αSat, ∼174 Mb) and classical human satellites 1, 2, and 3 (HSat1-3, ∼138 Mb) constituting the largest fractions, and with smaller contributions from beta satellite (∼13 Mb) and other centromeric satellite subclasses. Diploid, haplotype-resolved satellite annotation enables direct comparison of maternal and paternal chromosomes^13^ and reveals widespread megabase-scale differences in centromeric and pericentromeric arrays, with at least 19 autosomal loci exhibiting haplotype-specific size differences greater than 0.5 Mb (**STable 2**). The largest haplotype-specific differences occur in pericentromeric HSat1-3 arrays and in active αSat arrays, with the most extreme example at the chromosome 9 HSat3 locus, which differs by 10.1 Mb between haplotypes (21.07 Mb maternal vs 11.00 Mb paternal) and contains large inversions and other structural rearrangements (**SFig 2**). HSat2 on chromosome 1 differs by 6.74 Mb between homologs and contains distinct sequence compositions (**Methods, STable 3**), while the paternal haplotype contains an additional ∼500 kb polymorphic block of HSat3 with adjacent beta-satellite sequence (**SFig 3**). Interspersed HSat1A arrays are observed embedded within active αSat arrays on chromosomes 3 and 4, differing in size by ∼0.61 Mb on chr3 and ∼1.1 Mb on chr4, indicating haplotype-specific restructuring and mixed satellite composition within kinetochore-associated centromeric domains (**SFig 4**). Finally, the αSat HOR array on chromosome 5 shows the largest homolog-specific size difference among active arrays, with a total length of 9.49 Mb on the paternal haplotype and only 3.59 Mb on the maternal haplotype, likely resulting from a large triplication event within the array (**Fig 1b**). Based on more than 200 currently available long-read assemblies, the 9.49 Mb paternal chromosome 5 array appears to be the largest αSat array reported in a human genome assembly to date ^11–13,19^.

Differences between homologous centromeres are not limited to total array length but also extend to internal repeat organization and evolutionary structure within αSat arrays (**SFig 5**). αSat DNA is organized into multimeric higher-order repeats (HORs) that form homogenized arrays associated with kinetochore assembly and centromere function. Using HOR-level annotation together with structural variant (StV) classification to detect monomer-scale insertions and deletions within canonical HOR units, we observed broad variation in HOR structural composition across HG002 centromeres (**SFig 6, STable 4**). Some active arrays are structurally uniform, including chromosomes 11, 14, and 22 on both haplotypes, along with chrX, where a single structural HOR variant accounts for at least 95% of units. In contrast, other centromeres show high structural diversity, most prominently chr19 on both haplotypes, followed by chromosomes 1, 4, 5, and 10, each containing numerous distinct HOR structural variants at appreciable frequency.

Homologous centromeres can also differ markedly in variant composition even when total array sizes are similar. For example, the chromosome 13 active arrays have comparable lengths but highly distinct variant profiles (**SFig 7**). To further resolve this structure, we grouped HOR units sharing characteristic sequence variants into haplotype classes, termed HORhaps (**Methods, SNote 1**) and mapped their distribution across active arrays. HORhap composition often differs substantially between homologs, again illustrated by chr13, where profiles indicate near-complete haplotype turnover with only a small residual paternal haplotype tract at the edge of the maternal array (**SFig 7**). Using these HORhap patterns, we also created maps of superHORs, which are defined by recurring HOR combinations, which occupy large fractions of several arrays, including chromosomes 8 (∼0.46 paternal, ∼0.51 maternal), 15 (∼0.43 paternal, ∼0.40 maternal), and 9 (∼0.33 paternal, ∼0.38 maternal) (**Methods**; **SNote 1**).

To link the structural and evolutionary variation described above with functional centromere identity, we next analyzed CDRs, the localized hypomethylated domains within αSat HOR arrays that mark sites of CENP-A chromatin assembly (**Fig 1a,c, STable 5**). Using genome-wide methylation profiles, we identified the span of each CDR and defined sub-CDR domains across active centromeres on both homologs (**Methods**; **Fig 1a,c**). When classified by array position, in agreement with prior studies ^29^, we find approximately half of CDRs localized to the p-arm–proximal portion of the active array (41.3%), whereas fewer occurred near the q-arm (28.2%) (**SFig 8**). Consistent with prior observations in T2T-CHM13 ^15^ and recent observations across many assemblies ^19^, we observe that CDRs preferentially occur within more sequence-homogeneous αSat subregions (Quantitative comparison of local HOR similarity; permutation test, N = 100,000, p < 0.0001, **SFig 9**). Next we asked whether CDRs preferentially overlapped younger or older HORhap domains. Although CDRs more frequently overlapped younger HORhaps (28 vs. 9 chromosomes), this trend was not statistically significant after accounting for their relative genomic proportions (binomial test, p = 0.092; see Methods, STable 5, **SNote1**).

Although the total span from the first to last sub-CDR varies across chromosomes and between maternal and paternal homologs and dramatically across the genome (**SFig 10**), the summed length of sub-CDR segments is comparatively stable genome-wide, averaging ∼88 kb (range: 69 to 109 kb, **Fig 1d**), with no apparent relationship to the total active HOR array size (**SFig 11**). Despite extensive haplotype-specific differences in array size, structural composition, and inferred expansion age, the total hypomethylated core domain is remarkably constrained in length. Together, these results show that hypomethylated centromere cores can be reproducibly identified in diploid T2T assemblies, are associated with locally homogeneous αSat sequence, and maintain a shared functional sub-CDR aggregate length despite the highly variable sizes of the associated active HOR arrays.

### Single-molecule profiling reveals CENP-A domain organization and dosage at centromeres

Having established haplotype-resolved centromeric sequence organization and well-defined CDRs, we next examined variation in centromeric chromatin structure across homologs. CENP-A–containing nucleosomes mark sites of kinetochore assembly, yet their precise relationship to centromere dip regions (CDRs) remains unclear, including how consistently they colocalize with CDRs and whether CENP-A occupancy spans the full CDR or only subregions across a population of cells. To address this, we generated a high-coverage, ultra-long-read epigenomic dataset to enable detailed mapping of centromeric chromatin in HG002 LCLs. To do this, we performed DiMeLo-seq ^20^, which combines antibody-targeted methylation with nanopore sequencing (**Methods**), to directly map CENP-A–DNA interactions across HG002 active arrays. Because CDR spans typically extend ∼100–400 kb based on our HG002 analysis, high-coverage ultra-long nanopore reads are required to phase signals to maternal and paternal haplotypes, span the CDR, and measure CENP-A domain architecture on single DNA molecules.

Achieving the resolution needed to quantify epigenetic variation across CDRs and active arrays requires deep coverage; however, CENP-A–associated regions constitute only a small fraction of the genome. Existing restriction-based centromere enrichment approaches require substantial input material and introduce handling steps that reduce achievable read length ^20,30^. To overcome these constraints, we implemented nanopore adaptive sampling (with 100,000 bases flanking each active arrays, **STable 6**) after ultra-long library preparation, using a depletion strategy that selectively rejects non–αSat molecules in real time based on alignment of the initial few hundred bp of each read ^31^, thereby enriching centromeric arrays while preserving ultra-long read lengths (N50 63 kb; with ∼20 x coverage 100kb+ reads; **Methods, Fig 2a**). This strategy dramatically improved coverage of centromeric regions, as illustrated by the chrX αSat HOR array, where adaptive sampling achieves ∼54 x coverage, which is about 30 times higher than the rest of the genome (**Fig 2a**). Overall, compared to standard sequencing without adaptive sampling, this approach achieved approximately 4-fold enriched coverage of centromeric regions per sequencing flow cell without additional sample preparation (**Fig 2a**). Coverage is consistent across chromosomes and across each HOR array (**SFig 12**). Overall, this strategy yields a uniquely high-resolution, haplotype-resolved, centromere-targeted epigenomic dataset, establishing a foundation for systematic analysis of CENP-A chromatin across fully assembled human centromeres.

**Figure 2.**
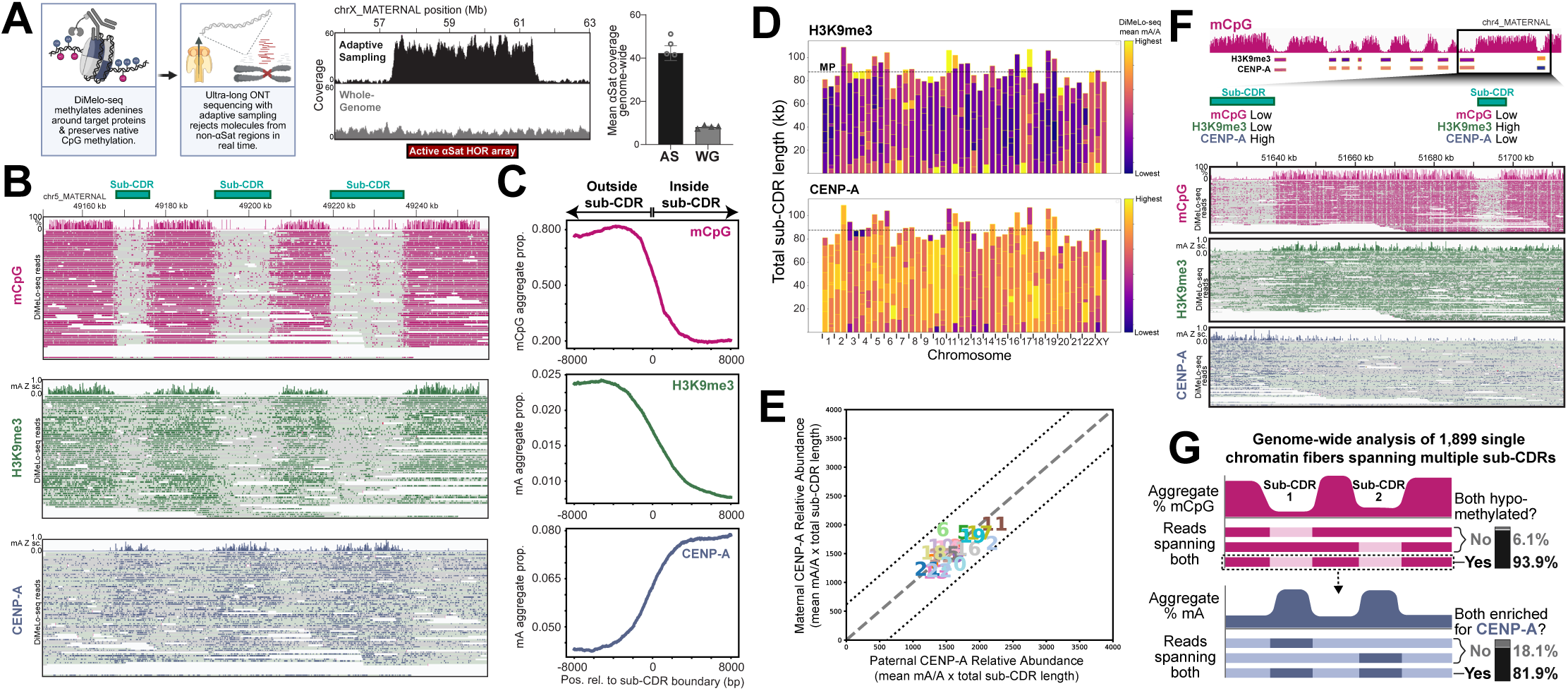
Single-molecule profiling of CENP-A chromatin architecture across haplotype-resolved human centromeres. (A) Schematic ^33^ of the adaptive sampling DiMeLo-seq enrichment strategy and data processing pipeline. Coverage comparisons between adaptive sampling and whole genome sequencing are shown for the chromosome X αSat HOR array and genome-wide, illustrating ∼30-fold enrichment of centromeric regions relative to the rest of the genome. (B) Integrative Genomics Viewer (IGV) ^34^ visualization of a region spanning a CDR on chromosome 5 (chr5_MATERNAL: 49150874-49257508) within the active αSat array. mCpG (magenta), H3K9me3 (green), and CENP-A (blue) DiMeLo-seq signal tracks are shown, illustrating the relationship between CENP-A enrichment and the heterochromatic (H3K9me3 and mCpG) boundaries of sub-CDR intervals (indicated by black bars). (C) Aggregate profile plots of mCpG, H3K9me3, and CENP-A signal centered on CDR boundaries across all centromeres, demonstrating an inverse relationship between CpG hypomethylation and CENP-A enrichment relative to flanking H3K9me3-marked heterochromatin. (D) Stacked heatmaps of H3K9me3 (top) and CENP-A (bottom) DiMeLo-seq 6mA signal density across all aggregated sub-CDRs, ordered by chromosomal position and separated by maternal and paternal haplotype. The dotted line indicates the mean length (∼90 kb) of aggregate sub-CDRs. The majority of sub-CDRs display an inverse relationship between CENP-A and H3K9me3 (where CENP-A signal is high, H3K9me3 signal is low), with variation observed primarily in the outermost sub-CDRs. (E) Scatter plot comparing DiMeLo-seq CENP-A 6mA relative abundance between maternal and paternal homologs across all centromeres, showing broadly balanced signal between haplotypes. (F) IGV visualization of an atypical sub-CDR at chromosome 4 (chr4_MATERNAL:51622694-51707732), where CENP-A enrichment is absent and H3K9me3 is present within the sub-CDR, representing a departure from the canonical centromeric chromatin architecture in which H3K9me3 and mCpG are excluded from sub-CDRs. (G) Schematic ^35^ illustrating co-occupancy of CENP-A across multiple sub-CDRs on individual ultra-long nanopore reads, as assessed using a decision-tree classifier trained on single-molecule signals from CDR and non-CDR regions. The majority of reads spanning two sub-CDRs exhibit CENP-A enrichment at both sub-CDRs.

Alignment of DiMeLo-seq reads to the diploid HG002 assembly enabled haplotype-resolved chromatin profiling across active centromere arrays. Mappability analyses confirmed that reads at these lengths align confidently and with haplotype specificity to active centromere arrays (**SNote 2**). Individual molecules showed multiple CENP-A–enriched segments that coincided with annotated sub-CDRs, and boundary-centered analyses (**Methods**) across all centromeres revealed a consistent inverse relationship between CpG methylation and CENP-A enrichment, with CENP-A signal increasing as CpG methylation decreased (**Fig 2b,c**).

To define the flanking heterochromatin context, we also generated ultra-long DiMeLo-seq datasets targeting H3K9me3, a hallmark of pericentromeric heterochromatin that typically co-occurs with CpG hypermethylation. Across centromeres, H3K9me3 enrichment closely tracked CpG methylation and showed coordinated depletion at sub-CDR intervals, forming reciprocal patterns relative to CENP-A signal and sharply delineating centromere core versus pericentromeric domains (**Fig 2c**, consistent with other recent work ^32^). The concordant CpG and H3K9me3 profiles, together with their exclusion from CENP-A–enriched regions, reveal a compartmentalized chromatin architecture at centromeres.

We observed that sub-CDR boundaries sometimes displayed mixed methylation states, in which only a subset of reads retained high 5mC while others were hypomethylated, consistent with plasticity in exact sub-CDR boundaries across the population of cells. Clustering based on CpG signal separated these reads into multiple 5mC-defined groups (Methods), supporting the interpretation that these patterns reflect genuine molecule-level epigenetic heterogeneity rather than measurement noise (**SFig 13**).

To further resolve chromatin footprints at single-molecule resolution, we generated and analyzed nanopore ultra-long Fiber-seq ^36^ data for HG002 LCLs (**Methods**). Consistent with prior observations ^37^, we observed smaller nucleosome footprints within sub-CDRs in the active arrays (versus non-active arrays and outside of sub-CDRs), with positioning adjacent to canonical CENP-B box motifs (**SFig 14**). Notably, the reduced footprint size is consistent with prior reports that CENP-A–containing nucleosomes adopt a more compact structure than canonical H3 nucleosomes ^38^, supporting the interpretation that these protected regions reflect centromere-specific chromatin architecture.

Given that the aggregate length of sub-CDR intervals is similar across centromeres and between homologs, we next tested whether CENP-A signal is also quantitatively balanced across these domains. We estimated CENP-A dosage using DiMeLo-seq 6mA signal density, defined as the number of detected mA bases divided by the total number of adenine positions on all reads within each interval (**Methods**). CENP-A density is largely consistent across sub-CDRs and shows similar levels between homologous centromeres, matching the constrained total span of aggregated sub-CDR domains (**Fig 2d**). These measurements indicate that CENP-A chromatin dosage is broadly balanced across human centromeres (**Fig 2e**), consistent with prior GFP-based estimates of genome-wide CENP-A abundance ^39^. In further support of this observation, we observe CENP-C DiMeLo-seq mA density to also be quantitatively balanced across chromosomes and between homologous centromeres (**SFig 15**). Notably, the greatest variability occurs in smaller, outer sub-CDRs, where reduced CENP-A density is accompanied by increased H3K9me3 signal (**Fig 2d**). These intervals mark localized departures from the typical reciprocal pattern among CENP-A, CpG hypomethylation, and H3K9me3, including sites where sub-CDR methylation signatures are present without corresponding CENP-A enrichment and, conversely, where heterochromatin marking extends into hypomethylated sub-CDR regions (**Fig 2f**). Overall, these dosage and boundary patterns indicate that CENP-A is broadly balanced across centromeres and between homologs, while limited edge variability is detectable at a fine scale.

We next leveraged the unique ultra-long, single-molecule nature of our DiMeLo-seq data to test whether multiple sub-CDR domains are co-occupied by CENP-A on the same chromatin fibers (as illustrated, **Fig 2g**), or whether they arise from averaging across heterogeneous cell populations. Single-molecule analysis supports the former. Of 247,716 ultra-long reads overlapping at least one sub-CDR, 1,899 fully span two or more sub-CDRs. Among these spanning reads, 93.88% show mCpG depletion across all annotated sub-CDR intervals, consistent with the predicted spacing and organization of sub-CDRs, with only a small fraction indicating variation in sub-CDR structure. CENP-A enrichment patterns show similar concordance, with 81.85% of reads spanning multiple sub-CDRs exhibiting CENP-A signal at more than one sub-CDR on the same molecule, whereas 18.15% lack signal at one or more spanned sub-CDRs (**Fig 2g**). Together, these single-molecule sequencing results indicate that the majority of centromeres in the sampled cell population contain multiple CENP-A–associated subdomains, while also revealing measurable cell-to-cell variation in sub-CDR occupancy by CENP-A.

### DNA methylation state controls sub-CDR boundaries and CENP-A domain size

Under typical conditions in HG002 lymphoblastoid cells, centromeric chromatin is organized into distinct and functionally specialized domains. In this organization, discrete islands of CENP-A chromatin are interspersed and largely mutually exclusive with H3K9me3-enriched, CpG methylated pericentromeric heterochromatin. However, it remains poorly understood whether this fine-scale epigenetic architecture is robust to larger epigenetic changes across different cell types and cell states. To begin to test this, we passaged HG002 LCLs for an additional 25 days (which we refer to as “late-passage” cells), as extended culture of transformed LCLs has been associated with CpG methylation changes ^40^, which may in turn affect centromere organization. We generated ONT ultra-long DiMeLo-seq profiles for CENP-A and H3K9me3 in these late-passage cells (**Methods**) and compared them to our initial (or “early-passage”) sub-CDR analyses. We found that methylated intervals separating neighboring sub-CDRs were diminished (**Fig 3a**), and the H3K9me3 signal was reduced at inter-sub-CDR boundaries, resulting in more frequent formation of continuous CENP-A–occupied regions (**Fig 3a,b**). Sub-CDR expansion and consolidation was reported genome-wide, with a 16.9% reduction in sub-CDR number (254 to 211) accompanied by a 444 kb increase in total CDR length (increasing by 10.9%) (**STable 7**, **SFig 17**). This reorganization was primarily driven by boundary resizing rather than wholesale domain gain or loss: 97.2% of early-passage sub-CDRs (247 of 254) were maintained in late-passage cells, with only 7 sub-CDRs lost entirely and 4 novel sub-CDRs appearing. Of the maintained sub-CDRs, 184 (74.5%) expanded while 63 contracted, further supporting that boundary shifts rather than de novo domain turnover are the principal mode of reorganization. Sub-CDRs were correspondingly larger, with mean length increasing from 16 kb to 21.4 kb and maximum sub-CDR size expanding to ∼100 kb in late-passage cells compared to 68 kb in early passage. This reorganization was widespread, with 17 of 24 chromosomes showing reduced sub-CDR counts and most chromosomes exhibiting increased total CDR span, although several centromeres showed minimal change (<10%), including chromosomes 3, 17, and Y, indicating chromosome-specific responses. Further, sub-CDR remodeling was positionally biased; that is, centrally located sub-CDRs preferentially expanded, whereas sub-CDRs flanking CDR boundaries more often gained H3K9me3 signal in late-passage cells (**Fig 3c**). CpG methylation levels did not differ between central and outer sub-CDRs, indicating that boundary-proximal CENP-A loss and heterochromatin gain are not explained by local CpG differences (**SFig 18**). Collectively, these results show that methylation loss is associated with boundary erosion and positionally structured sub-CDR remodeling, leading to consolidation of CENP-A chromatin into fewer, larger centromeric domains.

**Figure 3.**
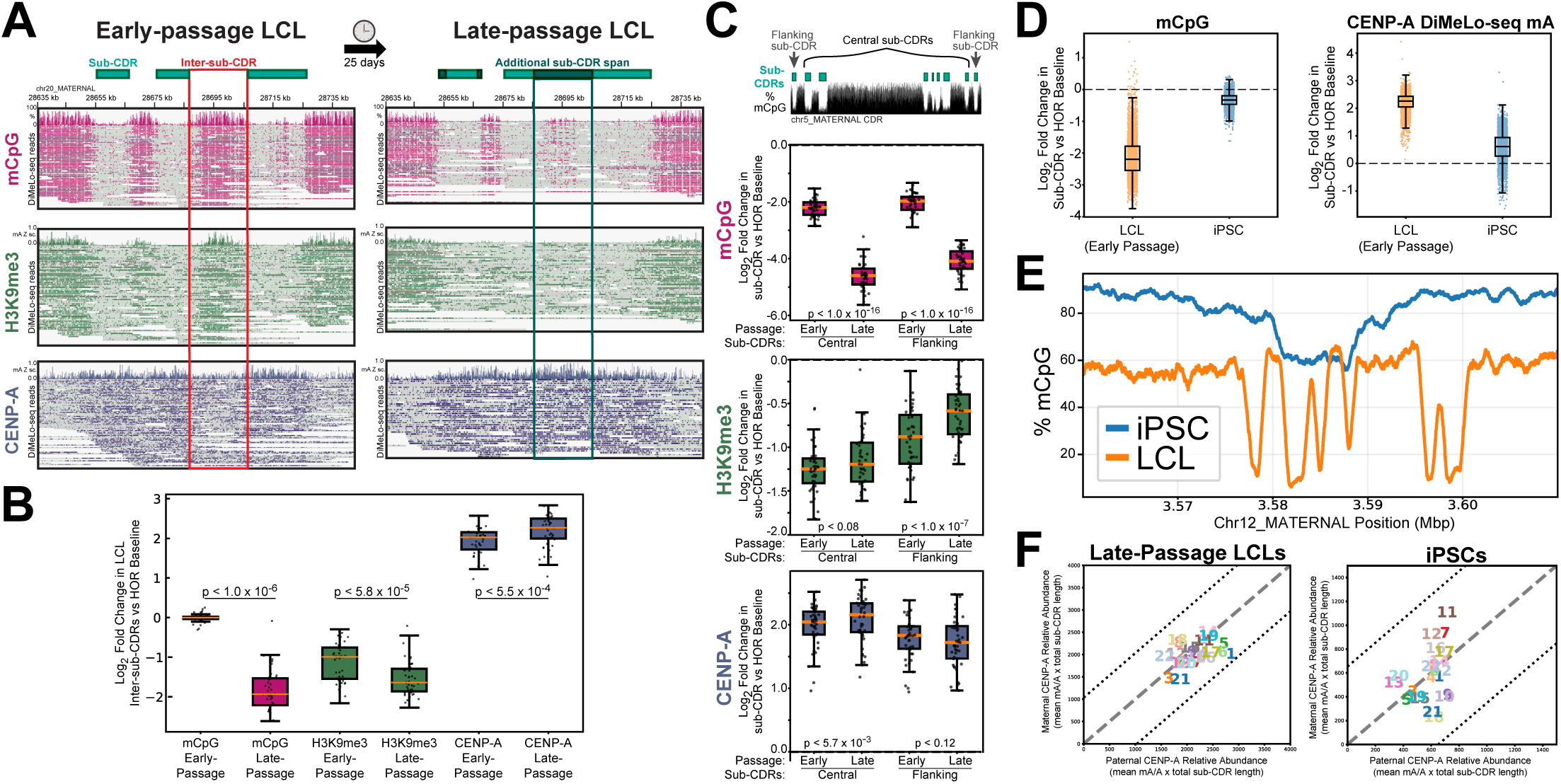
DNA methylation state influences sub-CDR boundaries and CENP-A domain organization. (A) IGV visualization of the chromosome 20 (chr20_MATERNAL:28635505-28746668) CDR region showing mCpG (magenta), H3K9me3 (green), and CENP-A (blue) DiMeLo-seq signal tracks in early- and late-passage (25 days) HG002 LCLs. Sub-CDRs are highlighted in light green. Sites where methylated intervals separating neighboring sub-CDRs were diminished are indicated by dark green bars and a red box, resulting in more frequent formation of continuous CENP-A–occupied regions. (B) Log2 fold-change of mCpG, H3K9me3, and CENP-A DiMeLo-seq signal densities at inter-sub-CDR boundaries between early- and late-passage LCLs. Within regions of boundary loss, there is a significant decrease in mCpG between early and late passage, a concurrent decrease in H3K9me3 and a corresponding increase in CENP-A. (C) Illustration of central and flanking sub-CDRs in the chr5_MATERNAL CDR region. Genome-wide fold-change plots show CENP-A density increasing in central sub-CDRs but not in flanking sub-CDRs in late-passage LCLs, with H3K9me3 showing the reciprocal pattern and mCpG remaining largely unchanged across sub-CDR positions. (D) Fold-change analysis of mCpG and CENP-A signal densities across 1 kb bins within sub-CDRs in HG002 early-passage LCLs and iPSCs, with non-CDR active array regions used as a baseline. HG002 hiPSCs show a reduction in CENP-A enrichment and an increase in CpG methylation within the active array. (E) Line plot comparing sub-CDR positions in hiPSCs and LCLs for chr12_MATERNAL. The hiPSC active array shows elevated CpG methylation in the active array (∼90%) and within sub-CDRs (∼60%) relative to LCLs (∼60% and ∼20%, respectively). Although the CDR in hiPSCs is broader than in LCLs, it occupies only a partial span of the corresponding LCL CDR. (F) Scatter plots of relative CENP-A abundance, estimated by multiplying mean 6mA/A density by total or aggregate sub-CDR(s) length, for maternal and paternal homologs in late-passage LCLs and iPSCs. Despite extensive sub-CDR reorganization, relative CENP-A abundance remains proportional across chromosomes and balanced between homologs.

Having observed that methylation loss is accompanied by sub-CDR boundary erosion and CENP-A domain expansion, we next asked whether, conversely, elevated DNA methylation levels associate with reciprocal changes in centromeric domain organization. To test this, we analyzed CENP-A and sub-CDR architecture in donor-matched HG002 induced pluripotent stem cells (iPSCs, with quality assessment in **SNote 3, 4**), which show increased CpG methylation across active centromeric arrays relative to early-passage LCLs (**Fig 3d, SNote 4**). Whereas LCL arrays typically exhibit ∼60% CpG methylation with pronounced CDR dips below 20%, iPSC arrays are more uniformly methylated (∼90%) and display only shallow CDR depressions (∼60%) (**Fig 3e**). Under these hypermethylated conditions, CDR organization was extensively reorganized (**SFig 19, STable 8)**. Total sub-CDR number decreased approximately two-fold (254 to 118), while mean CDR length increased 2.5-fold (16.0 kb to 40 kb), producing a strong shift toward large continuous domains with maximum sizes reaching ∼168 kb. Despite this, total CDR coverage increased (4.1 Mb early-passage LCL to 4.8 Mb in hiPSCs), indicating genome-wide merging of adjacent sub-CDRs. The majority of centromeres showed increased total CDR span despite fewer domains, with the largest gains on chromosomes 7, 11, 14, and 17 (**SFig 19**). Notably, sequence similarity analysis (1D ModDotPlot similarity data) indicated that these methylation-associated domain expansions are likely not driven by underlying sequence identity, as regions incorporated into expanded iPSC domains are indistinguishable from former LCL inter-sub-CDR gaps, and no consistent directional shift in similarity was observed across chromosomes.

We next examined whether the reorganization of sub-CDRs was accompanied by changes in centromeric chromatin composition, and we found that CENP-A density also decreased significantly relative to LCLs (**Fig 3d**). However, although overall CENP-A density is reduced, this reduction appears mostly uniform across chromosomes and between homologs, with average sub-CDR DiMeLo-seq 6mA signals staying within 2-fold (with two notable exceptions including small sub-CDRs at a distance with weak support, **SNote 4**, **SFig 20**, **Fig 3f**). While most homolog pairs show consistent 6mA signal density and aggregate sub-CDR lengths, a small subset of chromosomes display inter-homolog asymmetry (**Fig 3f**, **SFig 20**), which may reflect biological variability or technical limitations in defining sub-CDR boundaries within highly methylated active arrays. Notably, similar to early-passage lines (**Fig 2e**), we observe a preservation of relative CENP-A DiMeLo-seq 6mA signal density under hypomethylated conditions in late-passage LCLs: although sub-CDRs expanded and boundaries eroded, homolog balance was roughly maintained (**Fig 3f, SFig 21**). Whether this plasticity in centromeric architecture is a general feature–or whether it is amplified in pluripotent cells, where hypomethylation is pervasive and epigenetic landscapes are broadly reshaped–remains an important open question. Together, these findings support a model in which centromeric chromatin exhibits methylation-sensitive architectural plasticity, where shifts in DNA methylation levels reshape sub-CDR organization.

## Discussion

Our work has provided new insights into the organization and heterogeneity of human centromeres enabled by the HG002 benchmark diploid assembly in combination with long-read, single-molecule epigenomic technologies. By building and deploying automated satellite DNA annotation tools, we uncovered substantial variation in satellite DNA array size, structure, and sequence composition when comparing the maternal and paternal haplotypes of HG002. Furthermore, we optimized ONT’s adaptive sampling method to enable enriched coverage of centromeric regions without any additional sample preparation steps ^20^, overcoming challenges due to the repetitiveness and high polymorphism of these regions. By combining this approach with ultra-long nanopore sequencing, we obtained single-molecule reads that could span across CDRs, which are clusters of hypomethylated sub-CDR regions that form the core of human centromeres. Using DiMeLo-seq, we could use these long, centromere-enriched reads to examine the localization of CENP-A and H3K9me3 at fine scales, while simultaneously measuring endogenous mCpG on the same single chromatin fibers.

These unique data enabled us to measure several fundamental properties of human centromeres. We observed that centromeres in HG002 LCLs contain on average ∼5.5 hypomethylated sub-CDR regions (range: 2-9), which on average have a summed length of 88 kb (range: 69–109 kb), irrespective of the number of sub-CDRs or the total length of the active αSat array on that chromosome. From DiMeLo-seq data, we also found that the mean intensity of CENP-A-targeted DiMeLo-seq signal was relatively constant across haplotypes but with mild variation between chromosomes. Together, these data suggest that the span of centromeric chromatin is mostly uniform across chromosomes despite high variation in αSat array size, with little variability in the dose of CENP-A per chromosome, consistent with prior work from quantitative imaging of CENP-A ^39^ and immunofluorescence-FISH assays ^41^. However, these slight changes in CENP-A dosage could still directly impact centromere function, as previously observed ^41,42^.

We observed at fine scales on single chromatin fibers that CENP-A localization is strongly anti-correlated with mCpG, while H3K9me3 is strongly correlated with mCpG, consistent with other recent work using DiMeLo-seq ^19,21,27,32,43^. However, our single-molecule data allow us to observe important exceptions to these trends. For example, we observed several hypomethylated regions that lack CENP-A and are enriched for H3K9me3. Furthermore, given our ultralong sequencing reads, we observed that distinct sub-CDRs tend to be co-occupied by CENP-A on the same chromatin fibers. This allows us to rule out a model in which CENP-A heterogeneously occupies one of several sub-CDRs in each cell. Thus, on single chromatin fibers, CENP-A is highly enriched within multiple neighboring spans of hypomethylated DNA separated by hypermethylated DNA enriched for H3K9me3. Perhaps these separate domains of CENP-A along single chromatin fibers can explain recent observations of multi-partite mitotic centromere structures using super-resolution microscopy ^6^.

Intriguingly, we also observed that the length and number of individual sub-CDRs vary across chromosomes and correlate with changes in array methylation due to cell passaging or reprogramming. This raises the possibility that shifts in sub-CDR organization, and in CENP-A balance, may influence the capacity to form and faithfully inherit multi-partite centromere architectures, potentially even changing the number of kinetochores assembled on each chromosome. Understanding what drives this variability, and why certain sub-CDRs on particular chromosomes are more susceptible to re-organization than others, remain open questions. An important and unanswered question is whether shifts in sub-CDR organization occur naturally over time within an individual, for example, during aging, tissue differentiation, or in response to environmental epigenetic perturbations. If so, centromeric chromatin remodeling may represent an underreported somatic epigenetic variant with potential consequences for long-term chromosome segregation fidelity. We recently demonstrated in separate work that specifically perturbing DNA methylation at centromeres causally alters CENP-A localization and centromere function ^21^. Others have recently shown that perturbing pericentric histone modifications can lead to loss of centromeric chromatin confinement ^27,32,43^, and that fine-scale sub-CDR organization can undergo dramatic shifts in cancer cells ^9^ and sperm cells ^44^.

Beyond DNA methylation, the molecular mechanisms that establish and maintain sub-CDR boundaries and spacing remain poorly understood. Higher-order chromatin features, such as centromere-specific looping architectures, phase-separated chromatin compartments, or other secondary structures not yet well characterized in this context, may play important roles in organizing the multi-partite centromere core, and represent compelling targets for future investigation. Previous work has shown by microscopy that cellular reprogramming induces depletion of CENP-A at centromeres, accompanied by increases in segregation errors^45^. Our DiMeLo-seq results in HG002 iPSCs confirm this loss of CENP-A with an orthogonal genomic approach and further demonstrate that this loss of CENP-A is accompanied by DNA hypermethylation and restructuring of the fine-scale organization of centromeric chromatin. Together these insights suggest that the mechanisms governing centromeric architecture and function are complex and still not fully resolved, presenting an exciting opportunity for future exploration. The rich sequence annotation and scalable tools presented here, when combined with DiMeLo-seq across diverse cell types and additional genome assemblies, will provide a foundation for future investigation of the genetic and epigenetic influences on sub-CDR organization and its inheritance.

## Resource Availability

The lymphoblastoid cell line (LCL) GM24385, induced pluripotent stem cell (hiPSC) line GM26105 (reprogrammed from GM24385), and hiPSC line GM27730 (reprogrammed from peripheral blood mononuclear cells, PBMCs) were obtained from the NIGMS Human Genetic Cell Repository at the Coriell Institute for Medical Research.

## Lead Contact

Requests for further information and resources should be directed to and will be fulfilled by the lead contacts, Nicolas Altemose or Karen H. Miga.

## Code Availability

The code to reproduce the results in this manuscript is available on https://github.com/yxu405/HG002_epi_paper

## Declaration of Interests

NA is named as an inventor on patent applications related to DiMeLo-seq.

## Declaration of Generative AI and AI-assisted Technologies in the Writing Process

During the preparation of this work, the authors used LLMs in order to assist with specific programming tasks, and on rare occasions to improve the language of the manuscript. After using these services, the authors reviewed and edited the content as needed and take full responsibility for the content of the published article.

## Acknowledgements

Thank you to Daniele Fachinetti and other colleagues who provided helpful comments on this study.

## Funding

Grants supporting this work are from the US National Institutes of Health, awarded to KHM NIH/NHGRI R01 1R01HG011274, NIH/NHGRI UM1 HG010971. HL is supported by NIH training grant T32HG123442. DD is supported by NIH training grant T32GM141828. FR was supported by the HSE basic research program, MC is supported by MUNI Award in Science and Humanities StG/CoG (MUNI/SC/1916/2024). TAP is supported by R50CA305001 and SIMR. JLG is supported by R01CA266339 and SIMR. NA is a HHMI Hanna H. Gray Fellow, Pew Biomedical Scholar, and Biohub Investigator. KHM was supported in part by the Searle Scholars Program.

## Author Contributions

KHM, NA, and YX conceived of the study. YX, JMVG, and BM performed experiments. YX, HL, JM, FR, JKL, MC, LM, EX, DD, MA, IV, CO, TH, SOR, WT, and IAA analyzed the data. CC, TAP, JLG, JPS, AFS, and MWM provided key reagents and samples. KHM, NA, and YX wrote the paper. KHM and NA supervised the study.

## Methods

### EXPERIMENTAL MODEL AND STUDY PARTICIPANT DETAILS

The donor of the GM24385, GM26105 and GM27730 cell lines is male. The cell lines’ species identities are confirmed by Coriell by enzyme isoenzyme electrophoresis (nucleoside phosphorylase, glucose-6-phosphate dehydrogenase, and lactate dehydrogenase), but no additional STR-based authentication was performed in this study; the line was used as distributed by the Coriell Institute.

### Ethics statement

All research involving human induced pluripotent stem cell (hiPSC) lines GM26105 and GM27730 was conducted in compliance with the International Society for Stem Cell Research (ISSCR) 2021 Guidelines for Stem Cell Research and Clinical Translation (Category 1A: routine pluripotent stem cell research). Both cell lines were originally derived by the Coriell Institute for Medical Research under informed consent that permits research use, including differentiation into germ layer derivatives and the generation of stem cell–derived models of early development. No new embryos or gametes were created or used for the purposes of this study. Donors provided informed consent specifying that their donated materials may be used for research on stem cell differentiation and developmental modeling.

### Centromere Satellite Annotation Tool Suite

#### Centromere Satellite (‘censat’) Annotation Track

Centromeric satellites were annotated using the censat workflow. The complete workflow can be accessed via GitHub (https://github.com/kmiga/alphaAnnotation). αSat were annotated using a modified version of a HumAS-HMMER, which searches in a genome for sequences from a database of hidden markov models (HMMs) of HOR profiles ^46^. This database contains models of known HORs and suprachromosomal families (SFs) HumAS-HMMER ^47,48^. These annotations were then sorted, and merged into summary bins including active centromeric array, and pericentromeric inactive HOR, diverged HOR (dHOR), and monomeric alpha regions. Classical satellites (HSATII and HSATII) were annotated using a script created by Altemose et. al. which uses a database of human-specific classical satellite kmers ^49,50^. The final subset of pericentromeric satellites (HSat1A, HSat1B, bSats and gSats, and smaller species such as SST1, SATR, and ACRO) were annotated using RepeatMasker^48^ Ribosomal arrays were annotated using hidden markov models based on the first and last 700 bp of the rDNA repeat unit (GenBank accession U13369.1) ^51^, as well as two of the rDNA genes; 18S (NCBI Reference Sequence XR_007084227.1) and 5.8S (NCBI Reference Sequence XR_007084259.1) ^52^. Annotations were then compiled to create a complete annotation of centromere-associated satellites and tandem repeats, including resolving overlaps, filtering out small satellite arrays (<2kb) and merging adjacent annotations. Finally, centromere transition (CT) regions were identified by their proximity to centromeric satellites using a 2MB bedtools merge extending out from the active αSat array through the regions containing pericentromeric satellites.

#### HORhap methods

To analyze and compare haplotypes present in the αSat higher order repeats, HOR haplotype (horhap) annotation of the HG002 chromosomes, the horhap tool [https://github.com/fedorrik/horhap_tool] was used. This work is described in more detail in **SNote 1**.

For each chromosome, both homologs were processed simultaneously. ASat StV (Structural Variants) annotations^15^ were used to retrieve all HOR copies for two of the most frequent StVs. In cases where the second most frequent StV was less frequent than 1% of the total number of HORs, only the most frequent StV was used. In cases where the third most frequent StV was more frequent than 14% of the total number of HORs, top 3 StVs were used. Chromosome 19 was excluded from the horhap analysis due to complex StV structure. The number of selected StVs is shown in the **STable 9**.

Sequences of selected HORs were aligned using muscle3 ^53^. In cases of aligning three StVs per chromosome, muscle3 tends to align different monomers to each other and place deletions of different monomers in one spot. In order to avoid that, the alignment of three StVs was performed by stages. HORs of each of 3 StVs were aligned separately with one copy of full-length StV using muscle3. Then using a custom python script (<u>merge_alignments.py</u>), the alignments of 3 StVs were merged.

The pairwise Hamming distances between all the aligned sequences were calculated and the hierarchical Ward’s linkage clustering was performed to divide the HORs into k clades at a certain depth. For each pair of chromosomes, bed files and alignment figures for all k from 2 to 9 were generated and after inspection, only one k for each chromosome was selected to obtain a compact view of array architecture. In general, higher k values reveal more details of the array structure seen as a number of more or less uniformly colored domains. However, when k increased too much, no uniformly colored domains remained and all or most regions appeared as mixtures of different HORhaps. We aimed at selecting the k value which allowed “optimal regionalization” (i.e. maximal number of relatively homogeneous regions).

#### HORhap age analysis

In CHM13 centromeres, it has been shown that the kinetochore tends to sit on younger horhap arrays^15^. The horhap age analysis was used to demonstrate whether this statement holds true for HG002. For each horhap, the average divergence between the HORs of a given horhap was calculated and these values were used to estimate the age of the horhaps and mark the horhap tracks accordingly. Horhaps with higher divergence were presumed to have expanded earlier and were colored gray, while horhaps with lower divergence were presumed to have expanded more recently and were colored red. Also phylogenetic trees of horhap consensus sequences were created and phylogenetic relations of horhaps were taken into account to determine which horhap is the youngest. This work is described in more detail in **SNote 1**.

The horhap annotations were intersected with CDRs and the cases of CDRs overlapping younger and older horhaps were counted. 28 chromosomes had CDR overlapping younger horhap, 9 cases had CDR overlapping older horhap and 9 other chromosomes did not have well distinguished horhaps so they were not used in this analysis (**STbl 10**). Apparently, this is in line with kinetochore “preferring” more recently expanded horhaps, which was observed as a weak trend in Altemose 2022. However, when the relative collective sizes of younger/older horhap arrays are taken into account, this trend becomes less reliable. Using the binomial distribution formula, the null hypothesis (kinetochore has no preferences) cannot be rejected (p-value = 0.092).

#### SuperHOR annotation

To analyze if patterns of HOR form yet higher repetitive structure, superHOR annotation was produced using a notebook (https://github.com/fedorrik/superHOR_HG002). The script takes horhap annotation and searches for n-mers (n is 2 – 9) which are repeated at least 3 times in a row. SuperHOR hits are colored according to n. This work is described in more detail in **SNote 1**.

#### Subfamily Characterization of Human Satellites

Arrays annotated as HSat2,3 in the censat annotation for T2T-HG002 were further categorized into the 3 HSat2 subfamilies and 11 HSat3 subfamilies previously identified in Altemose et al. ^49^HuRef reads assigned to each subfamily^49^ were aligned to the T2T-HG002 reference using winnowmap (k=15) ^54^

#### Local identity calculation

Active arrays from the hg002v1.1 assembly were identified in the censat v2.0 BED file by selecting entries ending with “H1L”. Adjacent intervals (within 200 kb) were merged using bedtools. FASTA sequences for each active array were extracted with samtools and input to ModDotPlot^28^ using a 5 kb window size and delta = 0. The resultant bed files, with all-vs-all identities, were processed using a custom Python script that averages the identity values for each region’s closest neighbors (default of two regions on each side). Averaged values were assigned to a color scale determined either by a fixed minimum identity threshold (90%) or by the 10th percentile of observed values, and the results were written to BED files.

#### Lymphoblastoid cell line CDR prediction

For LCLs, mCpG wiggle tracks were generated from the processed methylation calls and visualized in IGV alongside the corresponding raw read-level mCpG signal. Sub-CDRs were manually annotated as contiguous regions in which both the wiggle-track mCpG percentage and the raw read mCpG signal showed a pronounced reduction, typically from an average of approximately 60–40% mCpG down to 25–10%. Intervening regions between sub-CDRs that did not display this marked decrease were excluded from the prediction, and CDR boundaries in LCLs were defined solely on the basis of these mCpG patterns.

#### hiPSC CDR prediction

Centromere Dip Regions in HG002 hiPSC haplotypes were identified using a sliding-window CDR caller applied to phased, per-CpG methylation bedgraph files derived from Oxford Nanopore long-read sequencing (Described in more detail **SNote 4**). For each haplotype array, a detection threshold was set at 10 percentage points below the array-wide mean methylation, accounting for the substantial variation in baseline methylation across centromeric arrays. Methylation signal was smoothed using a 10 kb sliding window stepped at 1 kb intervals (minimum 3 CpGs per window), and contiguous runs of below-threshold windows were called as candidate domains. Each candidate domain was assigned a composite reliability score (0–100) weighted by depth of hypomethylation (50%), CpG count (30%), and domain span (20%). Domains scoring ≥ 40 were retained for analysis; those scoring ≥ 50 were classified as HIGH_CONFIDENCE. CDR total span was defined as the interval from the start of the first to the end of the last HIGH_CONFIDENCE domain per haplotype.

#### Fiber-seq

To perform the Fiber-seq experiment^30^, all reagents were freshly prepared and syringe-filtered through a 0.2 μm filter before being kept on ice. Live cells (1M-5M per condition) were pelleted at 300 × *g* for 5 minutes and washed with PBS. Pelleted cells were resuspended in 1 ml of Dig-Wash buffer (0.02% digitonin, 20 mM HEPES-KOH, pH 7.5, 150 mM NaCl, 0.5 mM Spermidine, 1 Roche Complete tablet-EDTA (11873580001) per 50 ml buffer, 0.1% BSA) and incubated on ice for 5 minutes. Note that using detergents other than digitonin and Tween may reduce methylation efficiency. The nuclei suspension was split into separate tubes for each condition and spun down at 4°C at 500 × g for 3 minutes. All subsequent spins were performed under the same conditions, and all steps involving pipetting nuclei were performed with wide bore tips. The supernatant was removed, and the pellet was gently resuspended in 200 μl of Tween-Wash (0.1% Tween-20, 20 mM HEPES-KOH, pH 7.5, 150 mM NaCl, 0.5 mM Spermidine, 1 Roche Complete tablet-EDTA per 50 ml buffer, 0.1% BSA) containing the primary antibody at a 1:50 dilution. Samples were placed on a rotator at 4°C for 2 hours. Nuclei were then pelleted, washed twice with 0.95 ml Tween-Wash, and resuspended in 200 μl of Tween-Wash containing 200 nM Hia5. The concentration of Hia5 was measured using the Qubit Protein Assay Kit (Q33211). For methyltransferase binding, nuclei were placed on a rotator at 4°C for 2 hours. Nuclei were then spun down, washed twice with 0.95 ml Tween-Wash, and resuspended in 100 μl of Activation Buffer (15 mM Tris, pH 8.0, 15 mM NaCl, 60 mM KCl, 1 mM EDTA, pH 8.0, 0.5 mM EGTA, pH 8.0, 0.5 mM Spermidine, 0.1% BSA, 800 μM SAM). The nuclei were incubated at 37°C for 1 hour before spinning and resuspending in 100 μl of cold PBS. The extraction process was initiated by thawing the Extraction EB at room temperature, followed by vortexing and placing it on ice.In a separate tube, a pre-mix of 1.8 ml of NEB Monarch® HMW gDNA Tissue Lysis Buffer and 60 μl of Proteinase K was prepared. This pre-mixed lysis buffer was then added to the suspected nuclei from the experiment, followed by slow pipette mixing using a 1 ml wide-bore pipette. The samples were incubated at 56°C for 60 minutes at 300 rpm, allowing for thorough lysis. After this, protein separation was achieved by adding 900 μl of Protein Separation Solution, followed by mixing with a Hula Mixer and centrifugation at 16,000 x g for 10 minutes at 4°C. The DNA, present in the upper phase, was carefully aspirated using a wide-bore pipette tip to avoid disturbing the protein phase below. To facilitate DNA precipitation, three glass beads were added, followed by the addition of 2.5 ml of isopropanol and mixing on a rotator mixer at 10 rpm for 20 minutes. After resting the tubes at room temperature for 1 minute, the isopropanol was carefully aspirated, leaving the DNA bound to the glass beads. The samples were then washed twice with 2 ml of Wash Buffer, and the remaining wash buffer was removed by quick soft spinning. Finally, the beads were transferred into a 2 ml tube containing 200 μl of Extraction EB and incubated overnight at room temperature to allow DNA extraction. The DNA was then separated from the beads by spinning, and the final volume was adjusted to 750 μl by adding additional Extraction EB for downstream applications. To begin the data analysis, ONT Fiberseq data was aligned to the hg002v1.1 genome using winnowmap (v2.03) (https://github.com/marbl/Winnowmap). ONT dorado called MM/ML tags were filtering using modkit (v0.4.1) call-mods to 0.1 (https://github.com/nanoporetech/modkit). As per recommendation from the fibertools book (https://fiberseq.github.io/fibertools/fibertools.html). Nucleosome calls were added after filtering to individual reads using fibertools (v0.5.4) add-nucleosomes (https://github.com/fiberseq/fibertools-rs). Aggregate 6mA/A fraction modified bedMethyl files were calculated using modkit pileup (v0.4.1). Fibertools (v0.5.4) extract was used to calculate genomic positions of nucleosomes and MSPs for aggregation.

#### Cell culture and passage

HG002 lymphoblastoid cells (GM24385; Coriell Institute; mycoplasma tested) were maintained in RPMI 1640 (Gibco, 11875093 or 21875034; 2 g/L glucose) supplemented with 2 mM L-glutamine provided as GlutaMAX (Gibco, 35050061), 15% fetal bovine serum (Gibco, 26140079; lot-selected from a reputable source, Southern Hemisphere origin preferred), and 1% penicillin–streptomycin (Gibco, 15070063) at 37°C in a humidified atmosphere of 5% CO2.

HG002 LCL were passaged at a split ratio of 1:3 after growing to a density of 0.9 to 1.0 x 10^6 viable cells/ml. At each passage, twelve million cells were pelleted at 300 x g for 5 minutes and resuspended in cell freezing media (Sigma-Aldrich C6295) at a concentration of 5.0 million cells/ml. A Mr. Frosty freezing container (Thermo Scientific, 5100-0001) was used to cryopreserve the harvested cells at −80°C. Ten million cells were cryopreserved for DiMeLo-seq, and two million cells cryopreserved for future culturing.

GM27730*B iPSC (Coriell Institute; mycoplasma tested) were maintained in DMEM/F12 supplemented with 20% KnockOut Serum Replacement (KOSR) and 10 ng/ml basic fibroblast growth factor (bFGF) at 37 °C in a humidified atmosphere of 5% CO2 on CF1 mouse embryonic fibroblast (MEF) feeders prepared on 0.1% gelatin-coated surfaces. Cells were passaged every 5–7 days at a 1:6 split ratio using TrypLE Express.

GM26105*F iPSC (Coriell Institute; mycoplasma tested) were maintained in mTeSR1 at 37 °C in a humidified atmosphere of 5% CO2 on Matrigel-coated surfaces. Cells were passaged every 5–7 days at a 1:8 split ratio using Versene and recovered in 2 wells of a 6-well plate.

#### DiMeLo-seq

All reagents were freshly prepared and syringe-filtered through a 0.2 μm filter before being kept on ice. Cells (1M-5M per condition) were pelleted at 300 × *g* for 5 minutes and washed with PBS. For experiments targeting CENP-A, H3K9me3, and CENP-C, live cells were used. The standard DiMeLo-seq protocol’s^19^ nuclear isolation can then be continued. Pelleted cells were resuspended in 1 ml of Dig-Wash buffer (0.02% digitonin, 20 mM HEPES-KOH, pH 7.5, 150 mM NaCl, 0.5 mM Spermidine, 1 Roche Complete tablet-EDTA (11873580001) per 50 ml buffer, 0.1% BSA) and incubated on ice for 5 minutes. Note that using detergents other than digitonin and Tween may reduce methylation efficiency. The nuclei suspension was split into separate tubes for each condition and spun down at 4°C at 500 × g for 3 minutes. All subsequent spins were performed under the same conditions, and all steps involving pipetting nuclei were performed with wide bore tips. The supernatant was removed, and the pellet was gently resolved in 200 μl of Tween-Wash (0.1% Tween-20, 20 mM HEPES-KOH, pH 7.5, 150 mM NaCl, 0.5 mM Spermidine, 1 Roche Complete tablet-EDTA per 50 ml buffer, 0.1% BSA) containing the primary antibody at a 1:50 dilution. An antibody targeting CENP-A (ADI-KAM-CC006-E), H3K9me3 (ab8898), or CENP-C (PD030) was used. Samples were placed on a rotator at 4°C for 2 hours. Nuclei were then pelleted, washed twice with 0.95 ml Tween-Wash, and resolved in 200 μl of Tween-Wash containing 200 nM pA-Hia5 for CENP-C and H3K9me3 and anti-mouse nanobody for CENP-A. The concentration of pA-Hia5/anti-mouse nanobody was measured using the Qubit Protein Assay Kit (Q33211). For methyltransferase binding, nuclei were placed on a rotator at 4°C for 2 hours. Nuclei were then spun down, washed twice with 0.95 ml Tween-Wash, and resuspended in 100 μl of Activation Buffer (15 mM Tris, pH 8.0, 15 mM NaCl, 60 mM KCl, 1 mM EDTA, pH 8.0, 0.5 mM EGTA, pH 8.0, 0.5 mM Spermidine, 0.1% BSA, 800 μM SAM). The nuclei were incubated at 37°C for 1 hour before spinning and resuspending in 100 μl of cold PBS.

#### DNA extraction

The nuclei were combined with 1.8 ml NEB Monarch HMW gDNA Tissue Lysis Buffer and 60µL proteinase K (800 units/ml) in a 5 ml Eppendorf DNA LoBind tube, and then gently mixed by pipetting with a wide-bore tip. The samples were incubated at 56°C for 60 minutes at 300 rpm, allowing for thorough lysis. After this, protein separation was achieved by adding 900 μl of Protein Separation Solution, followed by mixing with a rotator mixer for 10 minutes at 10 rpm and centrifugation at 16,000 x g for 10 minutes at 4°C. The upper phase containing the DNA was carefully aspirated by pipetting with a wide-bore tip and transferred to a fresh 5 ml Eppendorf DNA LoBind tube.To facilitate DNA precipitation, three NEB Monarch DNA Capture Beads were added, followed by the addition of 2.5 ml of isopropanol and mixing with a rotator mixer at 10 rpm for 20 minutes. After resting the tubes at room temperature for 1 minute, the isopropanol was carefully aspirated, leaving the DNA bound to the glass beads. The glass beads were washed twice and transferred to a bead retainer where any remaining wash buffer was removed by a soft momentary centrifugation. Finally, the beads were transferred to a tube containing 200 µl elution buffer (ONT Extraction EB) and allowed to incubate at room temperature overnight. The DNA was then separated from the beads by centrifugation in a bead retainer for 1 minute at 1000 x g. The final volume was adjusted to 750µl elution buffer.

#### Nanopore Ultra-long PromethION library preparation and sequencing

In a 1.5 ml Eppendorf DNA LoBind tube, 6 µl fragmentation enzyme (FRA) was diluted with 244 µl dilution buffer (FDB), and mixed by pipetting. The resulting solution was then combined with 750 µl sample DNA and mixed thoroughly by pipetting fifteen times with a wide-bore tip. The sample reaction was then incubated at room temperature for 10 minutes, and placed on ice for 5 minutes. After cooling, 5 µl rapid adapter (RA) was added to the sample, mixed by pipetting with a wide-bore tip five times, and incubated for 30 minutes at room temperature. For DNA precipitation, thaw Elution Buffer and Precipitation buffer components, briefly centrifuge, and vortex to mix. Keep all kit components on ice. Following the addition of 500 µl of Precipitation Reagent using a regular pipette tip, we gently rotated the sample on a rotator mixer for 20 minutes at 10 rpm to promote DNA precipitation. Centrifuge sample at 2000xg for three minutes. Carefully remove the supernatant, taking care not to aspirate the DNA. Spin down the tube at 2000xg for three minutes and remove any residual supernatant, avoiding DNA aspiration. Add 275 µl of Elution Buffer and incubate overnight at room temperature. Gently mix the DNA library ten times with a P1000 wide-bore pipette tip to prevent sample heterogeneity. For storage, it is recommended to store libraries in Eppendorf DNA LoBind tubes at 4°C for short-term use or reloading flow cells between washes.

#### Promethion adaptive sampling

For each adaptive sampling run, we provided the PromethION platform with a reference file containing CHM13 coordinates corresponding to mappable regions (https://github.com/yxu405/HG002_epi_paper/blob/f56f7755f2d13d7d07ca65e6babd959fb46aa6f2/adaptive_s <u>equencing/chm13v2.0_allalpha.slop100k.complement.bed</u>). This BED file defines the complement of alpha-satellite (αSat) repeat regions expanded by 100 kb flanks, excluding easily mappable portions of the ∼3.3 Gb T2T-CHM13v2.0 genome while targeting ∼190 Mb (6.2%) of pericentromeric and centromeric repeats that are typically refractory to short-read mapping. The PromethION sequencer performed real-time basecalling and mapping of the first few hundred base pairs of each DNA molecule. All samples were sequenced on a PromethION 48 sequencer (PRO-SEQ048) using R10.4.1 Flow Cells (FLO-PRO114M). The SKU wash kits (EXP-WSH004) were used in this process. Critically, we ran the adaptive sampling sequencing run in DEPLETE mode for this bed file. If the DNA molecules mapped to the CHM13 reference coordinates (i.e., mappable euchromatin), they were rejected; otherwise, unmapped molecules—enriched for centromeric heterochromatin—were allowed to be fully sequenced. This approach specifically targets centromeres because they comprise long tracts of highly repetitive αSat DNA (2.8% of the genome alone), which evades alignment due to their length, homogeneity, and complexity.

### DiMeLo-seq Analysis

#### Basecalling and Filtering

Raw Nanopore sequencing data were basecalled from POD5 files using Dorado (Oxford Nanopore Technologies) to simultaneously detect nucleotide sequences and base modifications. Basecalling was performed with the model dna_r10.4.1_e8.2_400bps_sup@v4.2.0, which supports the detection of both 5-methylcytosine and N6-methyladenine. All basecalling was executed on the Phoenix HPC cluster using Dorado v0.6.1 to generate BAM files containing per-base methylation tags for 5mC and 6mA. These unaligned BAM files were then processed using a batch-submitted SLURM script (https://github.com/yxu405/HG002_dimelo_raw_data_processing) that performs conversion, alignment, and post-processing optimized for long-read Nanopore data. The script converts BAM to FASTQ preserving modification tags, aligns to the HG002 reference using Winnowmap, and generates sorted and indexed primary alignments.

#### mA visualization and aggregate quantification (include dosage analysis & aggregated metaplots)

Individual reads were visualized using IGV ^34^ to inspect methylation patterns in the centromere regions. Chromosomal paternal-maternal CENP-A, CENPC, correlations were visualized as scatterplots (x-axis: paternal mA/A density; y-axis: maternal mA/A density) with size-difference color-coding. Outliers were identified as chromosomes with standardized residuals |z| > 3 from the identity line (y = x), where residuals = maternal - paternal density, z = |(residual - μ)/σ|.

Furthermore, to visualize the presence of epigenetic marks (CENP-A, CENP-C, and H3K9me3) in chromosomal sub-CDRs, predicted CDR coordination bed files were used to extract reads in those genomic locations. For each chromosome, three metrics were compared between paternal and maternal haplotypes: mA/A density, desired read length, and their product (density × length) as an estimate of integrated CENP-A/C dosage. Pearson correlation coefficients and mean squared error from the identity line (y = x) were calculated for each metric. Outlier chromosomes were defined as those with standardized residuals |z| > 3, where residuals represent the maternal − paternal difference and z-scores are computed relative to the cross-chromosomal mean and standard deviation of those residuals.

#### Single-molecule analysis of sub-CDR co-occupancy

Single-molecule 6mA and 5mC densities were calculated for individual reads in CDR and non-CDR active αSat defined regions. For each read overlapping a defined CDR or non-CDR region, the modification score array was extracted from the MM/ML BAM tags and filtered by a minimum quality score threshold (6mA: ≥230/255; 5mC: ≥210/255). non-CDR regions were defined as active αSat intervals 10kb away from the CDRs, tiled into 5,000 bp non-overlapping windows with up to 15 windows retained per chromosome homolog.

To identify per-mark density thresholds distinguishing CDR from non-CDR regions, a decision tree algorithm was implemented in Python. Per-read densities from CDR and non-CDR regions were pooled and sorted, and candidate thresholds were evaluated at midpoints between consecutive density values. The threshold minimizing the weighted Gini impurity across the two resulting partitions was selected as optimal. This procedure was applied independently for each epigenetic mark, yielding the following genome-wide thresholds: CENP-A 6mA density: 0.011; CENP-C 6mA density: 0.014; H3K9me3 6mA density: 0.015. These thresholds were used to classify individual reads as CDR-enriched or non-CDR at single-molecule resolution.

#### Epigenetic comparisons between flanking and central CDRs in low-passage LCLs, high-passage LCLs

All raw reads were visualized in IGV ^34^. The 5mC methylation island regions are defined as regions exhibiting CpG methylation enrichment in low-passage LCLs that is absent in high-passage LCLs. To characterize the epigenetic status of these differentially methylated island regions, per-chromosome mean 5mC densities were calculated for center CDR, flanking CDR, and non-CDR baseline regions using single-molecule CpG methylation calls extracted from DiMeLo-seq BAM files. CENP-A and H3K9me3 5mC densities were then normalized to the non-CDR active array baseline mean, and log2 fold-changes were computed per chromosome. Distributions of these log2 fold-changes were compared between low- and high-passage conditions for center and flanking CDR subregions using Welch’s two-sample t-tests, with effect sizes reported as Cohen’s d.

#### Epigenetic comparisons between differentially methylation regions in low-passage LCLs, high-passage LCLs

To identify differentially methylated regions in-between sub-CDRs, CpG methylation profiles were compared between low- and high-passage LCLs across the chromosomal CDRs. Sub-CDR boundaries were first identified in both passage conditions. Regions exhibiting CpG methylation enrichment in the low-passage LCL that transitioned to a depleted state in the high-passage LCL were isolated by subtracting the high-passage sub-CDR coordinates from those of the low-passage condition. The resulting intervals — regions present in the low-passage methylation landscape but absent in the high-passage — were marked.

Next, to characterize the epigenetic state of differentially methylated regions, CENP-A, 5mC, and H3K9me3 signal densities were extracted from the differentiated regions. Per-read modification densities were calculated from DiMeLo-seq BAM files, with 6mA calls filtered to a minimum quality of 230/255 and 5mC calls to 230/255. The six resulting density tracks (CENP-A, 5mC, and H3K9me3 for each passage condition) were merged by genomic coordinates. Signal densities within each compartment were then normalized to chromosome-specific non-CDR baseline values computed separately for each mark and passage condition, and log2 fold-changes were calculated per region. Distributions of log2 fold-changes were visualized as boxplots with individual chromosome-level data points overlaid, enabling direct comparison of epigenetic signal enrichment or depletion across passage conditions.

#### Epigenetic comparisons between differentially methylation regions in low-passage LCLs, high-passage LCLs

To compare CENP-A and CpG methylation enrichment at CDRs between the low-passage LCL and iPSC cell states, per-read 6mA and 5mC densities were extracted from DiMeLo-seq BAM files across active αSat array windows and CDR windows. For each mark, a chromosome-specific background mean density was computed from active αSat windows after excluding any window falling within 50 kb of a CDR interval, ensuring that background estimates reflected chromatin state distal from CDR-proximal regions. CDR window densities were then normalized to this background mean, and log2 fold-changes were calculated per CDR window. Distributions of log2 fold-changes were compared between the low-passage LCL and iPSC and visualized as boxplots with individual CDR-level data points overlaid.

#### Single-molecule read clustering based on methylation

CDRs were classified into three categories: edge regions (outermost CDR boundaries), sinking regions (small disappearing sub-domains), and stable island regions (persistently hypomethylated CDR cores), each comprising ten genomic intervals defined in BED format. For each region, CpG dinucleotides were enumerated directly from the reference sequence.

Per-read CpG methylation matrices were constructed by extracting 5mC modification probabilities at each CpG site within a given CDR interval, retaining only reads spanning at least 95% of the region (windowFrac = 0.95). Each matrix was organized with individual reads as rows and CpG positions as columns, with values representing continuous methylation probability scores (0 = unmethylated, 1 = fully methylated), yielding a high-resolution, read-level view of methylation heterogeneity across each CDR.

These per-read methylation matrices were used as input to HDBSCAN (Hierarchical Density-Based Spatial Clustering of Applications with Noise) to identify subpopulations of reads sharing distinct methylation patterns within each CDR. Hierarchical linkage was computed using Ward’s minimum variance method, which minimizes within-cluster variance at each merge step and is well-suited to identifying compact, well-separated clusters in continuous data. Clustering parameters were set as follows: minimum cluster size = 5, minimum samples = 3, and cluster selection epsilon = 0.005. Reads assigned to noise (cluster label −1) were excluded from interpretation. Clustering was performed independently for each of the ten regions within each CDR category (edge, sinking, stable), allowing region-specific methylation substructure to be resolved without assumptions about the number of clusters. The resulting cluster assignments were visualized as heatmaps of per-read methylation probability, ordered by cluster membership, with accompanying dendrograms to illustrate hierarchical relationships between read groups.

## QUANTIFICATION AND STATISTICAL ANALYSIS

All statistical analyses were performed in Python using the SciPy, NumPy, and Pandas libraries. Statistical details for each comparison are reported in the corresponding figure legends.

### Dosage analysis

To assess allele-specific CENPA dosage, mean signal density and total CDR length in bp were computed separately for paternal and maternal chromosomes. A dosage proxy was calculated as the product of density and CDR length for each chromosome and haplotype. Pearson correlation coefficients and mean squared error (MSE) were computed between paternal and maternal values for each of the three metrics (density, length, and density × length) using NumPy. Chromosomes were flagged as outliers if their paternal–maternal residual exceeded a Z-score threshold of |Z| > 3.0, where Z-scores were computed from the mean and standard deviation of the residual distribution across all chromosomes. Analyses were performed separately for young, old, and iPSC-derived cells. N is the number of chromosomal haploid types.

### Read-level density thresholding

To classify individual reads as CDR or non-CDR based on their modification density, an optimal density threshold was determined for each chromosome for CENPA, H3K9me3 and mCpG independently. Non-CDR active alpha-satellite regions were used as the background reference. For each chromosome, read-level modification densities were computed across 5,000 bp segments within CDR and non-CDR regions, with up to 15 segments sampled per chromosome from non-CDR regions to balance class representation. The optimal threshold was identified by exhaustive search over all midpoints between sorted density values, selecting the threshold that minimized the weighted Gini impurity across CDR and non-CDR classes. Thresholds were determined separately for 6mA (filtering value ≥ 230) and 5mCG (filtering value ≥ 230) modification calls. Thresholds derived from this procedure were subsequently applied to classify reads in downstream single chromatin fiber analysis.

### Density normalization

For each histone mark (CENPA, H3K9me3, mCpG), mean signal density was computed per chromosome for each genomic region (center CDR, flanking CDR, and Non-CDR). Per-chromosome log2 fold-changes were calculated by dividing the regional mean density by a chromosome-specific baseline density derived from Non-CDR regions, followed by a log2 transformation. Separate baseline values were computed for young and old animals to control for age-associated global differences in signal levels. Values with a fold-change of zero or below were excluded prior to log2 transformation.

### Epigenetic comparisons between early and late passaged cells, or LCL and iPSCs

Differences in log2 fold-change between early and late passaged cells were assessed using a paired two-tailed Student’s t-test, with chromosomes serving as paired units. Only chromosomes present in both early and late datasets were included in statistical comparisons. The number of shared chromosomes per comparison is reported in the results. A significance threshold of p < 0.05 was applied. n represents the number of chromosomes in the HG002 cell line.

### HORhap age analysis

The relative averaged sizes of younger and older HORhap arrays were 0.64 and 0.36, respectively. Kinetochore positions were observed 28 times on younger arrays and 9 times on older arrays. A one-sided exact binomial test was used to assess whether younger arrays were overrepresented relative to their expected proportion. The null hypothesis of no positional preference could not be rejected (p = 0.092). P-values were calculated in Python using SciPy (scipy.stats.binomtest(28, 37, 0.64, alternative=“greater”)).

### hiPSC CDR prediction

Centromere Dip Regions in HG002 hiPSC haplotypes were identified using a sliding-window CDR caller applied to phased, per-CpG methylation bedgraph files derived from Oxford Nanopore long-read sequencing. For each haplotype array, a detection threshold was set at 10 percentage points below the array-wide mean methylation, accounting for the substantial variation in baseline methylation across centromeric arrays. Methylation signal was smoothed using a 10 kb sliding window stepped at 1 kb intervals (minimum 3 CpGs per window), and contiguous runs of below-threshold windows were called as candidate domains. Each candidate domain was assigned a composite reliability score (0–100) weighted by depth of hypomethylation (50%), CpG count (30%), and domain span (20%). Domains scoring ≥ 40 were retained for analysis; those scoring ≥ 50 were classified as HIGH_CONFIDENCE. CDR total span was defined as the interval from the start of the first to the end of the last HIGH_CONFIDENCE domain per haplotype. CDR calling was performed using cdr_caller_v2.py (available as Supplemental Code), implemented in Python 3 using NumPy and Matplotlib.

## Notes

### Competing Interest Statement

Nicolas Altemose is named as an inventor on patent applications related to DiMeLo-seq.

