## Supplementary Information for "Haplotype-resolved centromeric chromatin organization from a complete diploid human genome"


Supplementary Figure 1: Diploid centromeric satellite annotations for HG002 2
Supplementary Figure 2: Chromosome 9 HSat3 variation 3
Supplementary Figure 3: Chromosome 1 HSat SF variation 4
Supplementary Figure 4: Chromosome 3 and 4 Alpha and HSat1A variation 5
Supplementary Figure 5: Moddotplot with paired 1D moddotplot track 6
Supplementary Figure 6: HOR Structural Variant (StV) profiles 10
Supplementary Figure 7: Chromosome 13 Structural Variant (StV) and HORhap analysis 16
Supplementary Figure 8: CDR p-arm proximal regions 17
Supplementary Figure 9: CDR association with homogenized regions 18
Supplementary Figure 10: Total CDR span per haplotype in early passage LCL (HG002) 19
Supplementary Figure 11: CDR (subCDR and span) vs active array size 20
Supplementary Figure 12: Coverage estimates of adaptive sampling 21
Supplementary Figure 13: Clustering methylation data into groups 22
Supplementary Figure 14: Fiber-seq nucleosome organization relative to CENP-B motifs 23
Supplementary Figure 15: CENP-C DiMeLo-seq 6mA density (early passage) 24
Supplementary Figure 16: Methylation frequency across active arrays in early vs late passage 25
Supplementary Figure 17: Difference in sub-CDR structure in early vs late passage 26
Supplementary Figure 18: Enrichment within sub-CDRs across early- and late-passage 27
Supplementary Figure 19: Comparison between HG002 early passage and hiPSC CDRs 28
Supplementary Figure 20: CENP-A DiMeLo-seq Density in hiPSCs 29
Supplementary Figure 21: CENP-A DiMeLo-seq density in late passage lines 30
Supplemental Note 1: Higher-order repeat haplotype (HORhap) analysis 31
Supplemental Note 2: Centromere mappability assessment and aligner evaluation 42
Supplemental Note 3: Quality Control and Characterization of hiPSC Lines 45
Supplemental Note 4: HG002 hiPSC CDR predictions 48
References 65

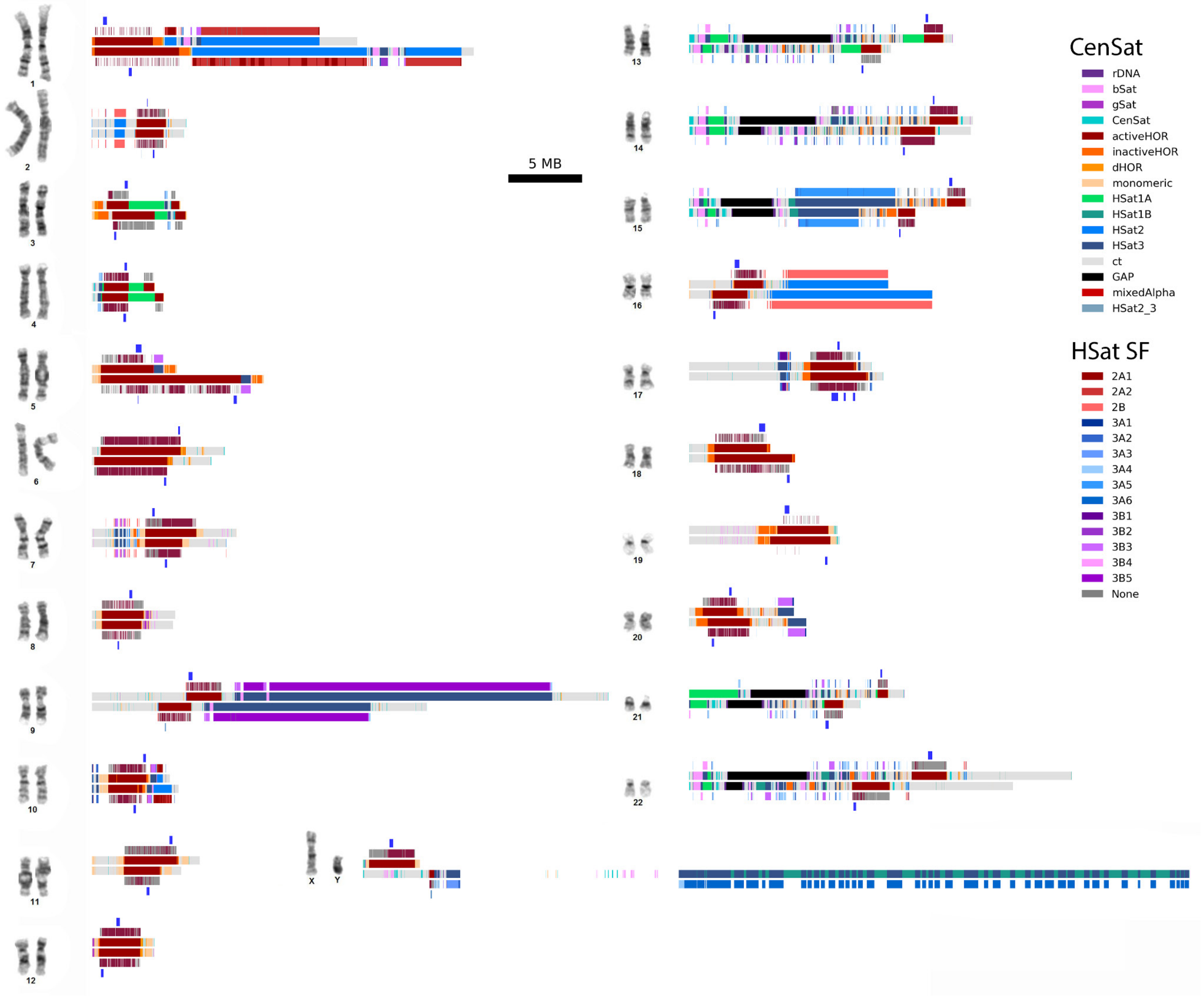


**Supplementary Figure 1: Diploid centromeric satellite annotations for HG002**

HG002 g-banding karyotype images displaying centromeric satellite (censat) annotation tracks for all chromosomes in the diploid HG002 genome, generated using an automated annotation method. For each chromosome, the maternal haplotype is shown on top and the paternal haplotype on the bottom. Annotation tracks are arranged symmetrically from the center outward and include: (1) cenSat family annotations, (2) alpha-satellite higher-order repeat (HOR) haplotypes and satellite superfamilies (HSat SF), and (3) centromeric dip regions (highlighted in blue). Alpha-satellite HOR haplotypes are colored on a gradient from younger HORs (red) to older HORs (grey)

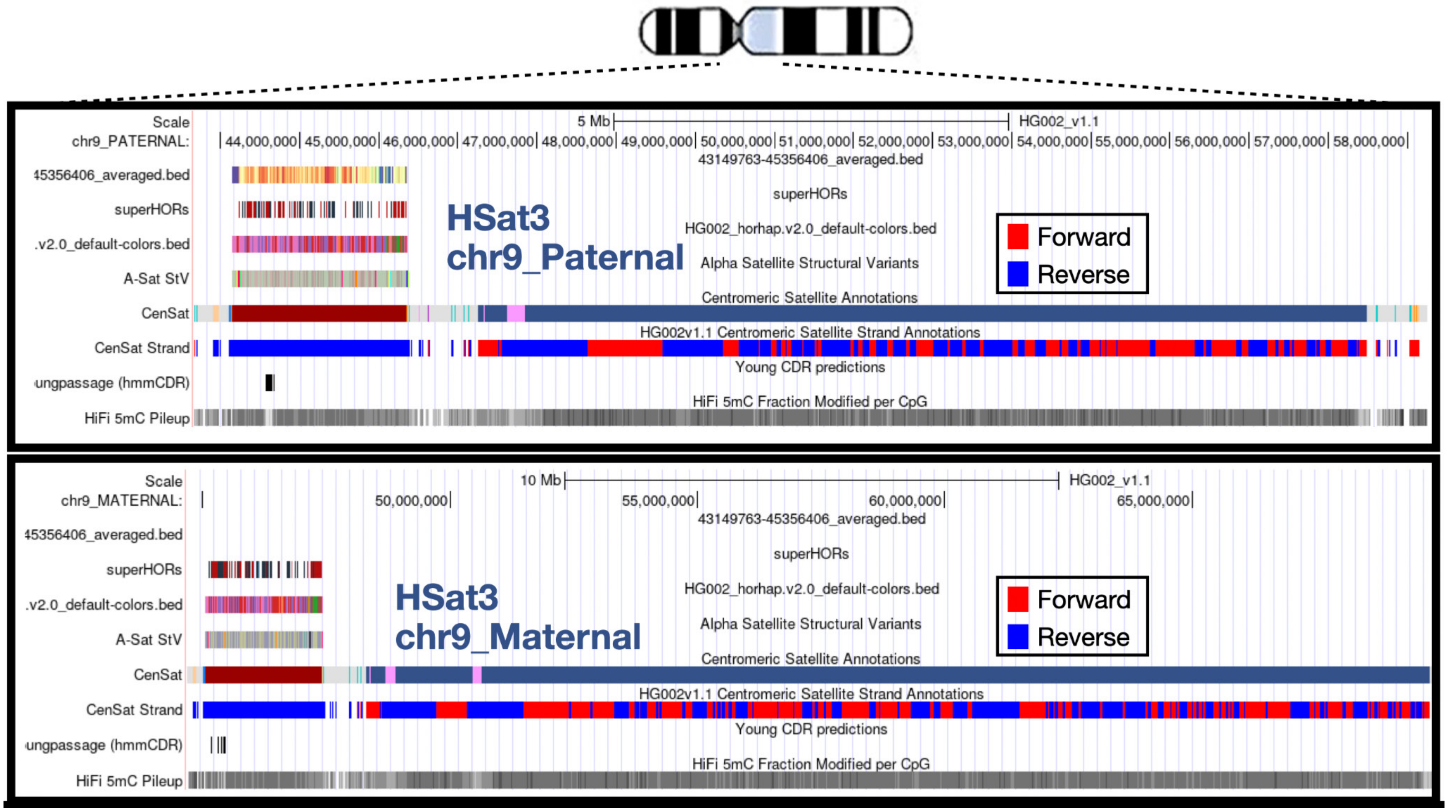


**Supplementary Figure 2: Chromosome 9 HSat3 variation.**

UCSC Genome Browser track hub [1] visualizations of the centromeric region of chromosome 9 for the maternal (chr9_maternal: 45–70 Mb) and paternal (chr9_paternal: 44–58 Mb) haplotypes, displaying cenSat satellite annotations. Unlike most satellite families, which are arranged head-to-tail in a single orientation (e.g., alpha satellite, dark red, cenSat track), Human Satellite 3 (HSat3, dark blue) on chromosome 9 exhibits both length variation and major shifts in repeat orientation across the array. Forward and reverse HSat3 orientations are shown in red and blue, respectively, revealing multiple megabase-scale strand domains arranged in large blocks. The order and polarity of these orientation segments differ between maternal and paternal chromosomes, consistent with large internal inversions and strand-polarized repeat substructure across the HSat3 array.

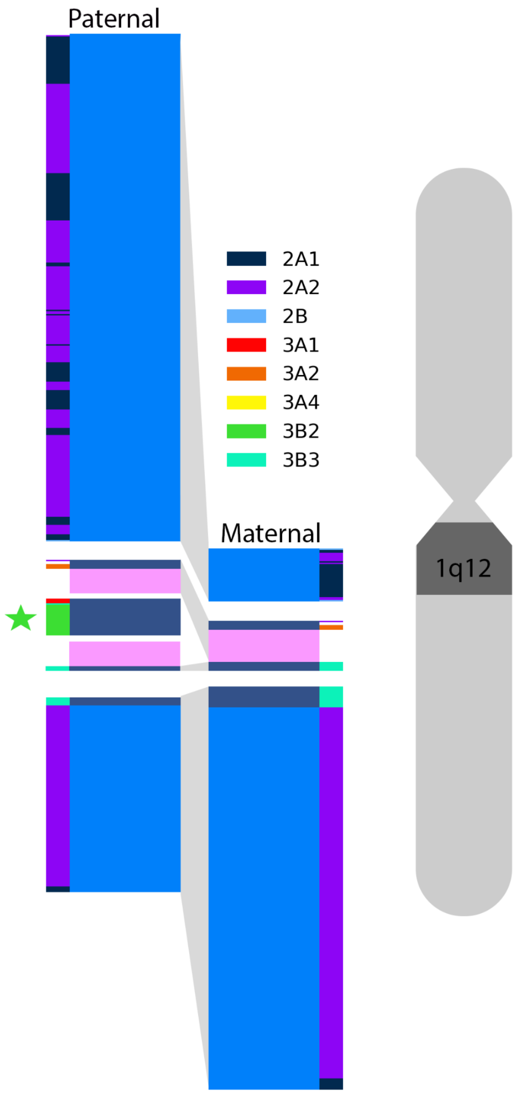


**Supplementary Figure 3: Chromosome 1 HSat SF variation**

Comparison of Chromosome 1q12 region between paternal and maternal haplotypes, large Hsat2 arrays have been truncated and are not to scale. Outer columns represent HSat superfamilies, and thicker inner columns represent CenSat annotation. Homologous arrays are connected. HSat2 on chromosome 1 differs by 6.74 Mb between homologs and contains distinct sequence compositions. The paternal haplotype contains a unique HSat3B2 array marked with a star and an adjacent beta-satellite sequence, while the paternal haplotype contains an additional ~500 kb polymorphic block of HSat3 with an adjacent beta-satellite sequence.

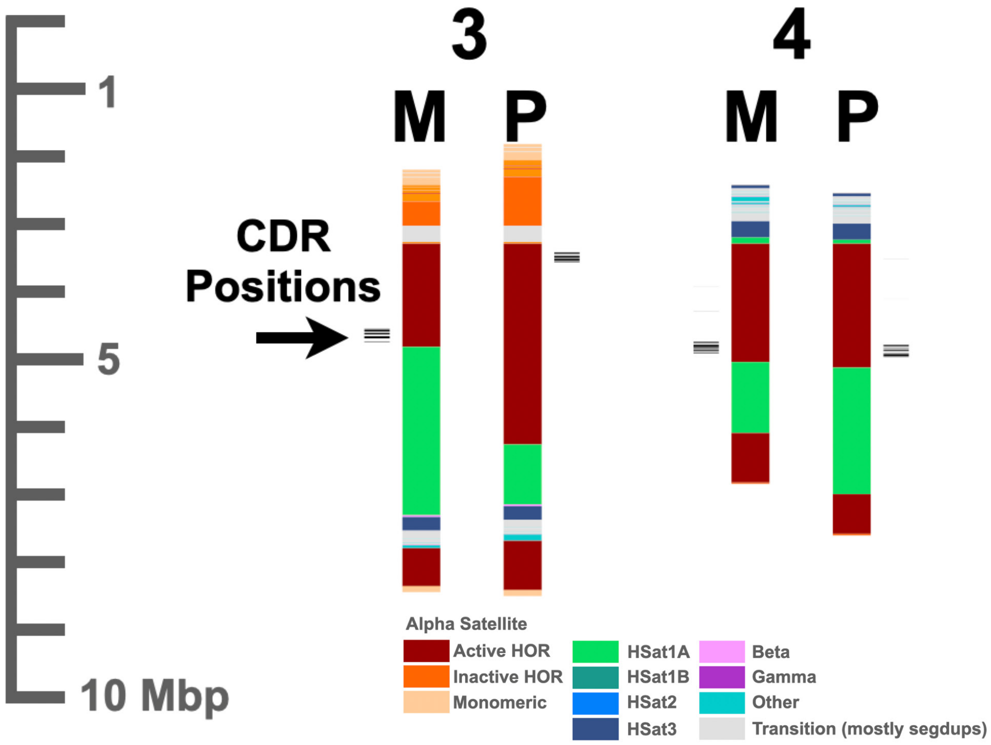


**Supplementary Figure 4: Chromosome 3 and 4 Alpha and HSat1A variation**

CenSat annotations are shown for the maternal (M) and paternal (P) haplotypes of chromosomes 3 and 4, with satellite subfamily identity indicated by color (see key). Scale bars are provided for each haplotype. Centromere dip regions (CDRs), which mark the position of the kinetochore-associated active alpha-satellite (αSat) array, are indicated by black tick marks; the CDR on chromosome 3M is highlighted by an arrow. These chromosomes illustrate two distinct forms of centromeric structural variation. First, interspersed HSat1A arrays (green) are embedded within the active αSat arrays on both chromosomes, splitting the array into discontinuous blocks and differing in size between haplotypes (~0.61 Mb on chr3; ~1.1 Mb on chr4). Second, the active αSat arrays differ in total length between haplotypes, reflecting haplotype-specific restructuring of kinetochore-associated centromeric domains. On chromosome 3, additional complexity is observed in the form of interspersed centromere transition (CT) regions containing lncRNA candidate loci (grey), as well as HSat3 and other smaller centromeric satellite classes, indicating a more structurally heterogeneous array organization relative to chromosome 4.

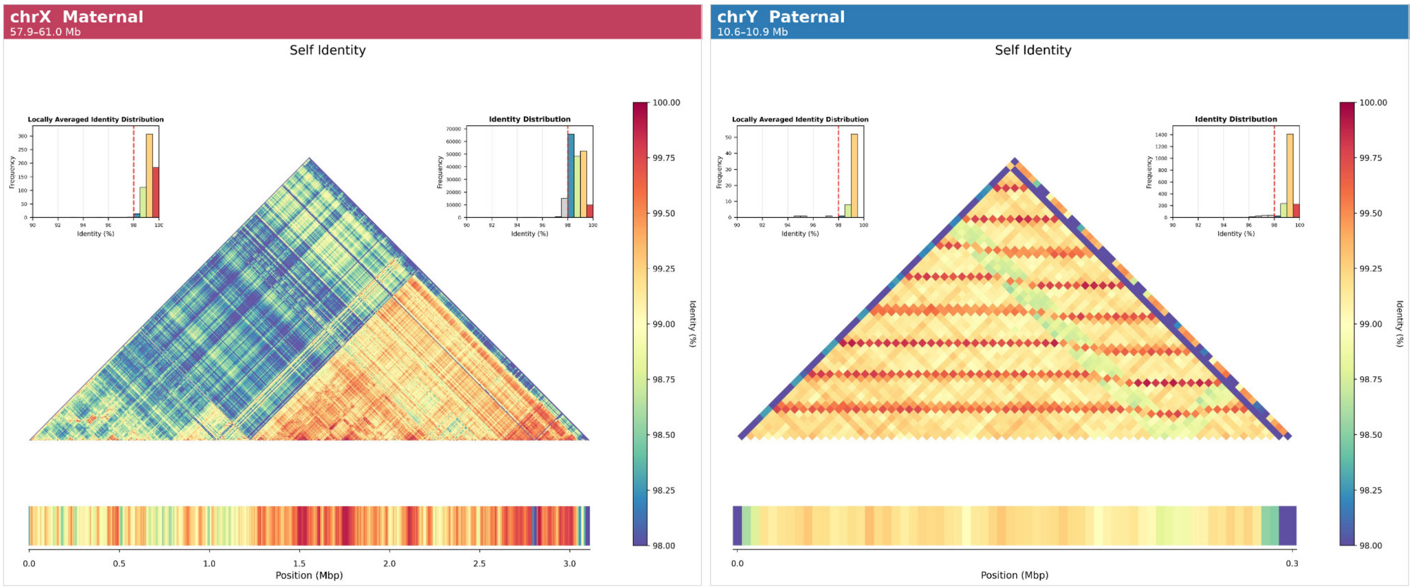

chrX Maternal
57.9–61.0 Mb
Self Identity
Locally Averaged Identity Distribution
Identity Distribution
Frequency
Identity (%)
Identity (%)
Position (Mbp)
chrY Paternal
10.6–10.9 Mb
Self Identity
Locally Averaged Identity Distribution
Identity Distribution
Position (Mbp)

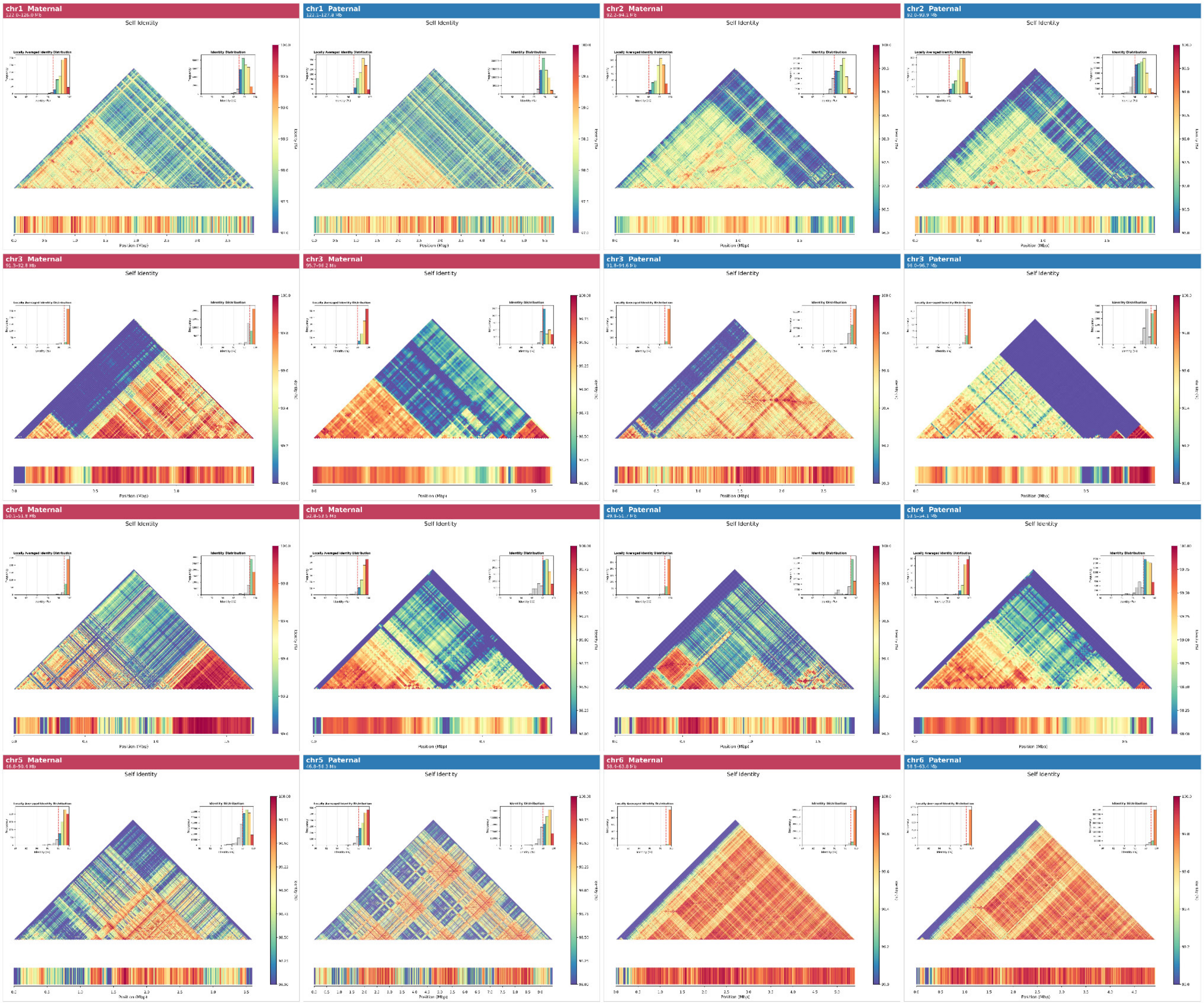
chr1 Maternal
Self Identity
chr1 Paternal
Self Identity
chr2 Maternal
Self Identity
chr2 Paternal
Self Identity
chr3 Maternal
Self Identity
chr3 Maternal
Self Identity
chr3 Paternal
Self Identity
chr3 Paternal
Self Identity
chr4 Maternal
Self Identity
chr4 Maternal
Self Identity
chr4 Paternal
Self Identity
chr4 Paternal
Self Identity
chr5 Maternal
Self Identity
chr5 Paternal
Self Identity
chr6 Maternal
Self Identity
chr6 Paternal
Self Identity
Position (Mbp)

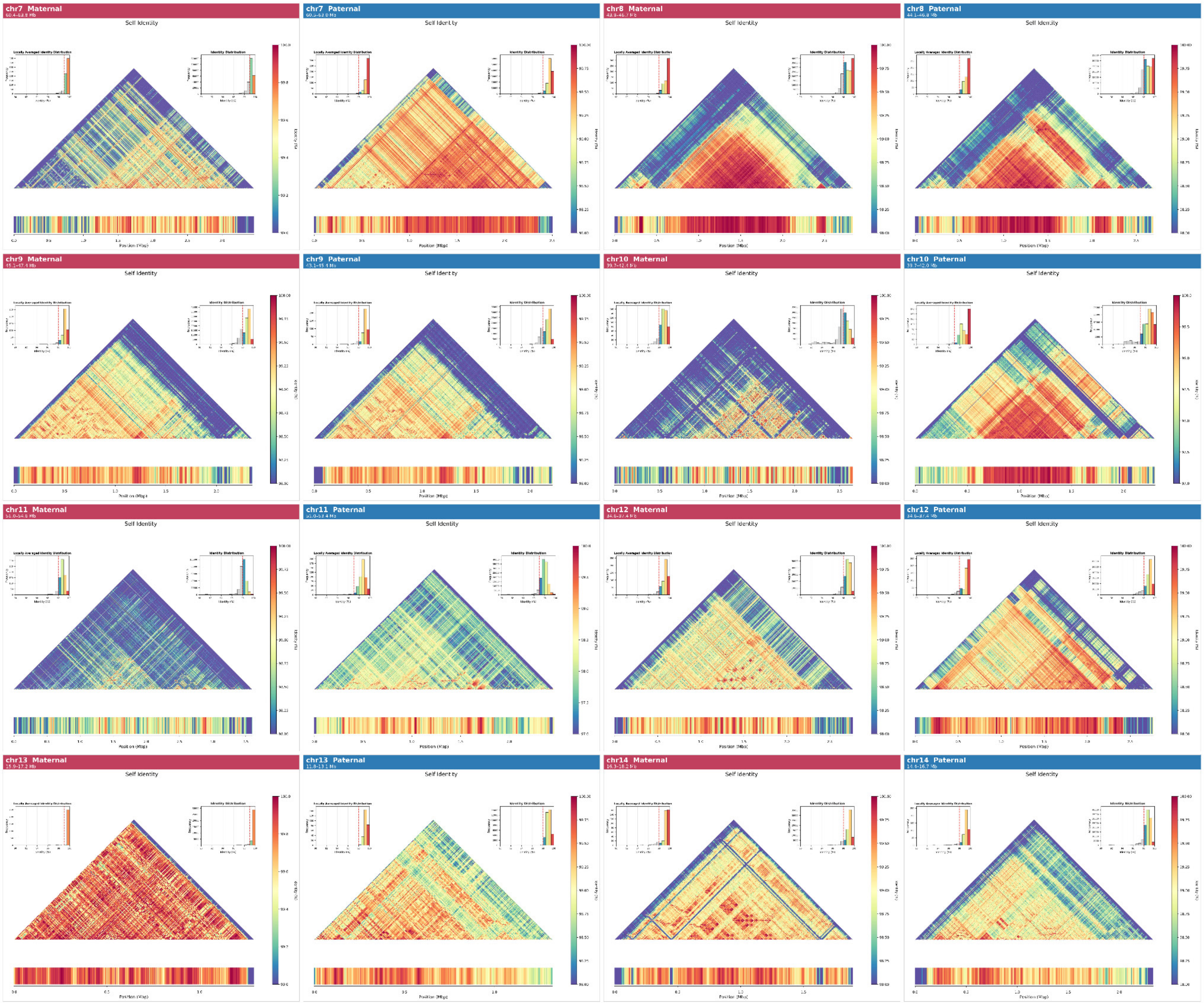

chr7 Maternal
chr7 Paternal
chr8 Maternal
chr8 Paternal
chr9 Maternal
chr9 Paternal
chr10 Maternal
chr10 Paternal
chr11 Maternal
chr11 Paternal
chr12 Maternal
chr12 Paternal
chr13 Maternal
chr13 Paternal
chr14 Maternal
chr14 Paternal
Self Identity
Locally Averaged Identity Distribution
Identity Distribution
Position (Mbp)

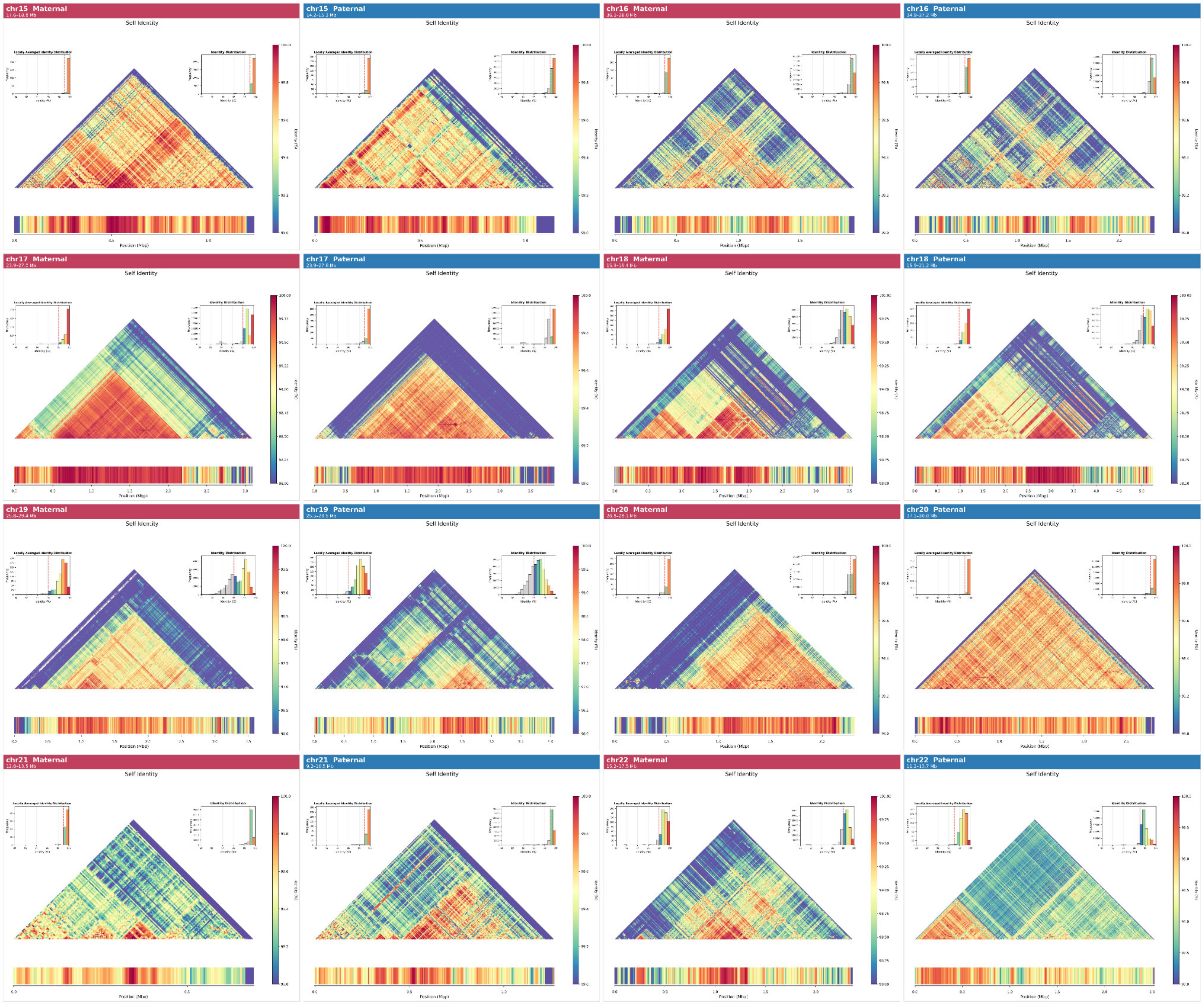


**Supplementary Figure 5: Moddotplot with paired 1D moddotplot track.**

Triangular self-identity heatmaps are shown for the active alpha-satellite HOR (αSat) arrays of each chromosome, with maternal (M) and paternal (P) haplotypes displayed separately. Active arrays were identified from the HG002v1.1 assembly using the cenSat v2.0 annotation by selecting entries with the suffix "H1L." Adjacent intervals within 200 kb were merged, and FASTA sequences for each array were extracted and input to ModDotPlot[2] (window size = 5 kb, δ = 0) to generate all-vs-all pairwise identity values. Identity values were locally averaged across the two nearest neighboring windows on each side using a custom Python script. The resulting values are displayed on a color scale ranging from a fixed lower bound of 90% identity (or the 10th percentile of observed values if higher) to 100%, with warmer colors (red/orange) indicating higher identity and cooler colors (blue/purple) indicating lower identity. The bar below each triangle represents the locally averaged identity at each position along the array. Inset histograms show the distributions of locally averaged identity (left) and raw all-vs-all pairwise identity (right); the red dashed line indicates the mean. Genomic coordinates (GRCh38) are indicated in the panel header.

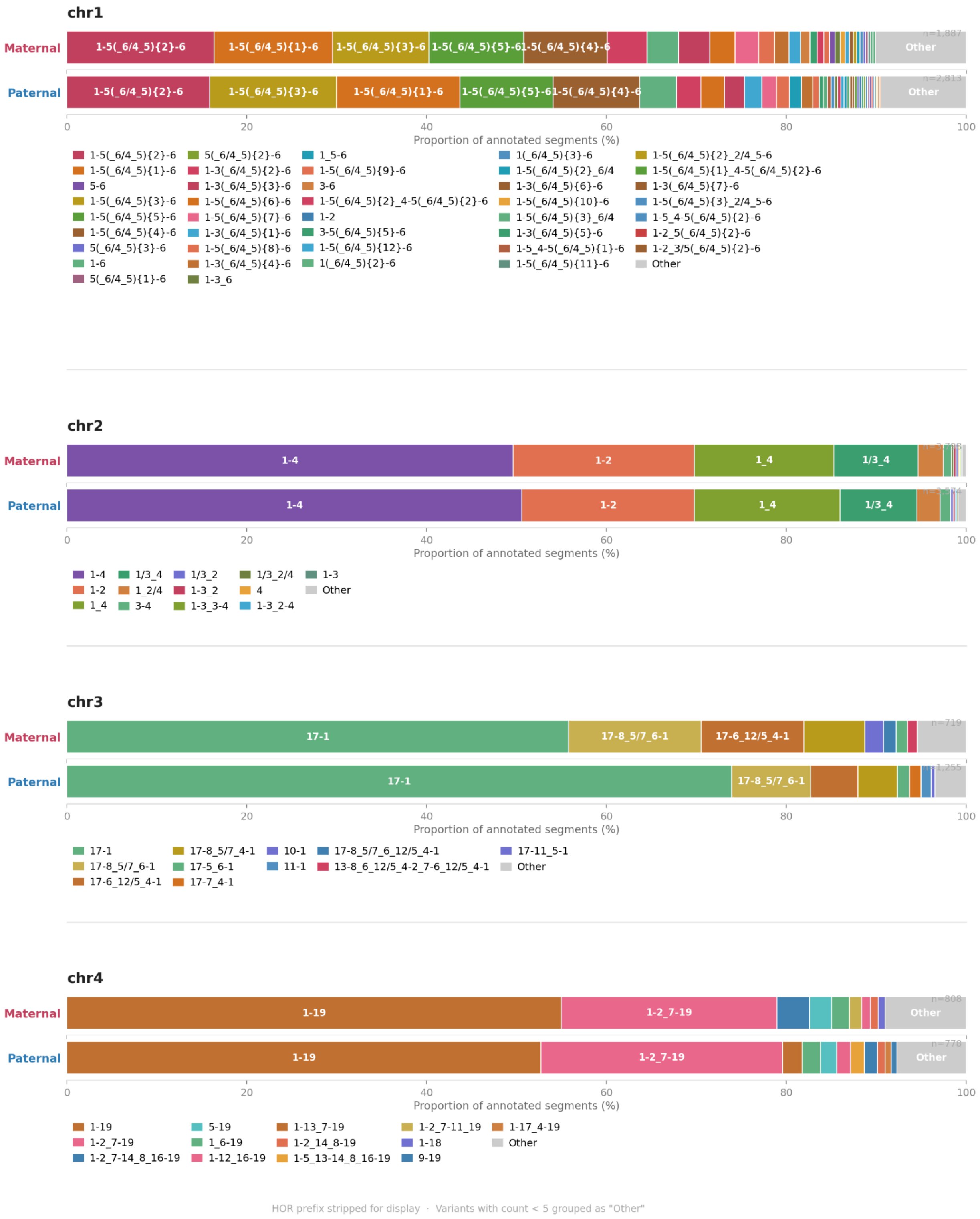
chr1
Maternal
Paternal
n=1,887
n=2,813
1-5(_6/4_5){2}-6
1-5(_6/4_5){1}-6
1-5(_6/4_5){3}-6
1-5(_6/4_5){5}-6
1-5(_6/4_5){4}-6
Other
0
20
40
60
80
100
Proportion of annotated segments (%)
1-5(_6/4_5){2}-6
1-5(_6/4_5){1}-6
5-6
1-5(_6/4_5){3}-6
1-5(_6/4_5){5}-6
1-5(_6/4_5){4}-6
5(_6/4_5){3}-6
1-6
5(_6/4_5){1}-6
5(_6/4_5){2}-6
1-3(_6/4_5){2}-6
1-3(_6/4_5){3}-6
1-5(_6/4_5){6}-6
1-5(_6/4_5){7}-6
1-3(_6/4_5){1}-6
1-5(_6/4_5){8}-6
1-3(_6/4_5){4}-6
1-3_6
1_5-6
1-5(_6/4_5){9}-6
3-6
1-5(_6/4_5){2}_4-5(_6/4_5){2}-6
1-2
3-5(_6/4_5){5}-6
1-5(_6/4_5){12}-6
1(_6/4_5){2}-6
1(_6/4_5){3}-6
1-5(_6/4_5){2}_6/4
1-3(_6/4_5){6}-6
1-5(_6/4_5){10}-6
1-5(_6/4_5){3}_6/4
1-3(_6/4_5){5}-6
1-5_4-5(_6/4_5){1}-6
1-5(_6/4_5){11}-6
1-5(_6/4_5){2}_2/4_5-6
1-5(_6/4_5){1}_4-5(_6/4_5){2}-6
1-3(_6/4_5){7}-6
1-5(_6/4_5){3}_2/4_5-6
1-5_4-5(_6/4_5){2}-6
1-2_5(_6/4_5){2}-6
1-2_3/5(_6/4_5){2}-6
Other
chr2
Maternal
Paternal
1-4
1-2
1_4
1/3_4
0
20
40
60
80
100
Proportion of annotated segments (%)
1-4
1-2
1_4
1/3_4
1_2/4
3-4
1/3_2
1-3_2
1-3_3-4
1/3_2/4
4
1-3_2-4
1-3
Other
chr3
Maternal
Paternal
n=719
1,255
17-1
17-8_5/7_6-1
17-6_12/5_4-1
0
20
40
60
80
100
Proportion of annotated segments (%)
17-1
17-8_5/7_6-1
17-6_12/5_4-1
17-8_5/7_4-1
17-5_6-1
17-7_4-1
10-1
11-1
17-8_5/7_6_12/5_4-1
13-8_6_12/5_4-2_7-6_12/5_4-1
17-11_5-1
Other
chr4
Maternal
Paternal
n=808
n=778
1-19
1-2_7-19
Other
0
20
40
60
80
100
Proportion of annotated segments (%)
1-19
1-2_7-19
1-2_7-14_8_16-19
5-19
1_6-19
1-12_16-19
1-13_7-19
1-2_14_8-19
1-5_13-14_8_16-19
1-2_7-11_19
1-18
9-19
1-17_4-19
Other
HOR prefix stripped for display · Variants with count < 5 grouped as "Other"

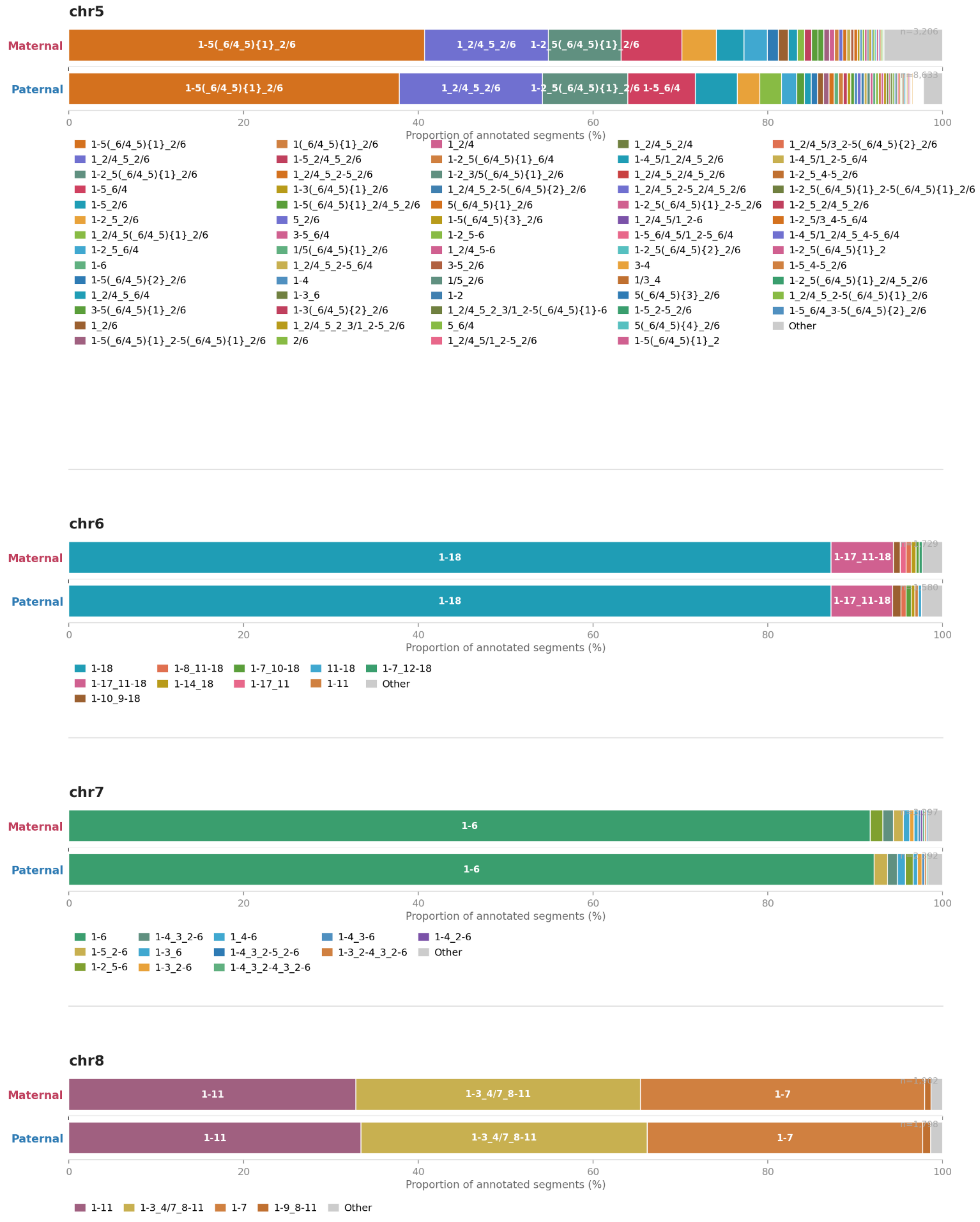


HOR prefix stripped for display · Variants with count < 5 grouped as "Other"

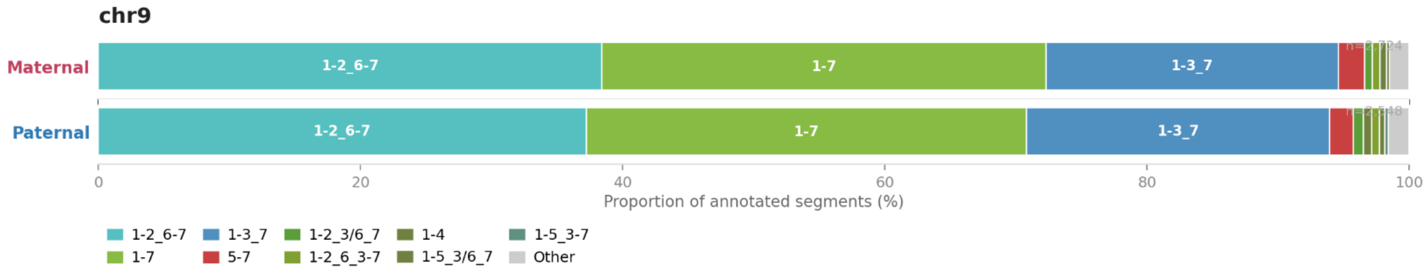


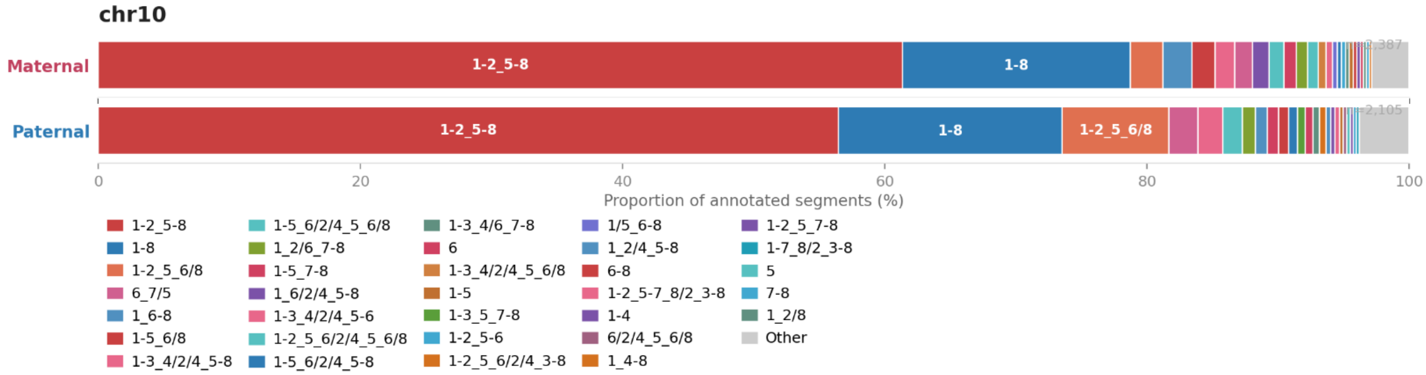


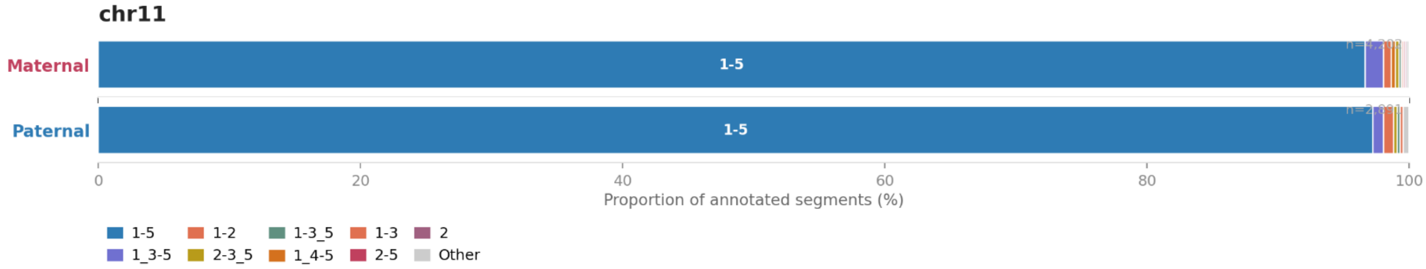


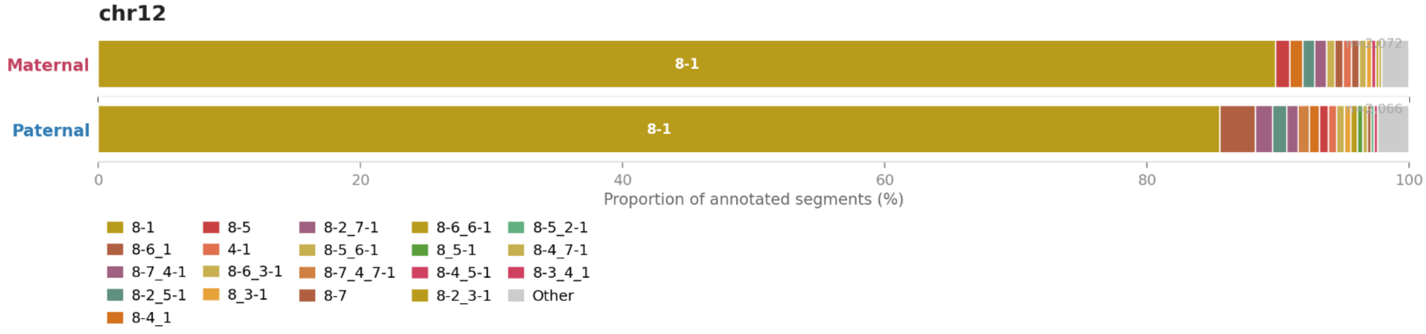


HOR prefix stripped for display · Variants with count < 5 grouped as "Other"

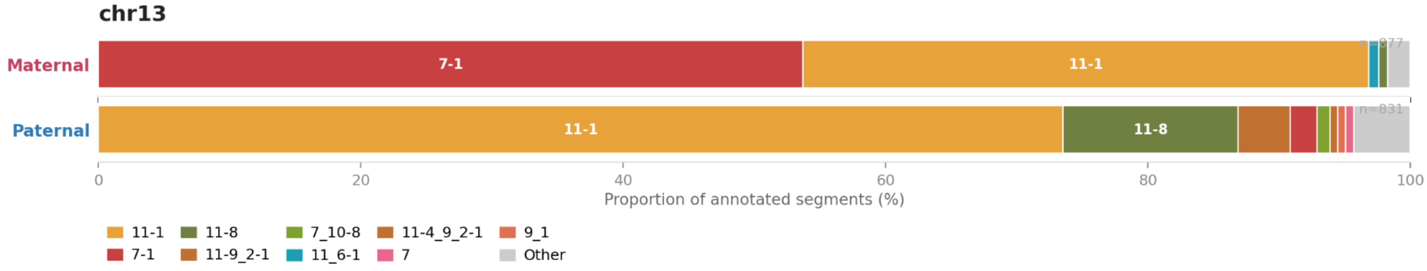


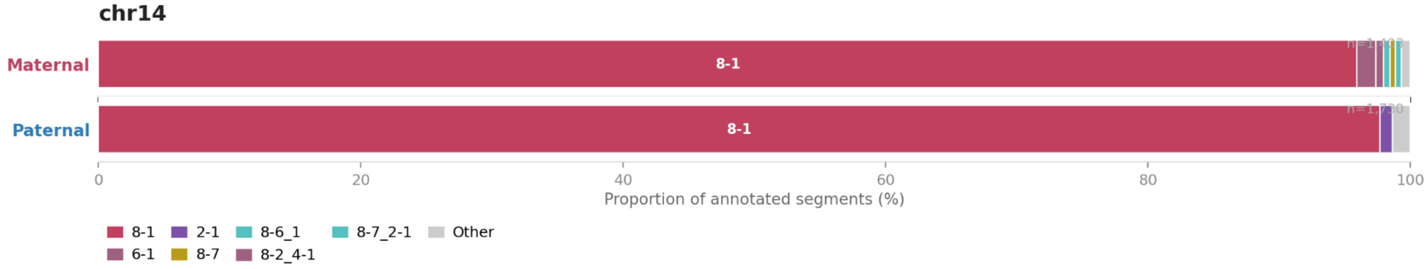


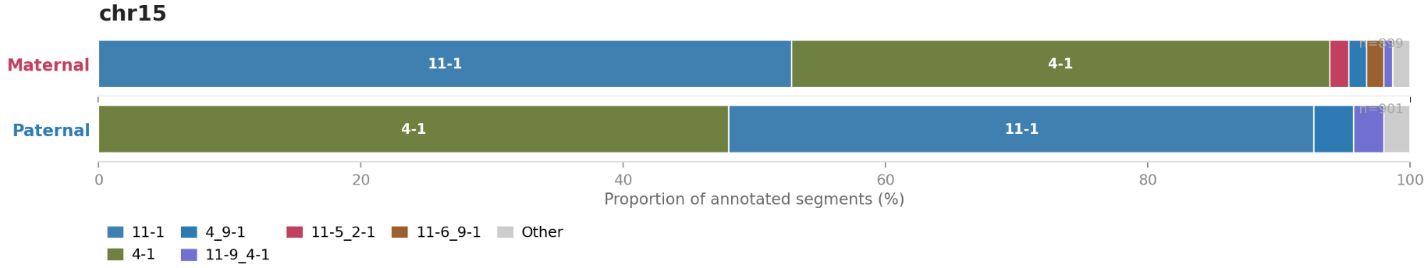


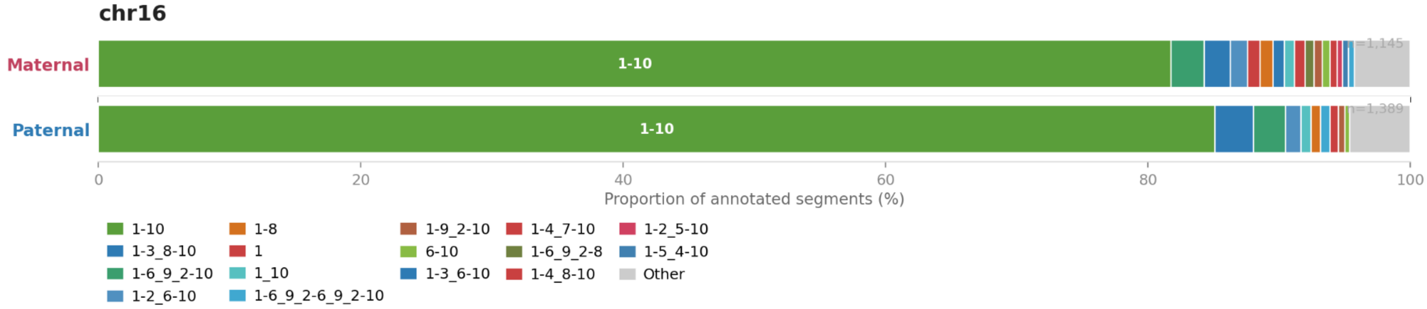


HOR prefix stripped for display · Variants with count < 5 grouped as "Other"

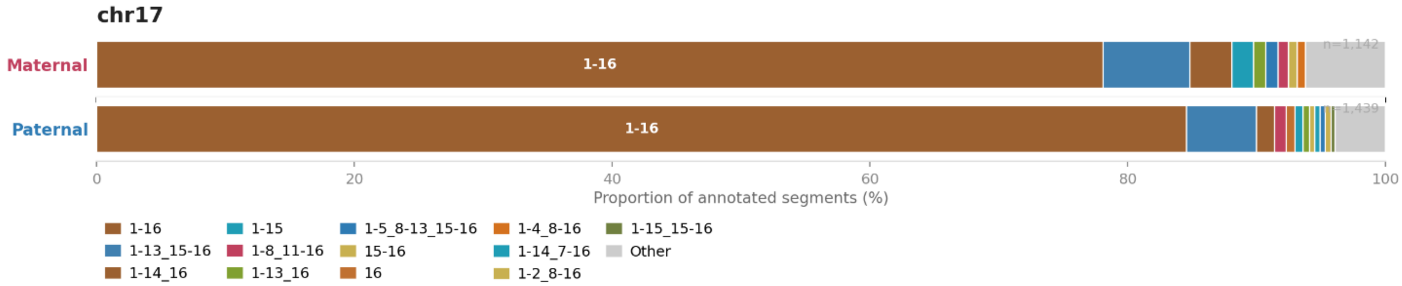


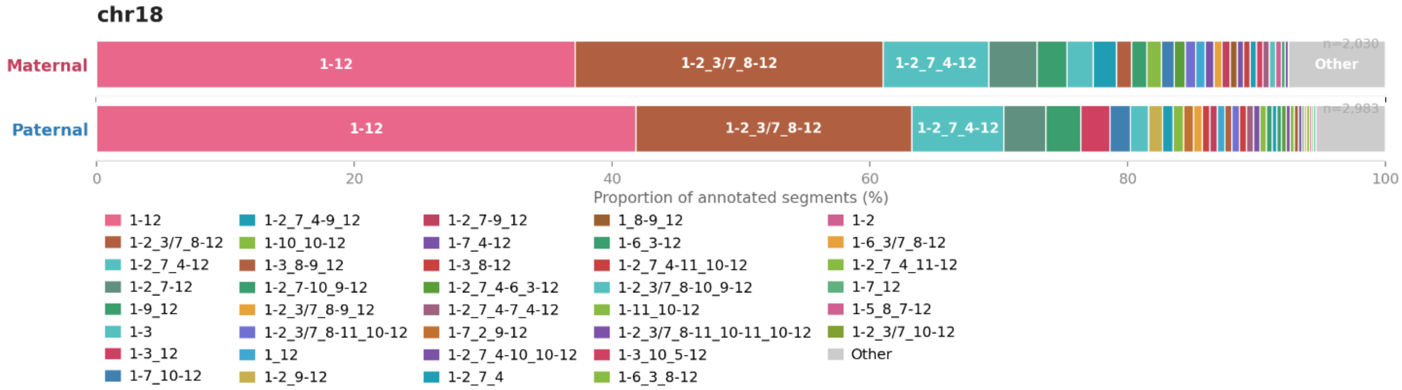


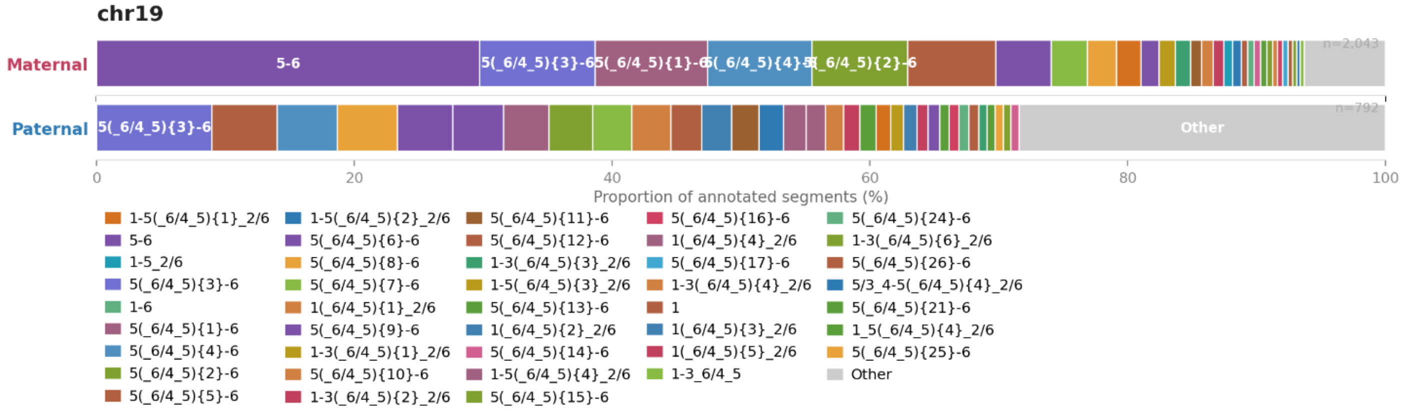


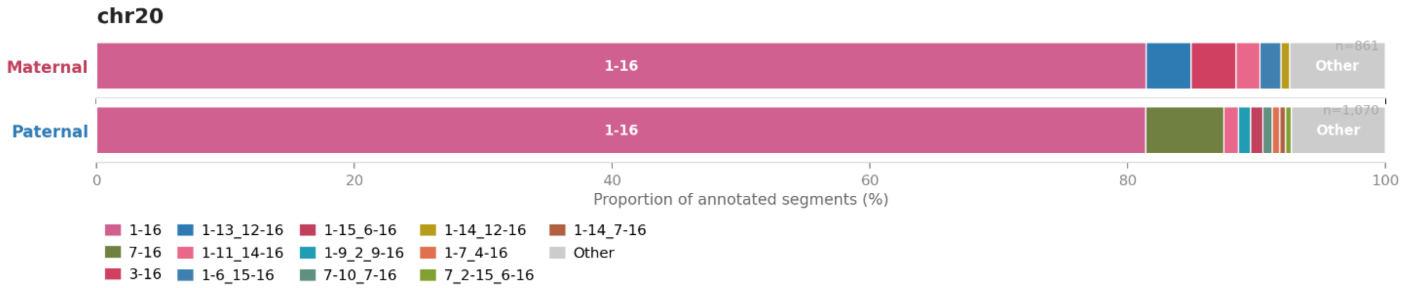




HOR prefix stripped for display · Variants with count < 5 grouped as "Other"

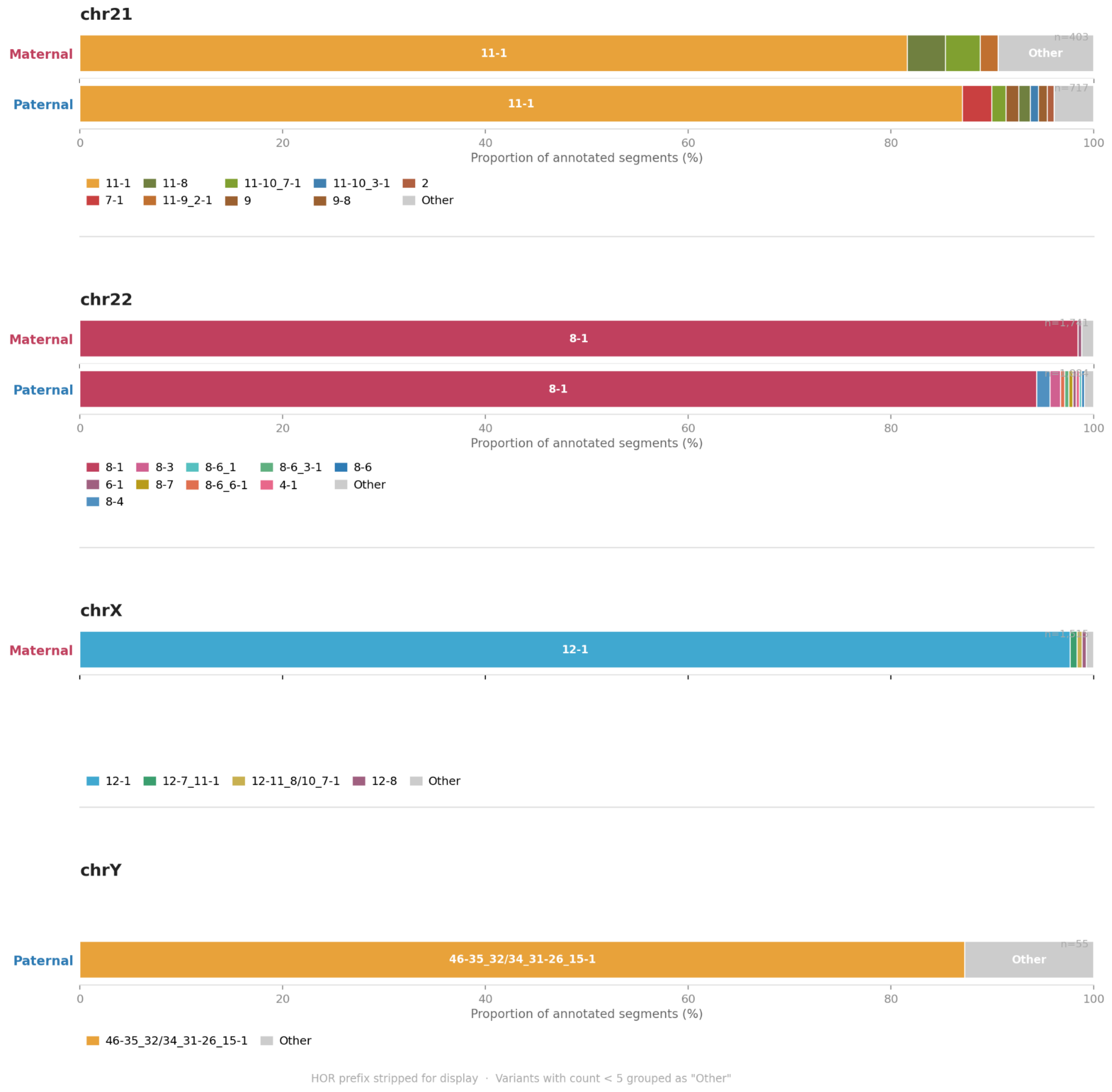


**Supplementary Figure 6: HOR Structural Variant (StV) profiles**
Stacked horizontal bar charts showing the proportional distribution of alpha-satellite HOR structural variants for each chromosome, with maternal (top) and paternal (bottom) haplotypes shown separately. Each colored segment represents a unique HOR variant, labeled by its structural suffix (common chromosome-specific HOR prefix removed for clarity). Segment width reflects the proportion of total annotated array segments occupied by each variant. Variants with fewer than 5 are grouped into "Other" (grey). The total number of annotated segments per haplotype (n) is indicated at the right of each bar.

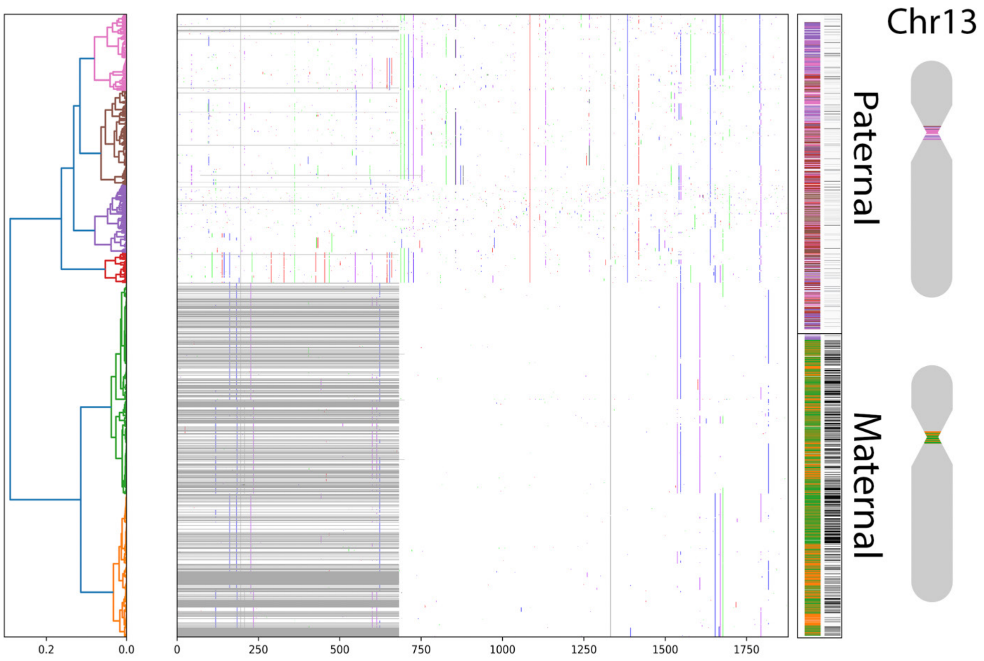


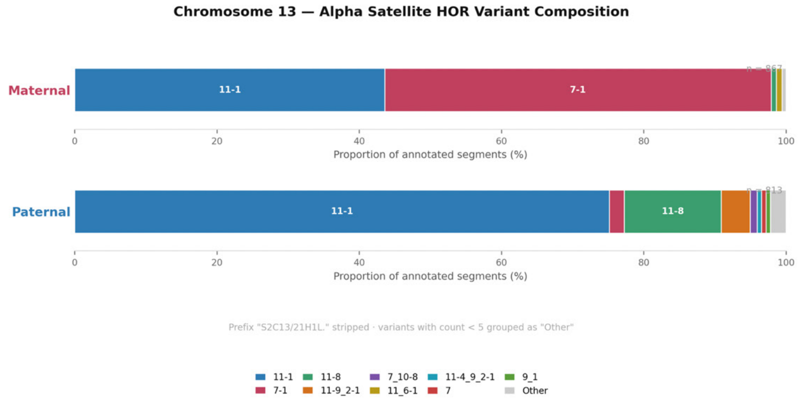


**Supplementary Figure 7: Chromosome 13 Structural Variant (StV) and HORhap analysis**

Ward HORhap clustering for chromosome 13 in HG002. Sequences of the two most frequent HOR structural variants (StVs) (the full-length S2C13/21H1L.11-1 and S2C13/21H1L.7-1 with a deletion of four monomers) from maternal and paternal arrays were aligned together. Only positions that differ from the consensus are colored (center). HORs in the alignment are ordered according to the Ward dendrogram (left), which is split into six colored clades (HORhaps). On the right, paternal and maternal centromeres (shown one above the other) are displayed as HORhap tracks with the corresponding colors (left) and tracks showing the positions of 7-mer StVs (right). On the far right, the positions of the centromeres within the entire chromosomes are indicated. The specificity of HORhaps to homologs is evident: the paternal homolog consists only of pink, brown, purple, and red HORhaps, corresponding to the upper major branch of the dendrogram, whereas the maternal homolog consists almost entirely of green and orange HORhaps, corresponding to the lower major branch, with only the very tip of the p-arm (upper end) represented by HORhaps from the upper major branch.

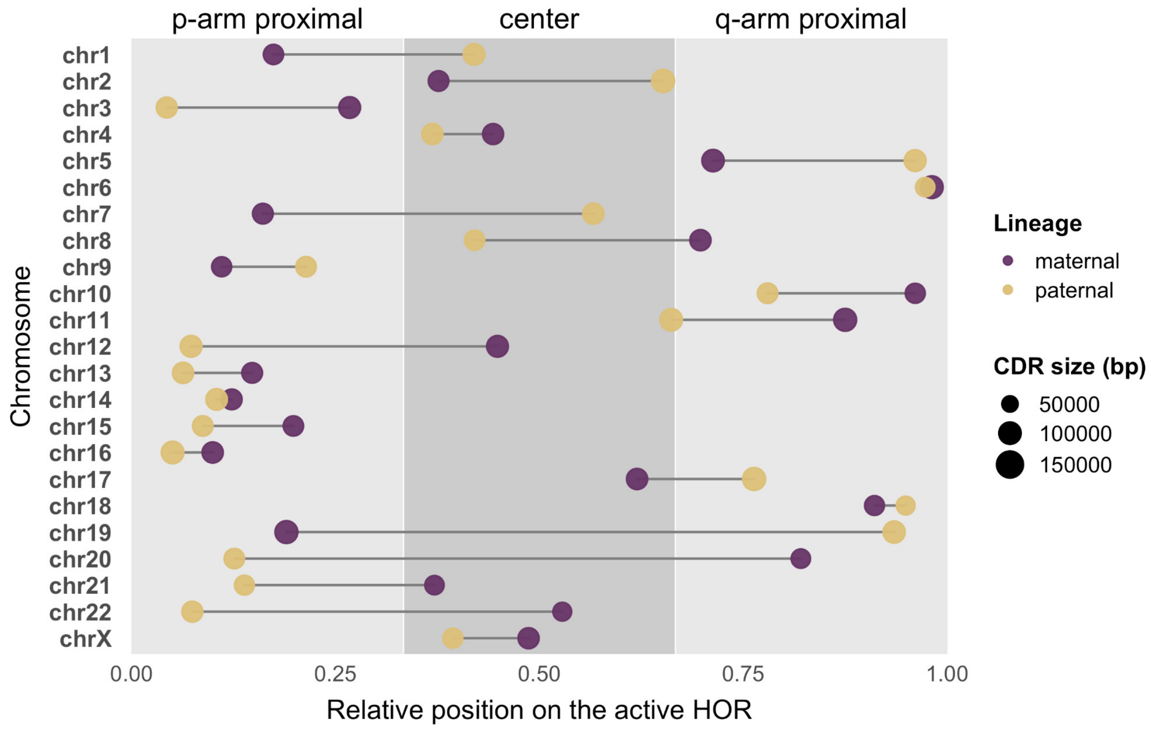


**Supplementary Figure 8: p-arm proximal position of CDR**
The relative position and size of CDRs within the active higher-order repeat array for each chromosome. The relative location of the CDR midpoint within the active array (x-axis; split equally between p-arm proximal, center, and q-arm proximal location) is noted for each chromosome (y-axis). The size of the dot is relative to the size of the CDR, here defined as the sum of all the subCDRs. Finally, the maternal and paternal CDRs for each homologous chromosome (including X and Y) are connected by a horizontal line. The p-arm proximal location is the most frequent out of the three possibilities.

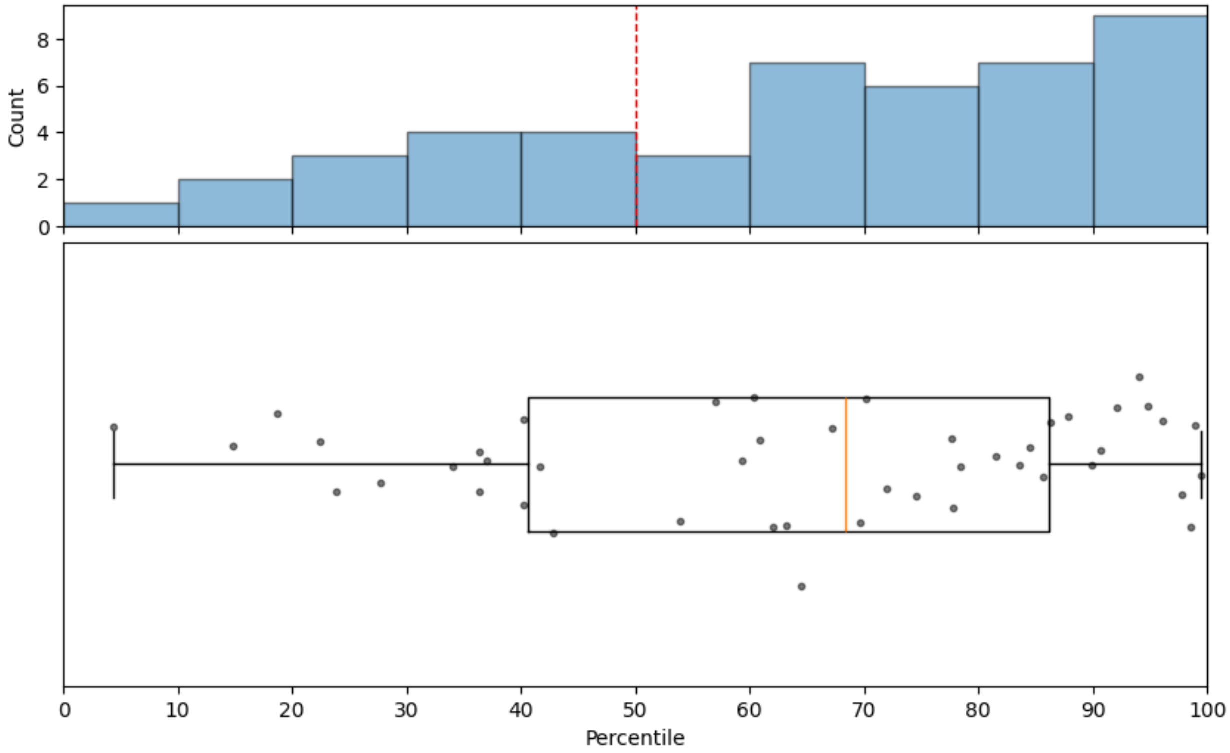


**Supplementary Figure 9: CDR association with homogenized regions**
CDR local identity as a percentile of the active alpha array distribution. For each chromosomal CDR, defined as the span between the two outermost subCDRs on a given haplotype, mean local identity was calculated across overlapping bins. To assess relative self-similarity, each CDR's mean local identity was compared to the background distribution of local identities across the whole active array. Each chromosome was assigned a percentile score reflecting where its CDR mean local identity falls within the active array distribution. The bottom panel shows per-chromosome CDR percentiles as a scatter-boxplot, with the top panel displaying the corresponding distribution as a histogram.

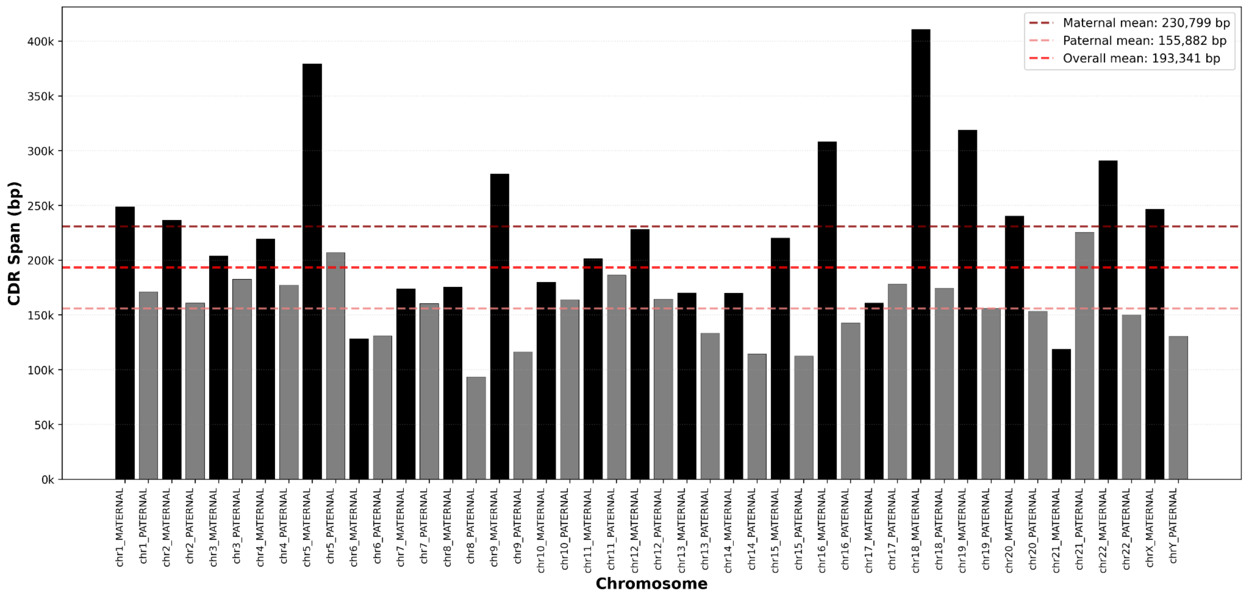


**Supplementary Figure 10: Total CDR span per haplotype in early passage LCL (HG002)**
Stacked bar plots showing the aggregate CDR span (kb) per haplotype across all 46 phased assemblies (chr1–chrY), as called by hmmCDR from DiMeLo-seq CENP-A data in the young passage LCL. Each bar represents a single haplotype (M, maternal; P, paternal), with individual stacked segments corresponding to discrete CDR subdomains; segment shading cycles from light to dark grey to distinguish adjacent subdomains. Horizontal reference lines indicate the mean aggregate CDR span across all maternal haplotypes (dark red dotted), all paternal haplotypes (light red dotted), and all haplotypes combined (red dashed). Mean aggregate CDR span was 87.4 kb (maternal), 89.4 kb (paternal), and 88.4 kb genome-wide across 254 total CDR domains.

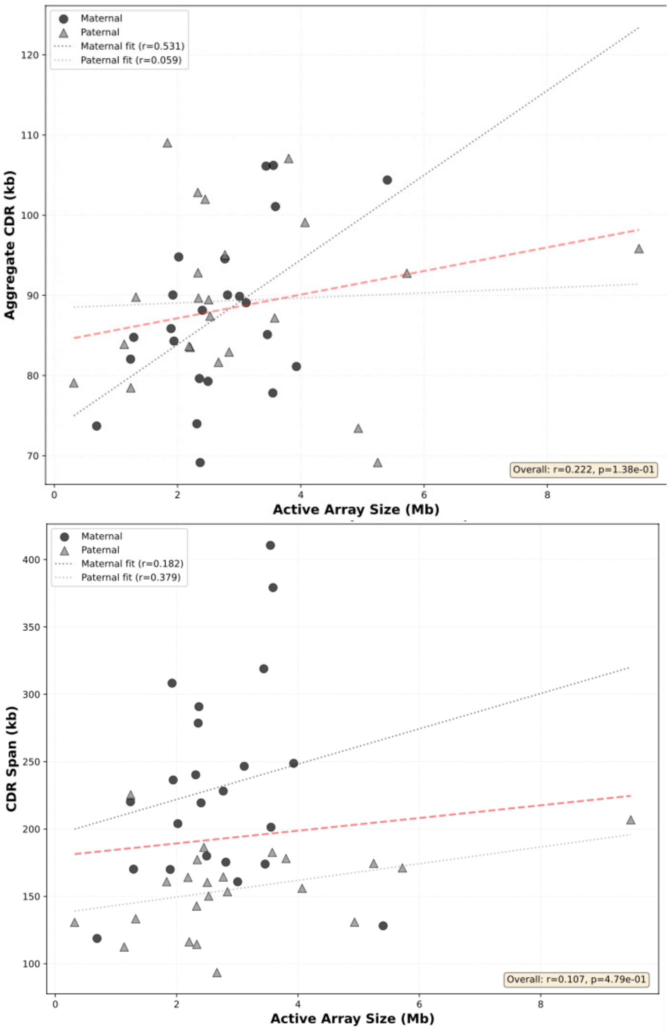


**Supplementary Figure 11: CDR (subCDR and span) vs active array size** Scatter plots showing the relationship between CDR length (x-axis) and total active αSat HOR array length (y-axis, Mb) for all chromosomes. Maternal haplotypes are shown as dark grey circles and paternal haplotypes as light grey squares; dashed lines indicate linear fits with associated Pearson r-values. (Top) Aggregated sub-CDR length, calculated as the sum of individual hypomethylated sub-CDR segments, excluding intervening regions between domains. The aggregated sub-CDR length is highly consistent genome-wide (mean ~90 kb; range: 69–109 kb) and shows no significant correlation with array size (maternal r = 0.059; paternal r = 0.059). (Bottom) Total CDR span, measured from the start of the first sub-CDR to the end of the last sub-CDR, including intervening non-hypomethylated sequence. Total span is more variable and shows a modest positive correlation with array size (maternal r = 0.182; paternal r = 0.379). Together, these plots demonstrate that while CDR span varies across haplotypes and chromosomes, the aggregate length of the hypomethylated sub-CDR core is remarkably stable and decoupled from the size of the underlying active HOR array.

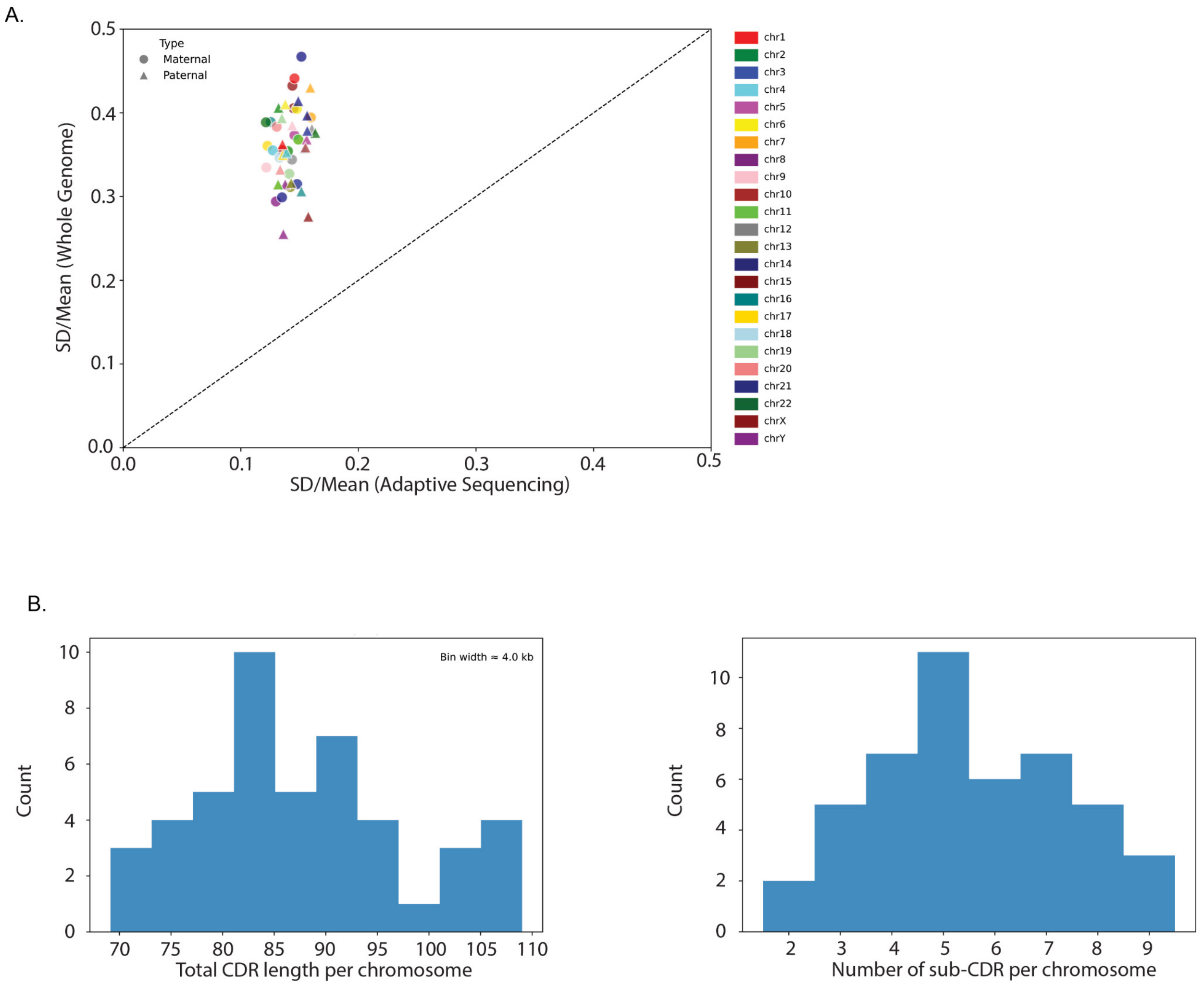


**Supplementary Figure 12: Coverage estimates of adaptive sampling**

Adaptive sampling achieves uniform coverage across centromeric arrays. (A**)** Scatter plot comparing coverage uniformity between whole genome sequencing and adaptive sampling across centromeric regions. The coefficient of variation (SD/mean) for whole genome sequencing (y-axis) is broadly distributed, whereas the coefficient of variation for adaptive sampling (x-axis) is consistently low, demonstrating that adaptive sampling yields substantially more uniform coverage across centromeric arrays relative to whole genome sequencing. (B**)** Overlaid density histograms of mean sequencing coverage across 5 kb genomic bins for adaptive sampling (+AS) and whole genome sequencing (−AS). Coverage values were calculated by aggregating per-base depth into non-overlapping 5 kb windows. The +AS distribution is narrow and concentrated around the mean, reflecting highly uniform coverage across centromeric regions, whereas the −AS distribution is broadly dispersed, indicating greater bin-to-bin coverage variability in whole genome sequencing.

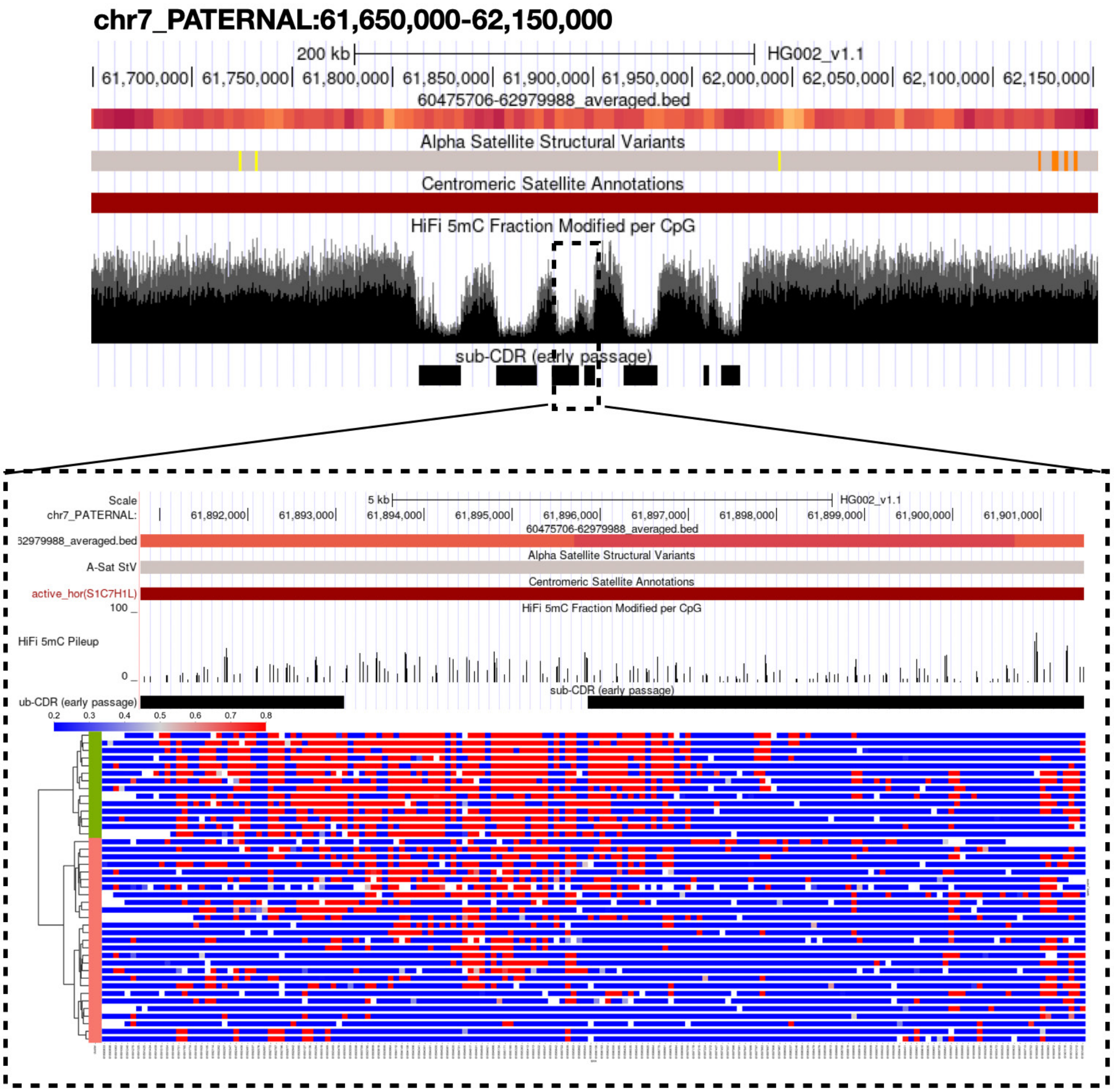


**Supplementary Figure 13: Clustering methylation data into groups** Clustered read matrices showing CpG methylation (5mC) signal across individual reads at the CDR edge in chromosome 7 paternal. Each row represents a single sequencing read, and each column a CpG position within the region. Reads were clustered using HDBSCAN based on their per-CpG 5mC probability profiles, with only reads covering at least 95% of the locus retained (windowFrac = 0.95). Distinct clusters reflect discrete molecule-level methylation states, supporting the interpretation that epigenetic heterogeneity at these regions is a genuine biological signal.

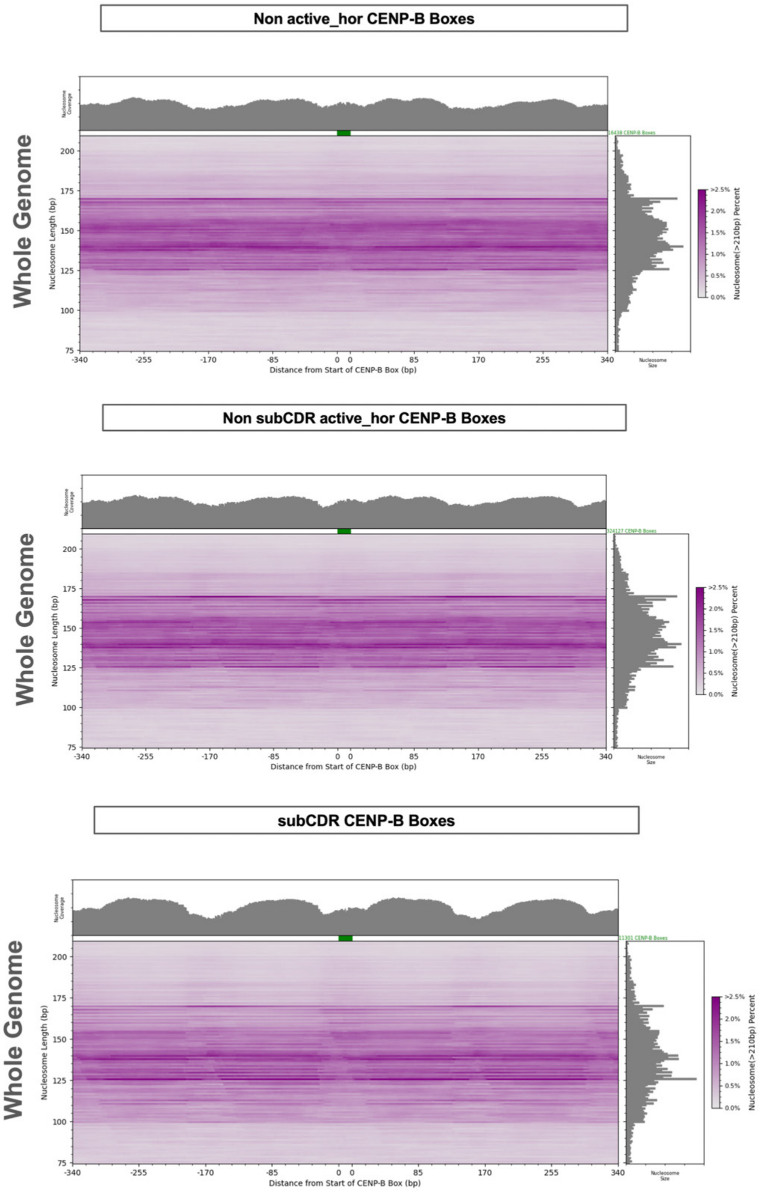


**Supplementary Figure 14: Fiber-seq nucleosome organization relative to CENP-B motifs**
Fiber-seq[3] comparative nucleosome organization relative to centromere protein B (CENP-B) 17-bp motifs[4] in CDR, Active, and Inactive Alpha Arrays. Nucleosome positions with respect to reference coordinates were extracted using fibertools [5] extract. For each of the three regions (within subCDRs, non-CDR active alpha array, and inactive alpha array), a matrix was filled with the position of the nearest CENP-B motif and nucleosome-size. The matrix was then column-normalized to estimate the percentage of nucleosomes at specific sizes per distance from a CENP-B motif. Darker heatmap regions indicate a higher density of nucleosomes at a given size and position. Count totals are projected onto each axis as marginal histograms, summarizing the row and column distributions of the matrix. The top histogram summarizes nucleosome positional distribution relative to the CENP-B box motif, while the right histogram shows the overall nucleosome size distribution within each of the three regions observed.

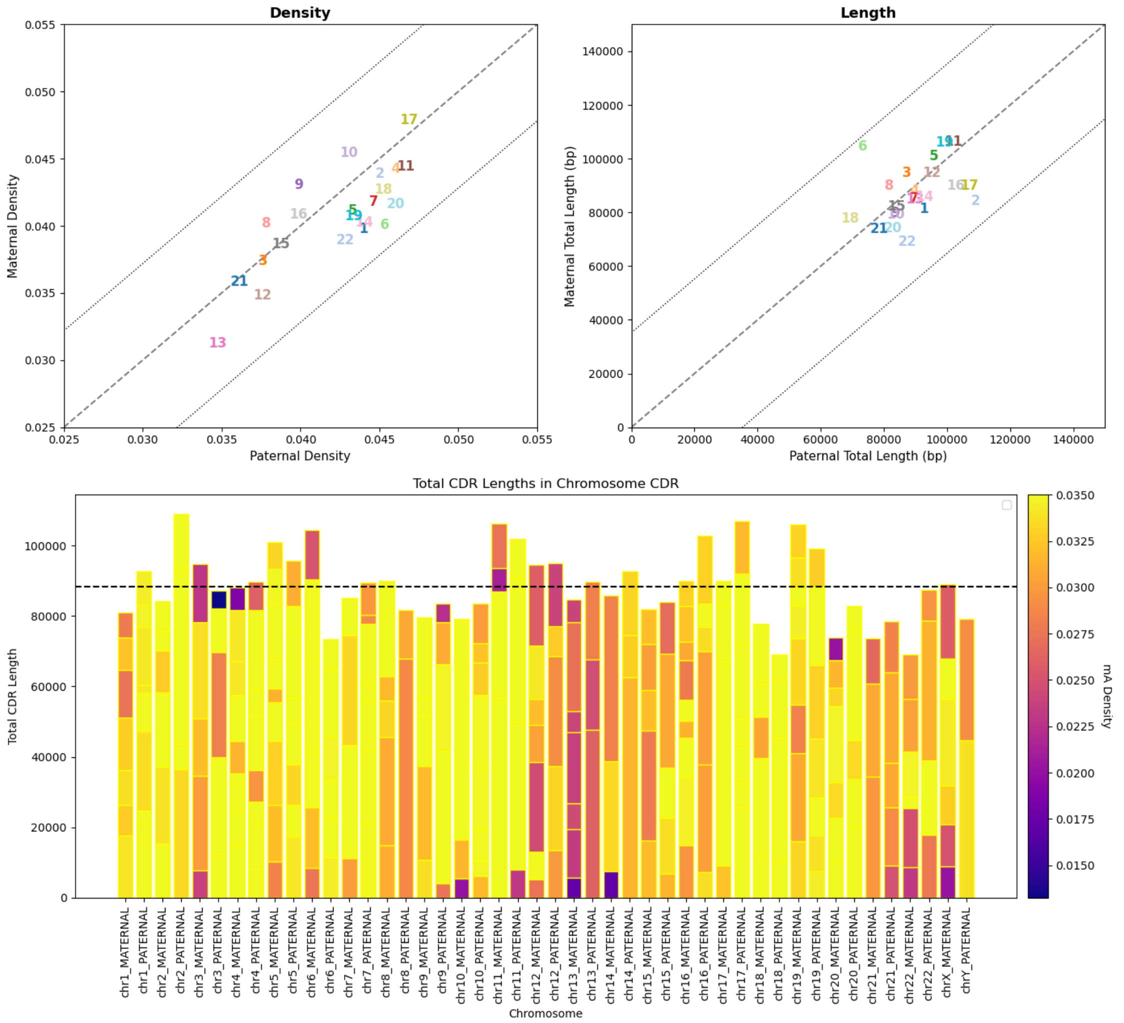


**Supplementary Figure 15: CENP-C DiMeLo-seq 6mA density (early passage)**

Centromere protein C (CENP-C) chromatin dosage is balanced between homologous centromeres in early-passage LCLs. (A) Scatter plots comparing paternal (x-axis) and maternal (y-axis) CENP-C DiMeLo-seq[6] 6mA signal density (left) and total sub-CDR length (right) across all chromosomes in early-passage HG002 LCLs. Each chromosome is labeled and color-coded. The dashed diagonal line represents the identity line; dotted lines indicate ±3 SD outlier thresholds. Chromosomes deviating beyond these thresholds represent haplotype-specific departures from dosage balance. (B) Stacked bar plots showing total CDR length per chromosome for maternal and paternal haplotypes for CENP-C DiMeLo-seq. Each bar is subdivided into individual sub-CDR segments colored by 6mA signal density using a plasma colormap. The dashed horizontal line indicates the mean total CDR length across all homologs. A color bar indicates the range of 6mA density values. Maternal and paternal haplotypes are displayed as adjacent pairs for each chromosome, ordered from chr1 to chrY.

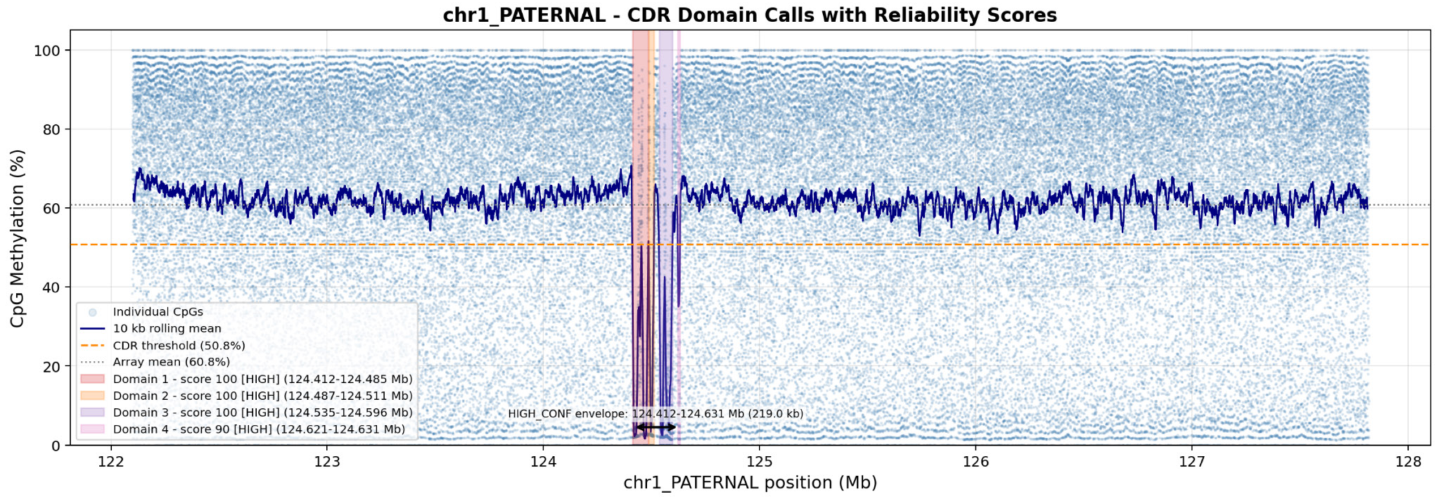


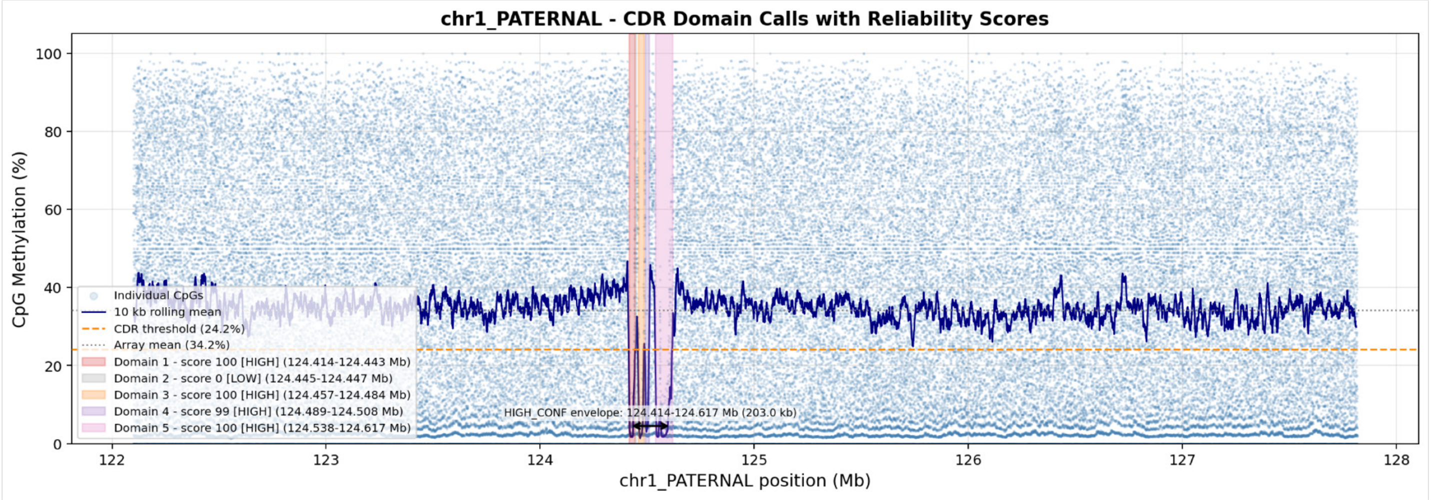


**Supplementary Figure 16: Methylation frequency across active arrays in early vs late passage**
Methylation plot pileup (specifically BEDmethyl/bedGraph formats) summarizes per-base DNA methylation frequencies (e.g., 5mC) from CENP-A DiMeLo-seq data across the active array of chromosome 1_PATERNAL early passage (EP) (top) and late passage (LP) (bottom). Individual windows of CpG are provided in light blue, with a 10kb rolling mean in dark blue. The dotted line indicates the array mean (as shown for chromosome 1_PATERNAL, EP=60.8%, and LP=34.2%). CDRs are highlighted as individual domains that pass the threshold (10% difference from the array mean), and are shown to highlight that the CDRs remain in the same span, but have altered sub-CDR boundaries.

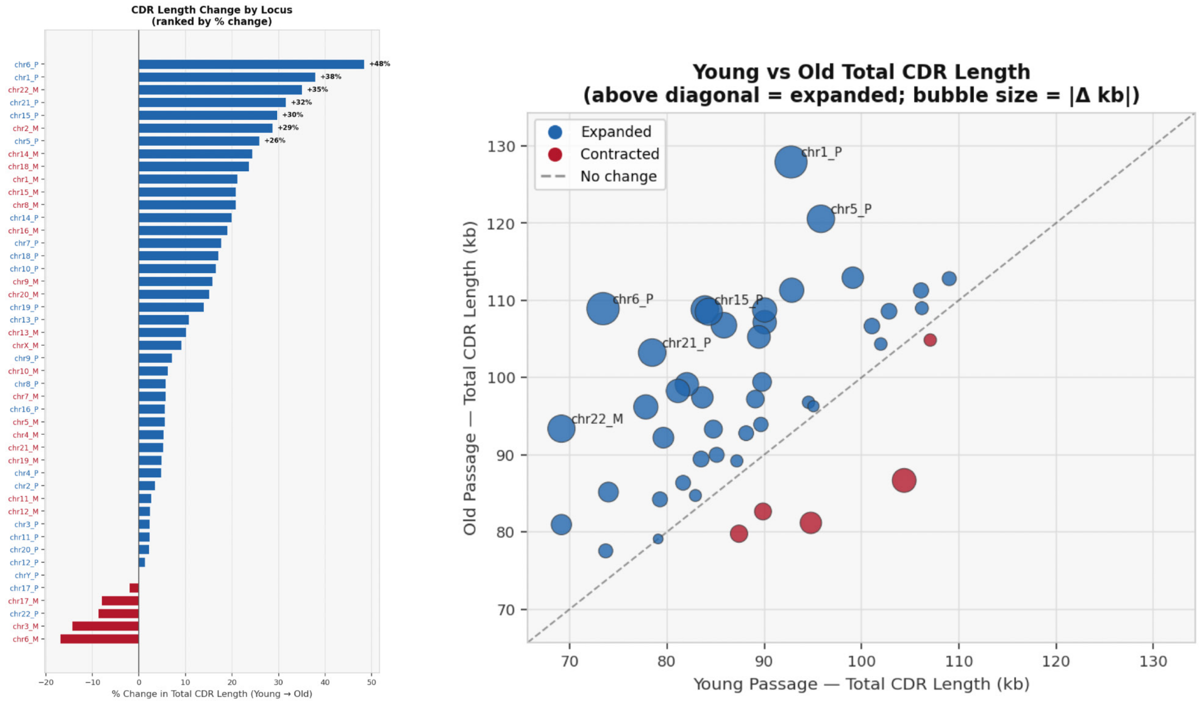


#### Supplementary Figure 17: Difference in sub-CDR structure  in early vs late passage

(Left) Ranked percent change in total sub-CDR aggregate length between early and late passage, shown for all active αSat arrays. Arrays are rank-ordered by magnitude of change, with increases shown in blue and decreases in red. Maternal and paternal haplotypes are labeled in red and blue, respectively. (Right) Scatter plot comparing total sub-CDR aggregate length (kb) between early (x-axis) and late (y-axis) passage. Each point represents a single array, with circle size proportional to the absolute magnitude of change between passages. The dashed line indicates the expectation under no change.

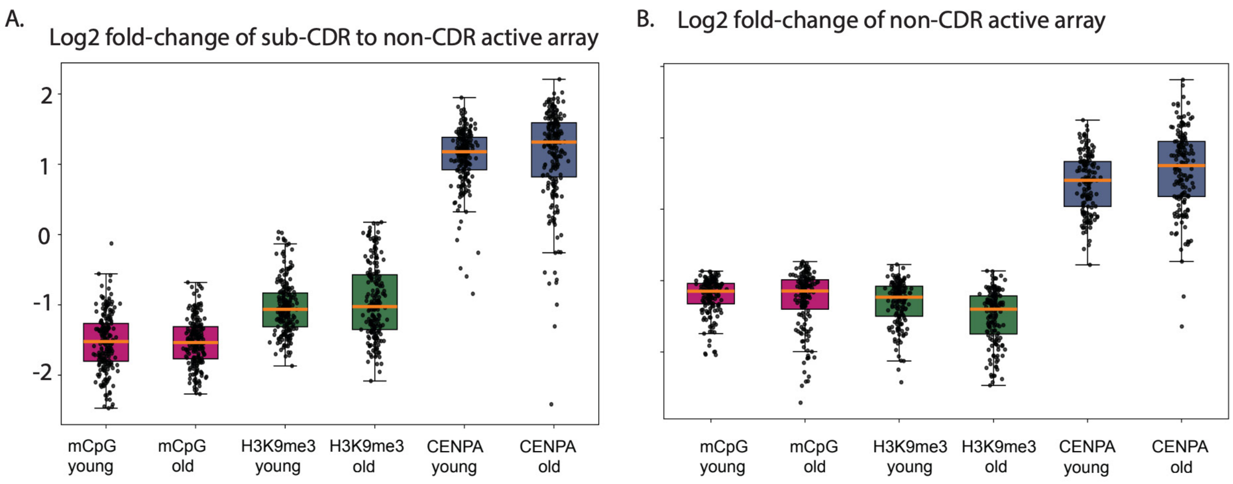


**Supplementary Figure 18: Enrichment within sub-CDRs across early- and late-passage**

Epigenetic marks within general CDRs are stable between early- and late-passage LCLs. Box plots showing log2 fold-change of mCpG, H3K9me3, and CENP-A signal densities within sub-CDRs in early- and late-passage HG002 LCLs, calculated relative to chromosome-specific non-CDR active array baselines. Across all three marks, early- and late-passage cells show highly similar enrichment profiles within sub-CDRs: mCpG is comparably depleted in both conditions (median log2 fold-change ~−1.5), H3K9me3 is similarly reduced, and CENP-A is equivalently enriched, indicating that the relative epigenetic identity of the overall CDR domains is robustly maintained despite extended culture.

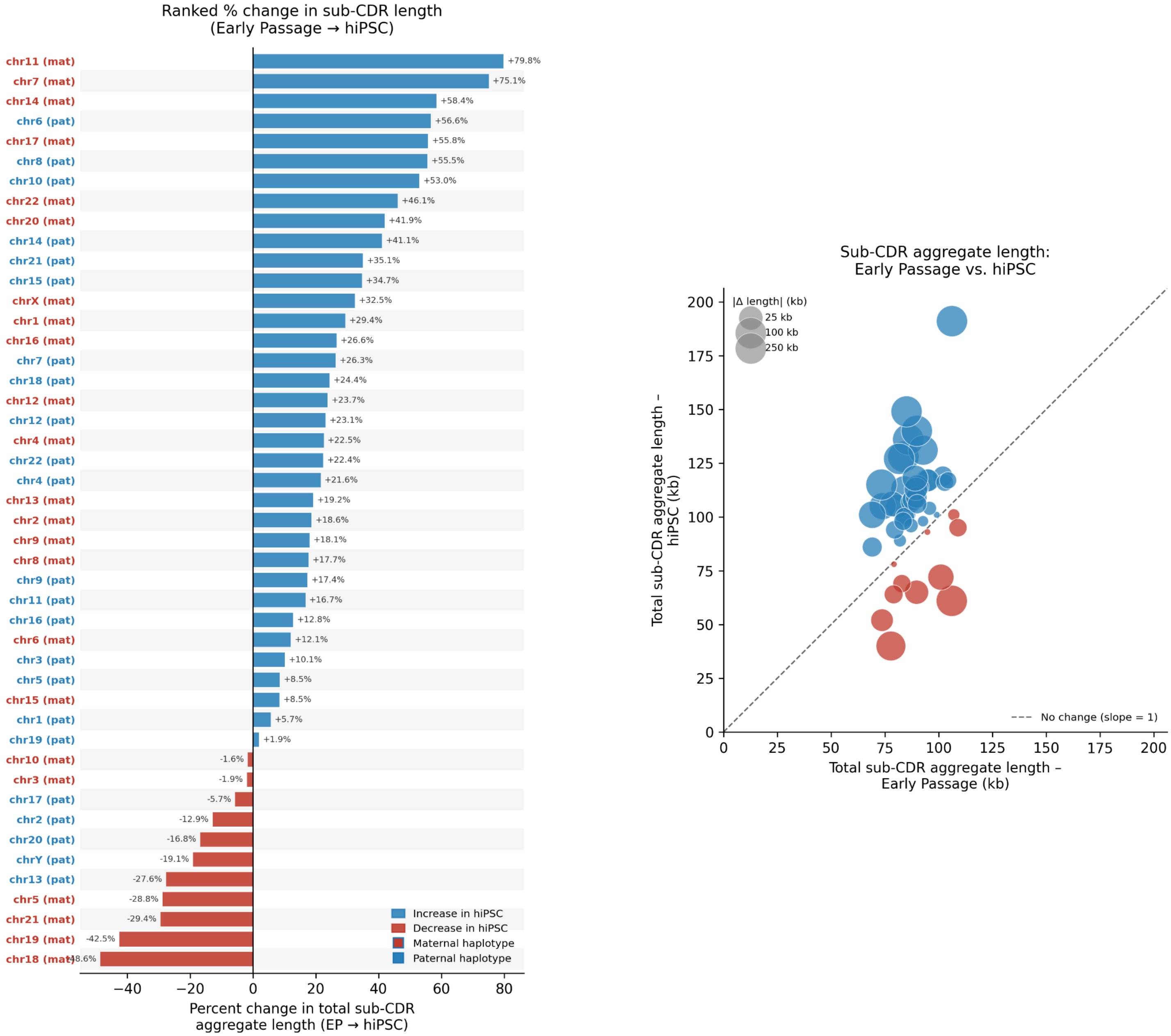


**Supplementary Figure 19: Comparison between HG002 early passage and hiPSC CDRs**

(Left) Ranked percent change in total sub-CDR aggregate length between early passage (EP) and hiPSC, shown for all active αSat arrays across 46 haplotypes. Arrays are rank-ordered by magnitude of change; increases in hiPSC relative to EP are shown in blue and decreases in red. Maternal and paternal haplotypes are indicated by red and blue labels, respectively. (Right) Scatter plot comparing total sub-CDR aggregate length (kb) between EP (x-axis) and hiPSC (y-axis). Each point represents a single array; circle size is proportional to the absolute magnitude of change between conditions. The dashed line indicates the expectation under no change.

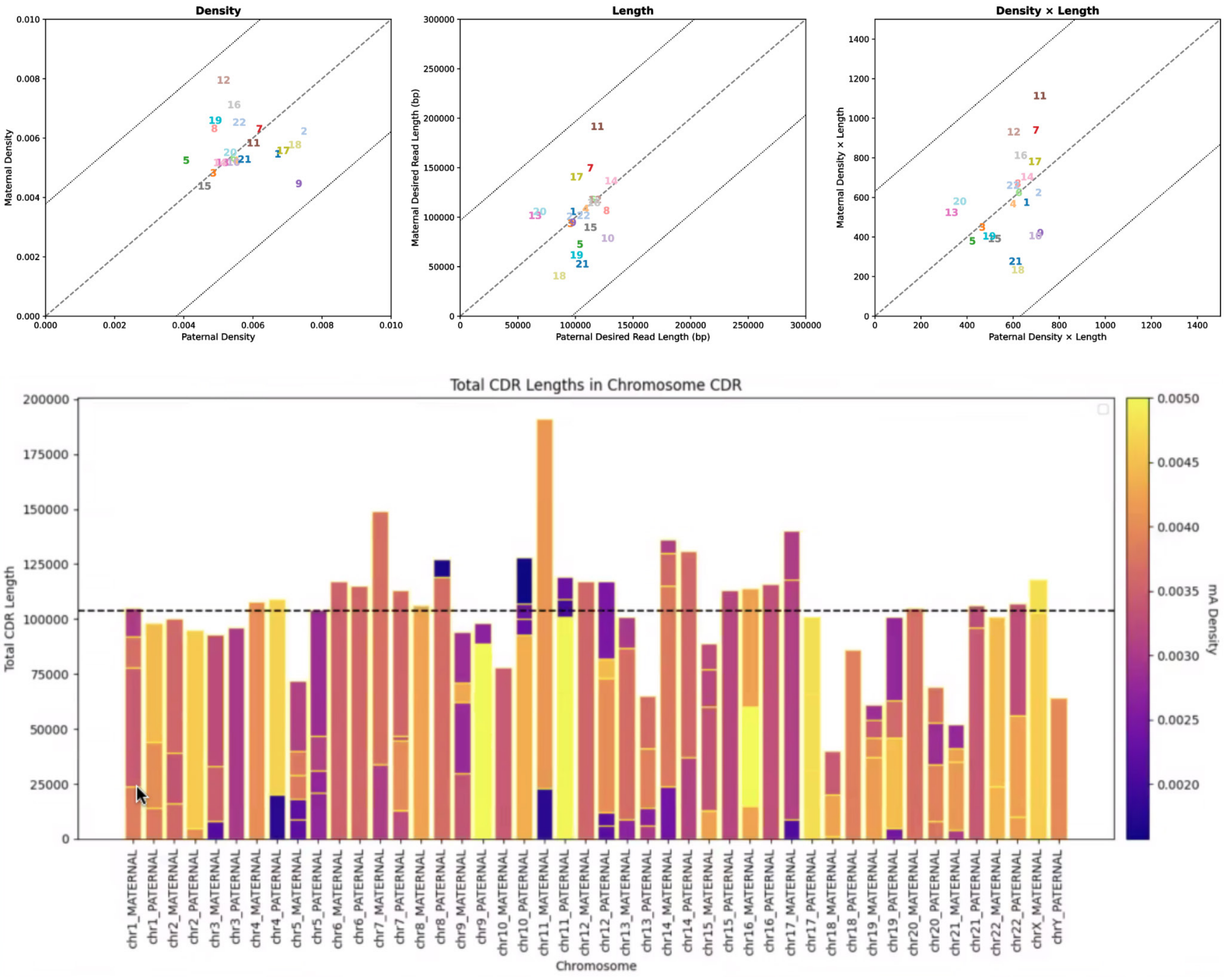


**Supplementary Figure 20: CENP-A DiMeLo-seq Density in hiPSCs**

CENP-A chromatin dosage is balanced between homologous centromeres in PBMC-derived iPSCs. (A) Scatter plots comparing paternal (x-axis) and maternal (y-axis) CENP-A DiMeLo-seq 6mA signal density (left) and total sub-CDR length (right) across all chromosomes in PBMC-derived iPSCs. Each chromosome is labeled and color-coded. The dashed diagonal line represents the identity line; dotted lines indicate ±3 SD outlier thresholds. (B) Stacked bar plots showing total CDR length per chromosome for maternal and paternal haplotypes for late-passaged CENP-A DiMeLo-seq. Each bar is subdivided into individual sub-CDR segments colored by 6mA signal density using a plasma colormap. The dashed horizontal line indicates the mean total CDR length across all homologs. A color bar indicates the range of 6mA density values. Maternal and paternal haplotypes are displayed as adjacent pairs for each chromosome, ordered from chr1 to chrY.

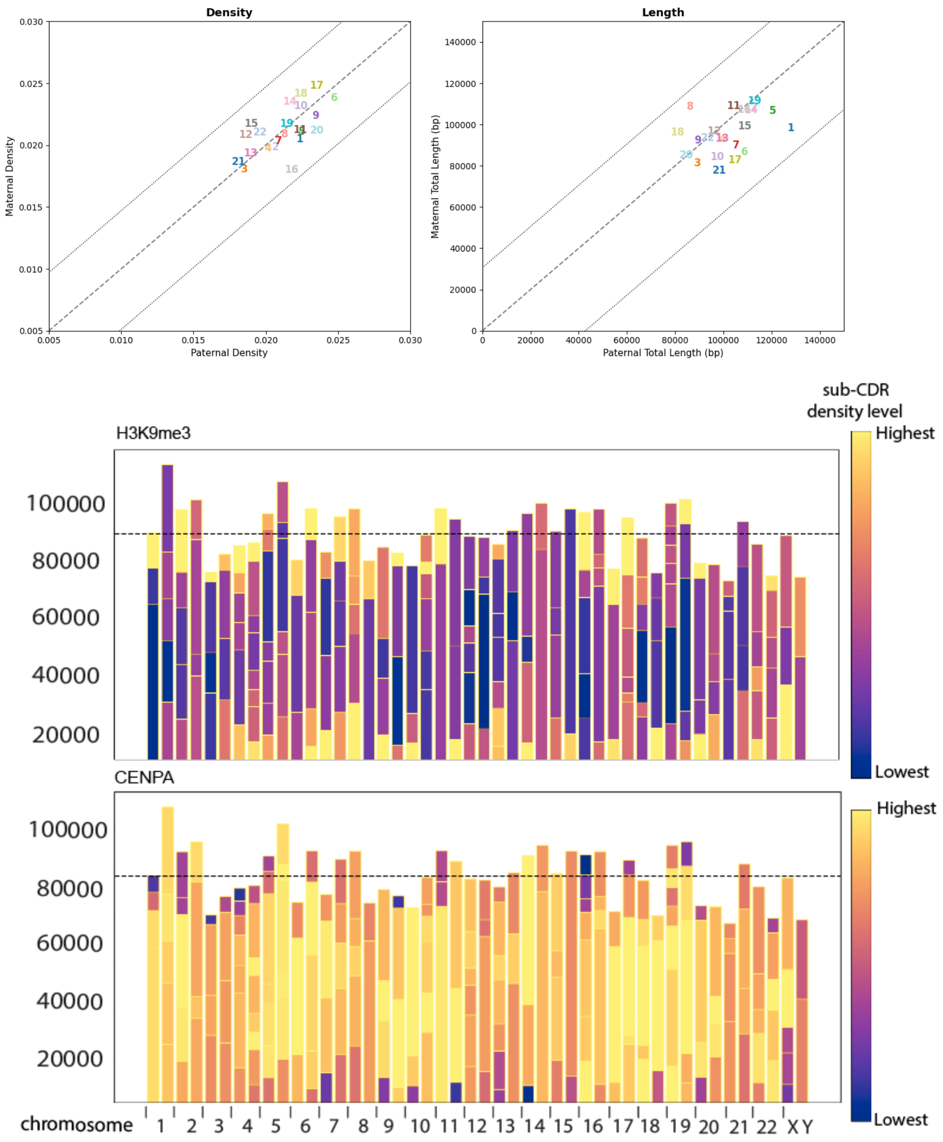


#### Supplementary Figure 21: CENP-A DiMeLo-seq density in late passage lines

CENP-A chromatin dosage is balanced between homologous centromeres in late-passage LCLs. (A) Scatter plots comparing paternal (x-axis) and maternal (y-axis) CENP-A DiMeLo-seq 6mA signal density (left) and total sub-CDR length (right) across all chromosomes in late-passage HG002 LCLs. Each chromosome is labeled and color-coded. The dashed diagonal line represents the identity line; dotted lines indicate ±3 SD outlier thresholds. Chromosomes deviating beyond these thresholds represent haplotype-specific departures from dosage balance. (B) Stacked bar plots showing total CDR length per chromosome for maternal and paternal haplotypes for late-passaged CENP-A DiMeLo-seq. Each bar is subdivided into individual sub-CDR segments colored by 6mA signal density using a plasma colormap. The dashed horizontal line indicates the mean total CDR length across all homologs. A color bar indicates the range of 6mA density values. Maternal and paternal haplotypes are displayed as adjacent pairs for each chromosome, ordered from chr1 to chrY.

### Supplemental Note 1: Higher-order repeat haplotype (HORhap) analysis

#### Overview

Within an active alpha-satellite higher-order repeat (HOR) array, individual HOR units accumulate subtle nucleotide variants. Groups of HORs that share specific variants are referred to as HOR haplotypes (HORhaps). Early classification approaches relied on manual inspection or k-means clustering [7]. Our current approach extends these methods by constructing multiple sequence alignments that include a few HOR structural variants (StV) (HOR units with various monomer compositions, e.g., full-length and deleted). Pairwise Hamming distances are then computed across alignment, and hierarchical clustering using Ward's method is applied to generate compact, interpretable trees, from which HORhap classes are defined by cutting the dendrogram at a selected depth. This pipeline was applied to HG002, as detailed below, and provided a scalable framework for broader population-scale analyses.

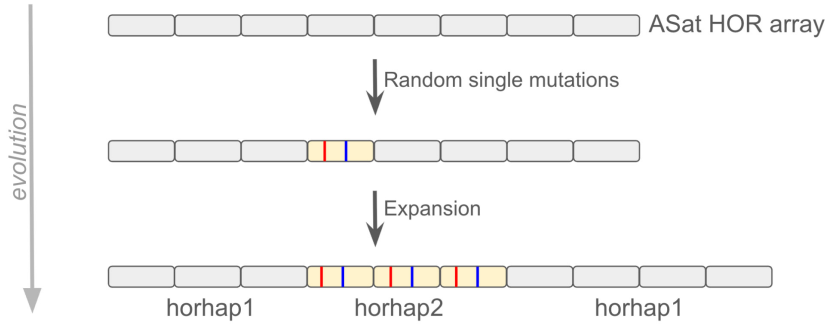


**SNote Figure 1.** Basic description of HORhaps, where HORs are expected to acquire mutations over time (shown for simplicity as happening at two sites (red and blue, in yellow HOR unit) at random). Through processes of non-homologous exchange (e.g., unequal cross-over) leading to local expansion and contraction of yellow HORs in the array. As a result, we can classify HOR positional mutations in the grey repeats vs the yellow repeats, to characterize their organization in the array (labeled "HORhap 1" vs. "HORhap2").

#### HORhap classification

To characterize haplotype-level variation within centromeric alpha-satellite HOR arrays, we developed a three-stage annotation pipeline: HOR monomer annotation, structural variant (StV) annotation, and HOR haplotype (HORhap) assignment. The first 2 steps were described in detail previously [7], so they are mentioned only briefly below.

In the first stage, HOR monomer annotations were generated using the HumAS-HMMER tool (https://github.com/fedorrik/HumAS-HMMER_for_AnVIL) with the AS-HORs HMM profile database, as described in Altemose et al., 2022 [7]. Each annotated record includes the HOR name (e.g., S2C9H1L) and the number of each monomer within the HOR unit (e.g., S2C9H1L.2).
In the second stage, StV annotations of active alpha-satellite arrays were generated from the monomer annotations using stv_tool (https://github.com/fedorrik/stv). This tool merges consecutive monomers into continuous HOR units, such that each annotated item corresponds to a full-length HOR (e.g., S2C9H1L.1-7) or a deletion variant (e.g., S2C9H1L.1-3_6-7, indicating deletion of monomers 4–5). More complex variants involving insertions are represented in an analogous manner.

In the third stage, StV annotations served as input to horhap_tool (https://github.com/fedorrik/horhap_tool). For each chromosome, both homologs were processed simultaneously. StV annotations were used to retrieve sequences of all the HOR copies for the two most frequent StVs. In cases where the second most frequent StV was less frequent than 1% of the total number of HORs, only the most frequent StV was used. In cases where the third most frequent StV was more frequent than 14% of the total number of HORs, the top 3 StVs were used. Chromosome 19 was excluded from the HORhap analysis due to the complex StV structure. The number of selected StVs is shown in the supplement table and SNote Table 1. Sequences of selected HORs were aligned using muscle3[8]. In cases of aligning three StVs per chromosome, the alignment was performed in stages in order to avoid wrong alignment. HORs of each of the three StVs were aligned separately with one copy of full-length StV using muscle3. Then, using a custom Python script (merge_alignments.py from https://github.com/fedorrik/horhap_tool), the alignments of three StVs were merged. The pairwise Hamming distances between all the aligned sequences were calculated, and the hierarchical Ward's linkage clustering was performed to divide the HORs into k clades. For each pair of chromosomes, bed files and alignment figures for all k from two to nine were generated, and after inspection, only one k for each chromosome was selected (supplement table with selected k). As a result, most HORs got HORhap assignments, which were added to the HOR name (e.g., S2C9H1L.1-7:C2, where C2 is the 2nd HORhap).

HOR monomer, STV, and HORhap tracks are available on the UCSC Genome Browser track hub[1] (PMID: 24227676) (https://genome-euro.ucsc.edu/s/fedorrik/HG002v1.1.PAT), an example of tracks is in SNote Figure 2. A graphical overview of the HORhap workflow is shown in SNote Figure 3. Ward dendrograms and alignment plots generated by the horhap tool for all analyzed chromosomes are shown in SNote Figure 4. HORhap tracks of all analyzed chromosomes are shown in SNote Figure 7. Note that visible gaps in the HORhap tracks are caused by local high concentrations of relatively rare StVs, which did not get HOR assignments.

HORhap-assigned fasta datasets can be used to train HMM models, and along with the HumAS-HMMER tool, they can be used to rapidly annotate other assemblies without de novo generation of HORhaps. In this work, the HMMer stage was not applied since both homologs were already covered by HORhap analysis.

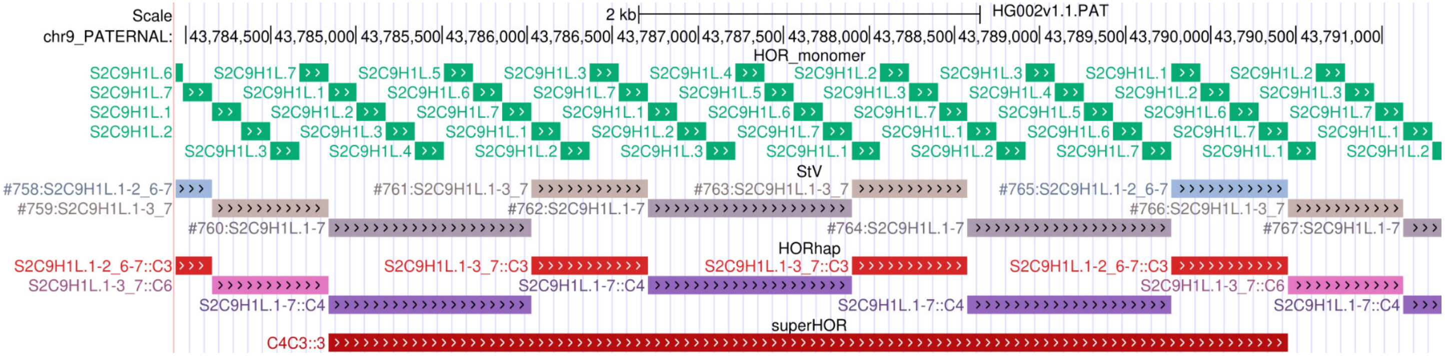


**SNote Figure 2**. Screenshot from the UCSC browser shows tracks (from top to bottom): HOR monomer, StV (monomers merged in complete HORs), HORhap (subtle sequence variants of HORs), superHOR (periodical structures formed by HORhap assignments).

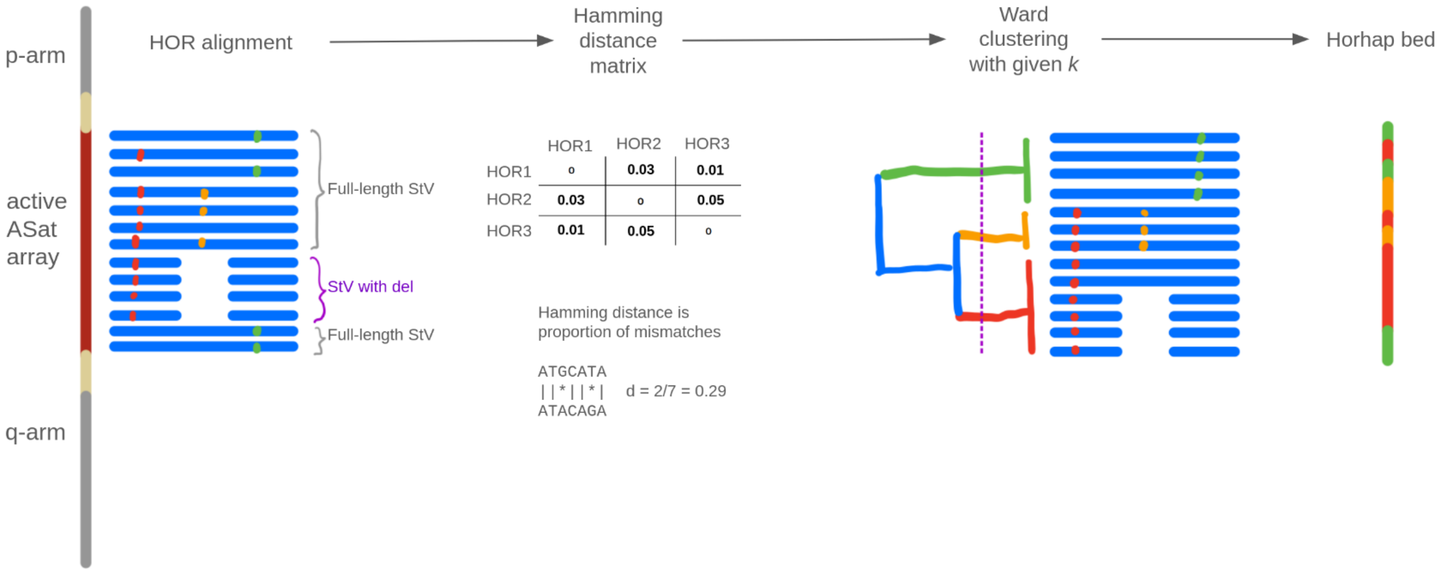


**SNote Figure 3.** Overview of the HORhap workflow, where the first step is generating an alignment of all annotated HORs from the active alpha satellite array on each HG002 phased assembly. In this step, HOR (shown in blue) may be full-length (Full-length StV) or the most frequent structural variation ('StV with del'). Small variants (SNVs and submonomeric INDELs) are highlighted in color (red, orange, and green). Next, we perform pairwise comparisons with each HOR to populate a Hamming distance matrix. We use Ward clustering with a given k to form HOR groups and issue group labels back to the original HOR annotation (HORhap bed).

| chrom | 1 | 2 | 3 | 4 | 5 | 6 | 7 | 8 | 9 | 10 | 11 | 12 | 13 | 14 | 15 | 16 | 17 | 18 | 19 | 20 | 21 | 22 | X | Y |
|---|---|---|---|---|---|---|---|---|---|---|---|---|---|---|---|---|---|---|---|---|---|---|---|---|
| number of HORhaps k | 4 | 4 | 5 | 9 | 4 | 5 | 4 | 5 | 6 | 3 | 4 | 6 | 6 | 8 | 9 | 8 | 5 | 4 | - | 6 | 5 | 7 | 7 | 3 |
| number of StVs | 2 | 3 | 2 | 2 | 2 | 1 | 1 | 3 | 3 | 2 | 2 | 2 | 2 | 1 | 2 | 2 | 2 | 2 | - | 2 | 2 | 1 | 1 | 2 |

**SNote Table 1.** Selected number of HORhaps and number of frequent StVs taken into the analysis for each chromosome.

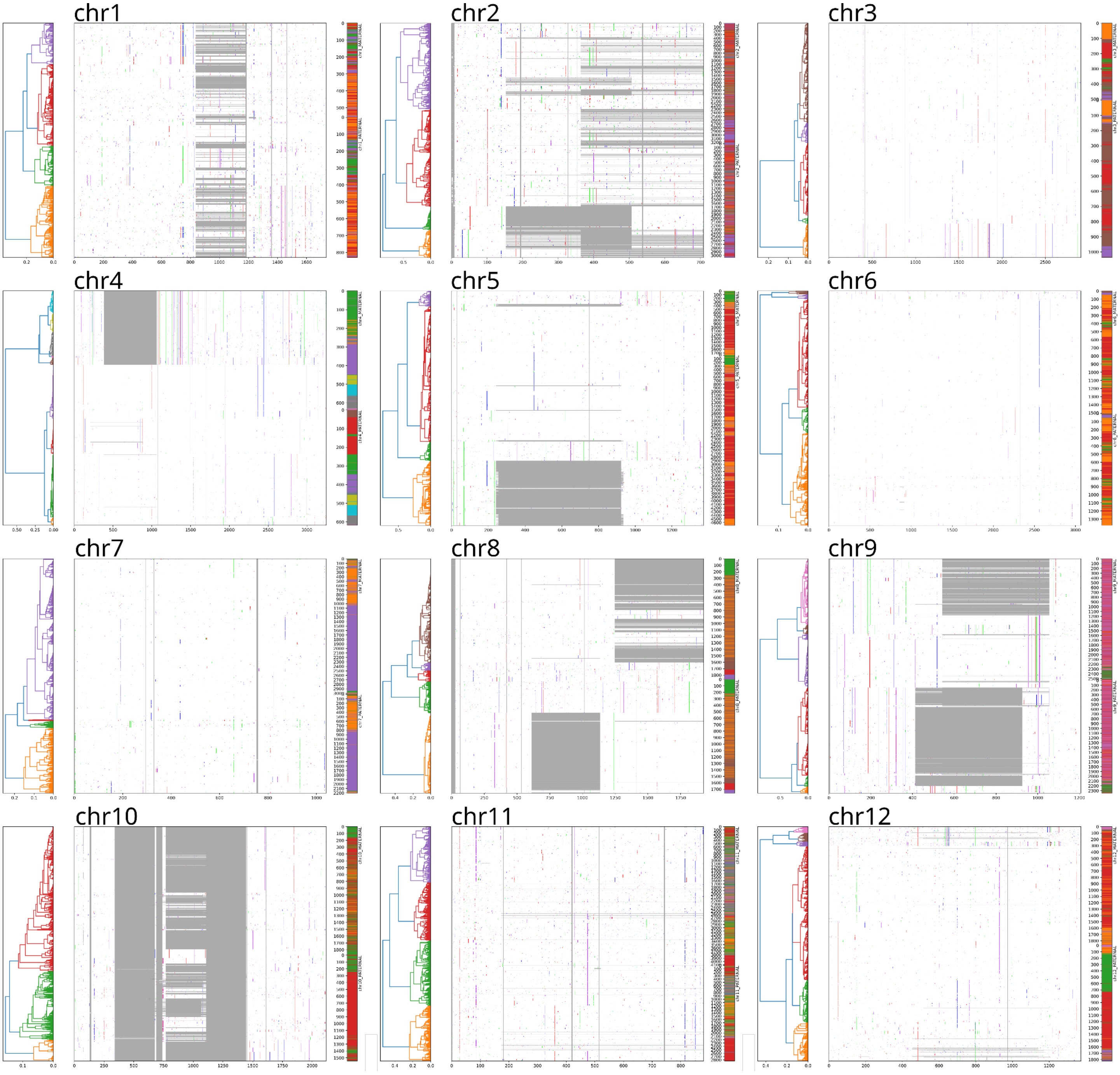
chr1
chr2
chr3
chr4
chr5
chr6
chr7
chr8
chr9
chr10
chr11
chr12

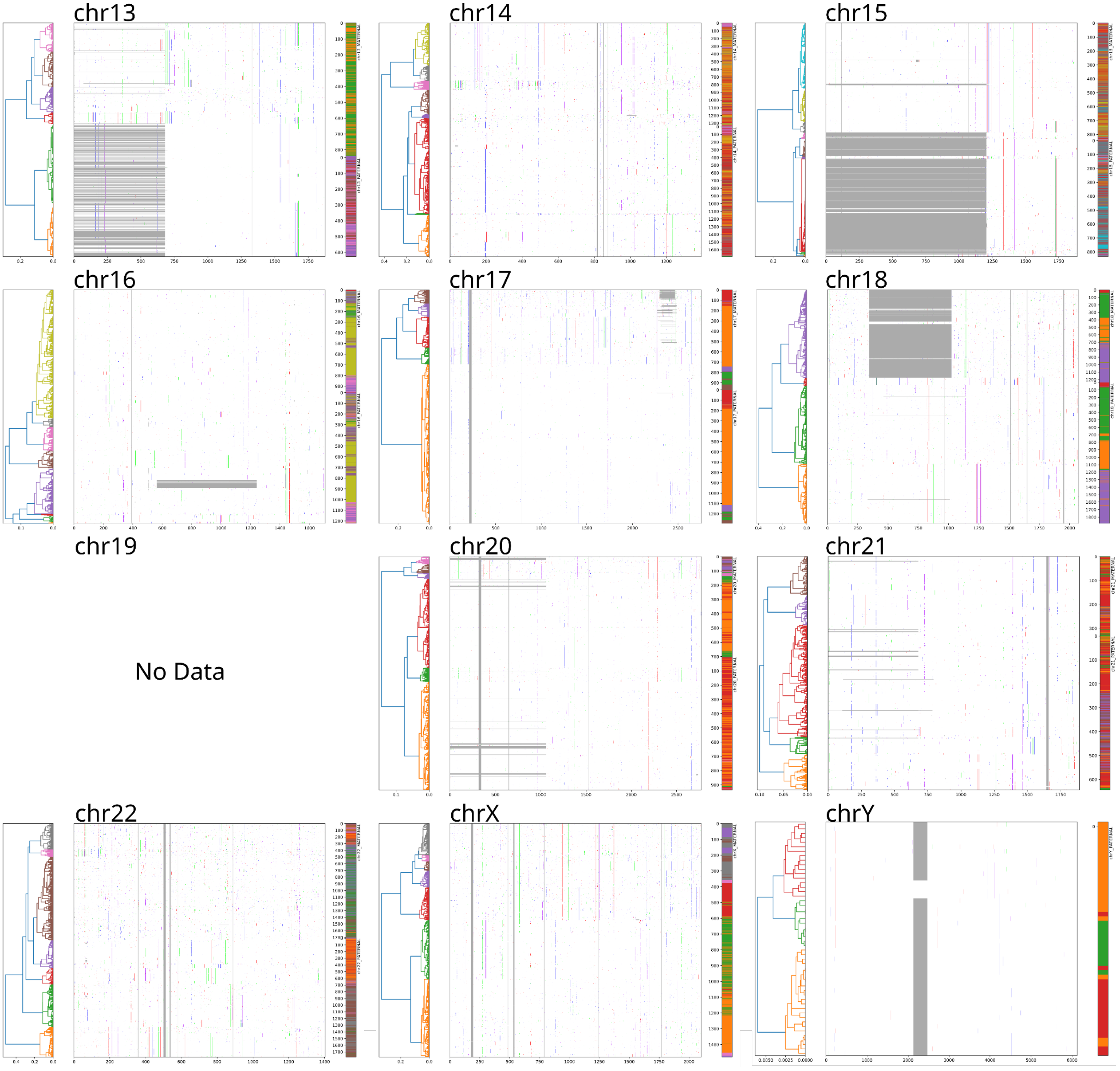


**SNote Figure 4**. Ward HORhap clustering for all chromosomes in HG002. Both maternal and paternal HORs are aligned together and clustered in k groups (center), and the clustering hierarchy is visualized as a tree with k clades highlighted by different colors (left). Paternal and maternal centromeres (one on top of the other) are shown on the right using respective coloration. In the alignment view (center), variants not shared with a consensus are shown along the length of a HOR (x-axis), and deletions are visible in grey. Whole-monomer deletions characterize StVs used for analysis.

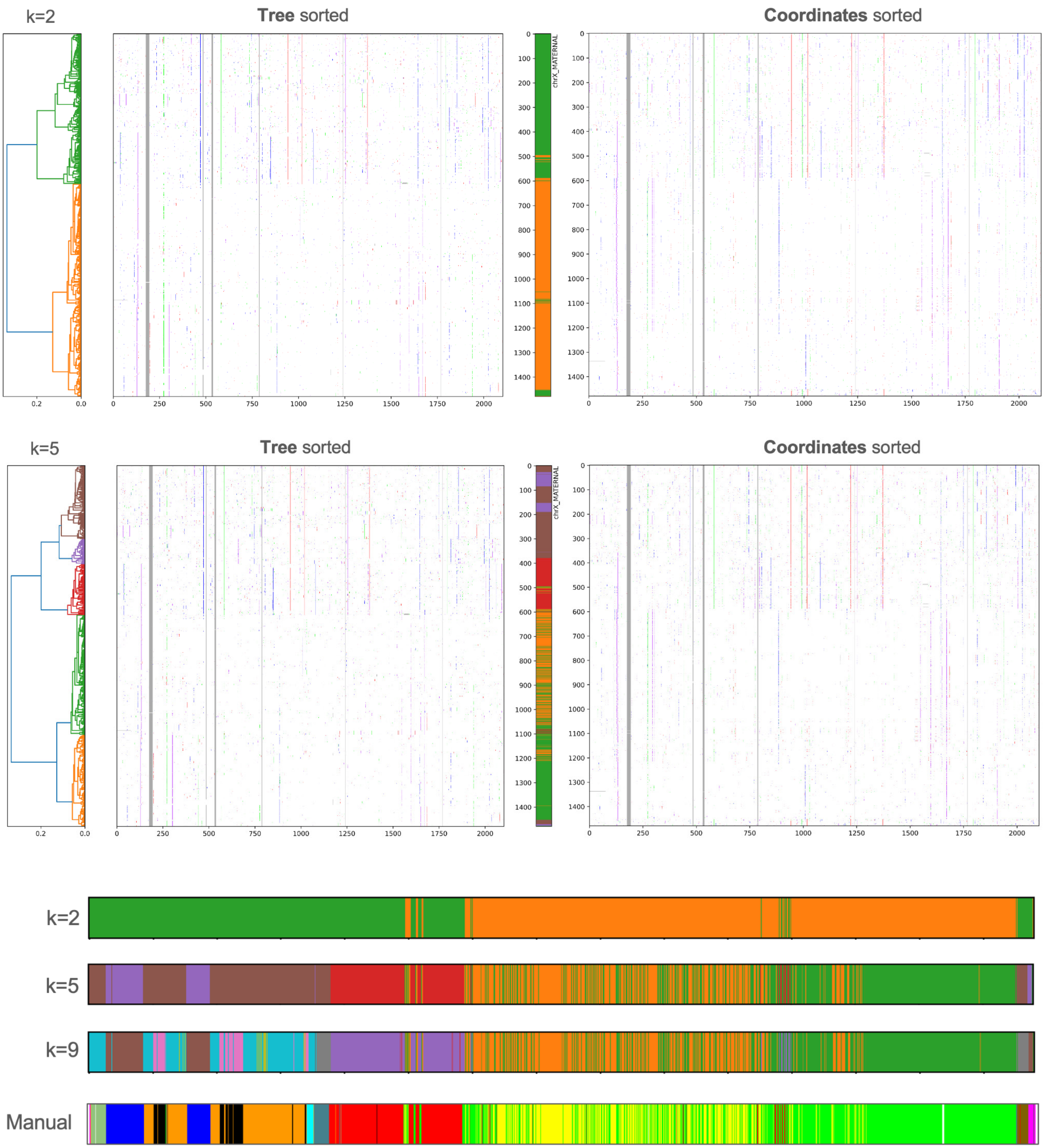


**SNote Figure 5**. Comparison of CHM13 chrX HORhaps with different k (number of HORhaps) produced by the horhap tool with reference manual HORhap annotation from Altemose 2022[7]. K9 annotation looks very similar to the manual one, showing that the 2 methods give consistent results.

**HORhap Age Analysis**
Prior work in CHM13[7] demonstrated that the kinetochore tends to localize over younger HORhap arrays, suggesting a localization preference for more recently expanded centromeric sequence or, conversely, a kinetochore-facilitated expansion (kinetochore selection hypothesis) [7,9]. To assess whether this pattern holds in HG002, we estimated the relative age of each HORhap by calculating the average pairwise sequence divergence among HOR units within a given HORhap. HORhaps with higher divergence are presumed to have expanded earlier, while those with lower divergence reflect more recent expansions (SNote Figure 6). HORhaps were colored accordingly (gray = older, red = younger), and phylogenetic trees of HORhap consensus sequences were constructed to further inform age relationships where divergence values alone were ambiguous **STable 9, 10**. HORhap annotations were then intersected with centromere dip regions (CDRs), used here as a proxy for kinetochore location. Of 46 chromosomes with well-distinguished HORhaps, 28 had CDRs overlapping a younger HORhap, and 9 overlapped an older one (9 chromosomes were excluded due to poorly resolved HORhap structure) **STable 9, 10**. While this distribution is consistent with the trend observed in Altemose et al 2022[7], accounting for the relative sizes of younger versus older HORhap arrays reduces the apparent effect. A binomial test against the null hypothesis of no kinetochore preference failed to reach significance ($p = 0.092$), suggesting that while younger HORhaps are nominally favored, this cannot be established as a statistically robust trend in HG002 alone.

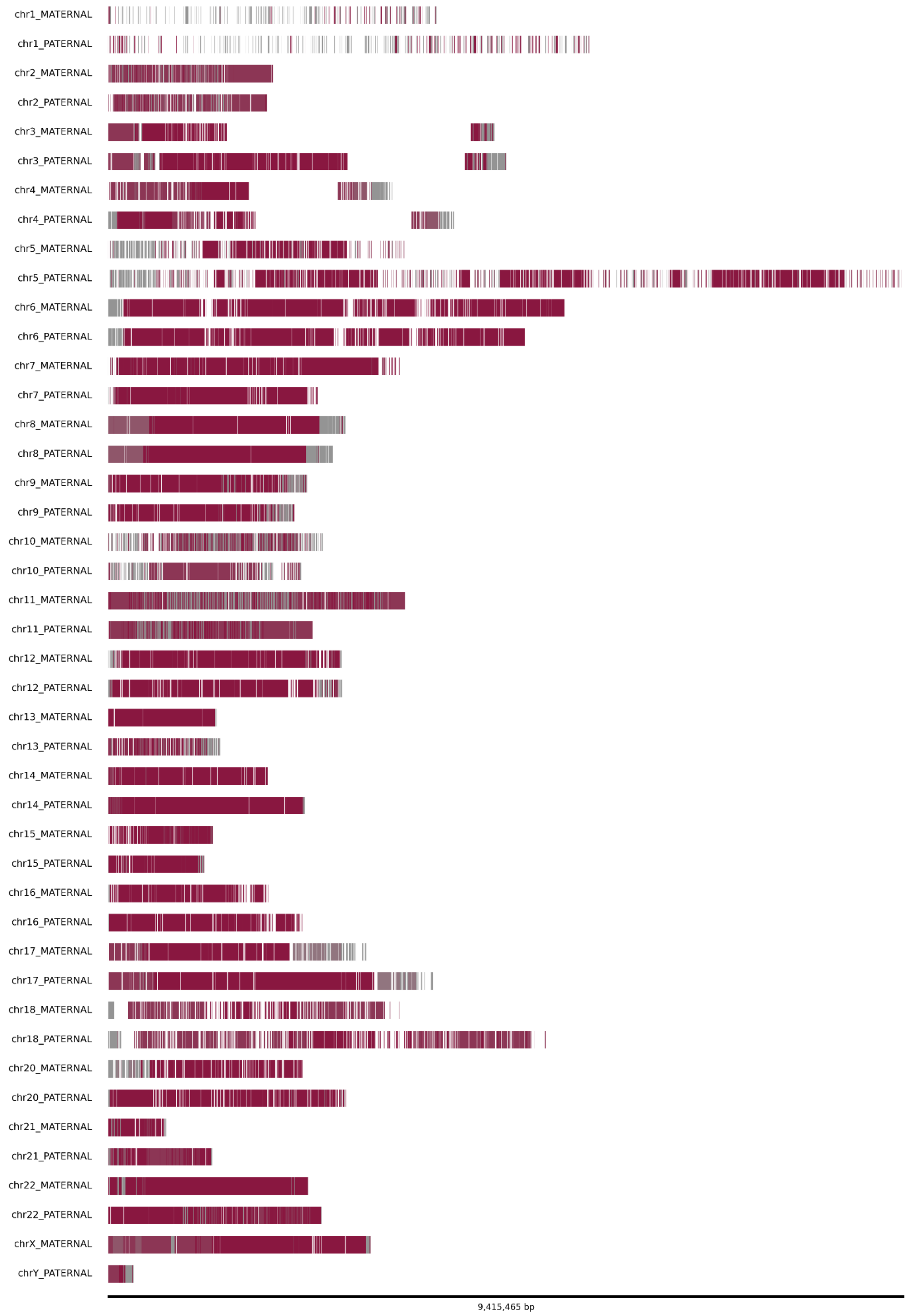


**SNote Figure 6.** HORhap tracks are colored according to divergence-based relative age estimates (gray = older, red = younger).

**SuperHOR Annotation**
To investigate whether HOR units form higher-order repetitive structures beyond the standard HOR periodicity, we produced a superHOR annotation for HG002. Using a custom script (https://github.com/fedorrik/superHOR_HG002), HORhap annotations were scanned for n-mers of HORs (n = 2–9) repeated at least three times consecutively. Each "unit" in the n-mer is itself a HOR, and at least one HOR in an n-mer should have a HORhap assignment different from the others (SNote Figure 7). Identified superHOR arrays are colored by their n value, providing an immediate visual indication of the periodicity in each superHOR array.SuperHOR annotation track is available on the UCSC browser, and an example is shown in SNote Figure 2.

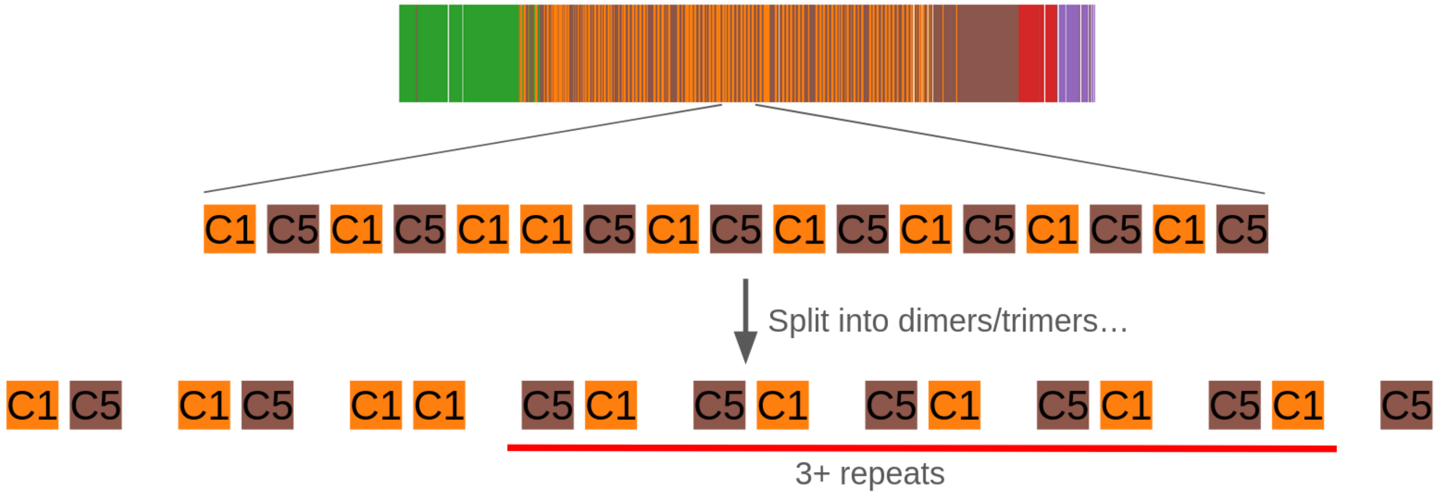


**SNote Figure 7**. Schematics of superHOR annotation. Combinations of two HORhap assignments C5 and C1 are repeated 5 times, forming a superHOR array.

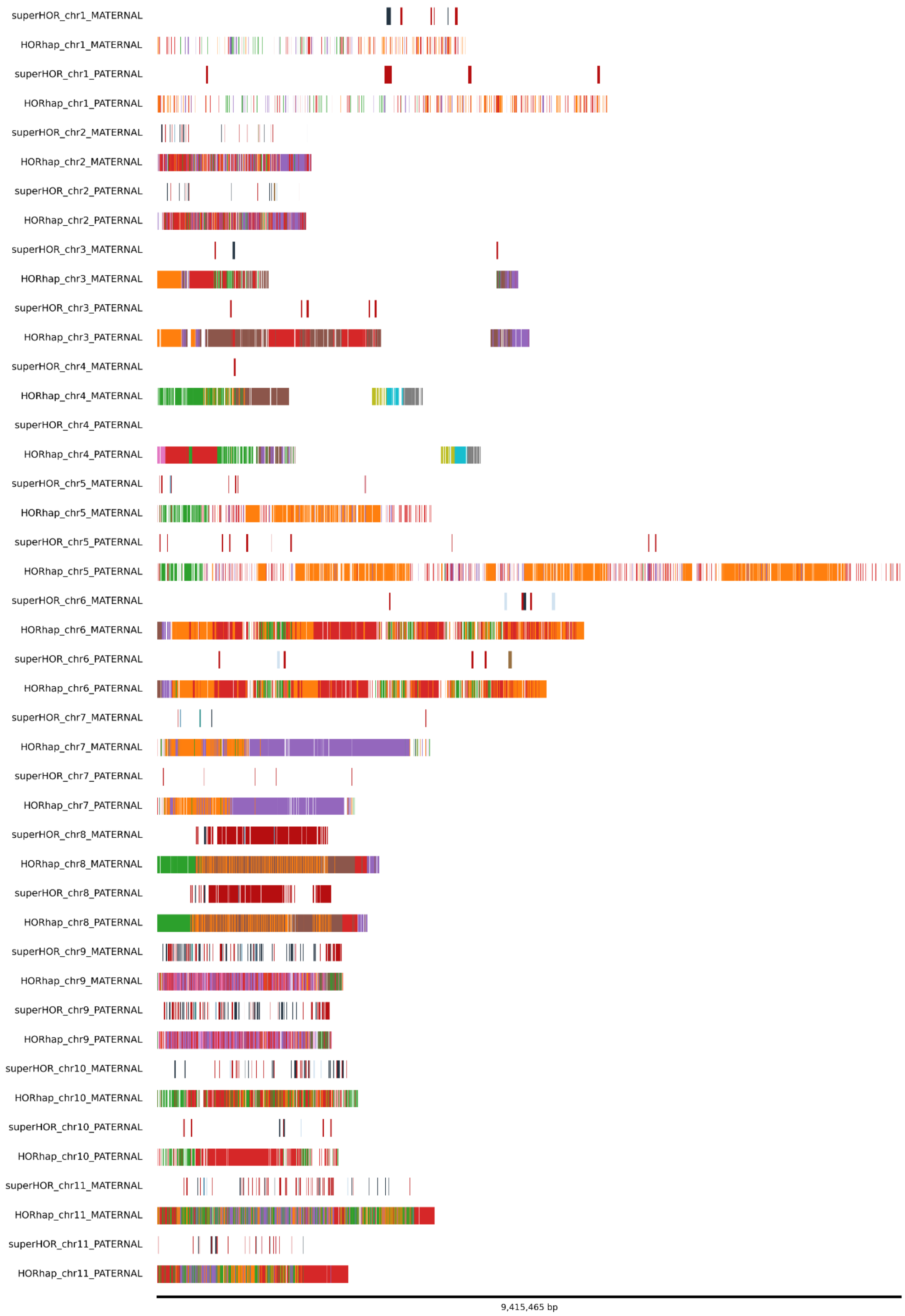
superHOR_chr1_MATERNAL
HORhap_chr1_MATERNAL
superHOR_chr1_PATERNAL
HORhap_chr1_PATERNAL
superHOR_chr2_MATERNAL
HORhap_chr2_MATERNAL
superHOR_chr2_PATERNAL
HORhap_chr2_PATERNAL
superHOR_chr3_MATERNAL
HORhap_chr3_MATERNAL
superHOR_chr3_PATERNAL
HORhap_chr3_PATERNAL
superHOR_chr4_MATERNAL
HORhap_chr4_MATERNAL
superHOR_chr4_PATERNAL
HORhap_chr4_PATERNAL
superHOR_chr5_MATERNAL
HORhap_chr5_MATERNAL
superHOR_chr5_PATERNAL
HORhap_chr5_PATERNAL
superHOR_chr6_MATERNAL
HORhap_chr6_MATERNAL
superHOR_chr6_PATERNAL
HORhap_chr6_PATERNAL
superHOR_chr7_MATERNAL
HORhap_chr7_MATERNAL
superHOR_chr7_PATERNAL
HORhap_chr7_PATERNAL
superHOR_chr8_MATERNAL
HORhap_chr8_MATERNAL
superHOR_chr8_PATERNAL
HORhap_chr8_PATERNAL
superHOR_chr9_MATERNAL
HORhap_chr9_MATERNAL
superHOR_chr9_PATERNAL
HORhap_chr9_PATERNAL
superHOR_chr10_MATERNAL
HORhap_chr10_MATERNAL
superHOR_chr10_PATERNAL
HORhap_chr10_PATERNAL
superHOR_chr11_MATERNAL
HORhap_chr11_MATERNAL
superHOR_chr11_PATERNAL
HORhap_chr11_PATERNAL
9,415,465 bp

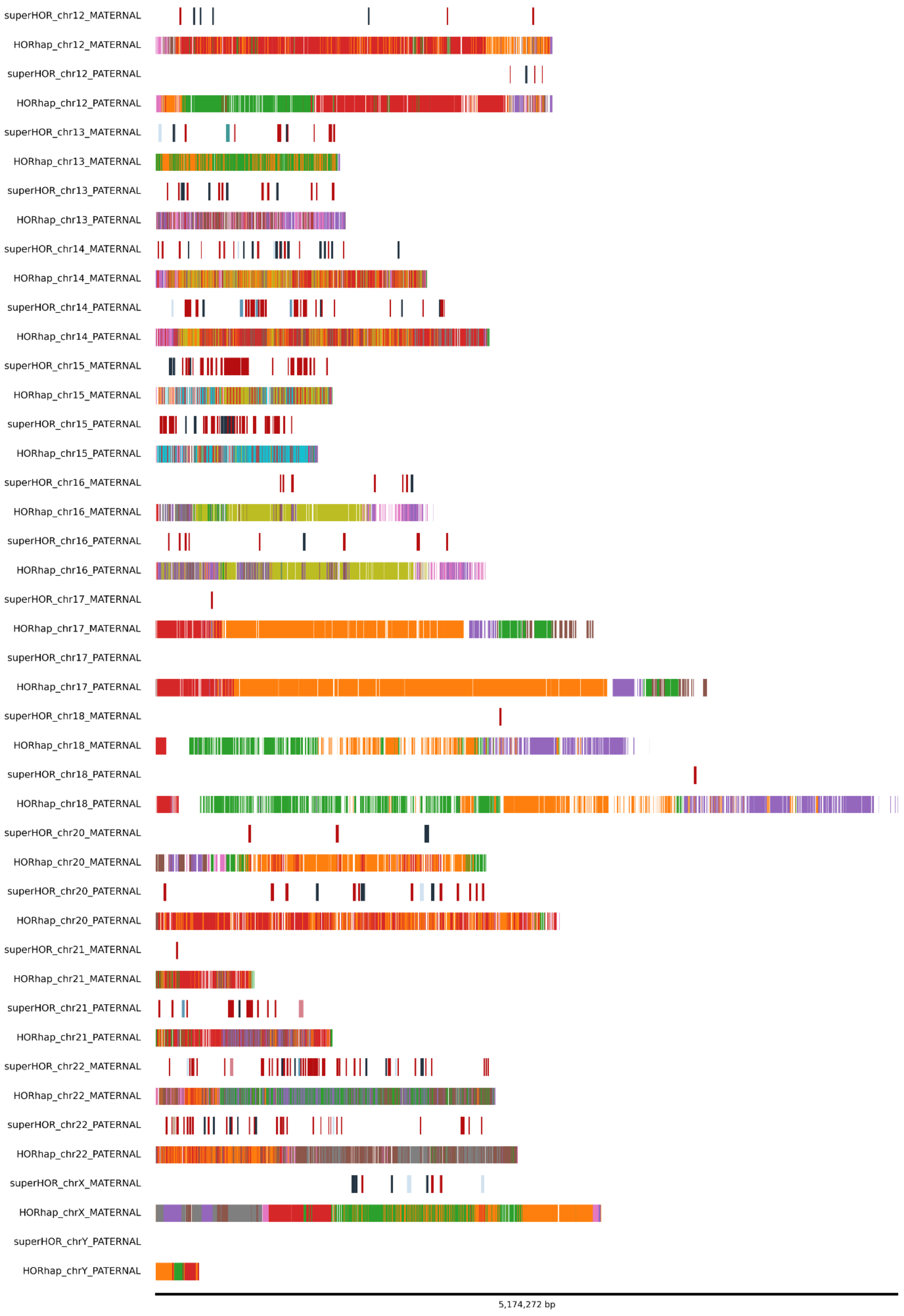


**SNote Figure 8.** HORhap and superHOR tracks of each analyzed chromosome of HG002.

### Supplemental Note 2: Centromere mappability assessment and aligner evaluation

#### Overview

Accurate genome assembly of human centromeres remains a fundamental challenge due to the highly repetitive nature of centromeric satellite DNA, including alpha satellites organized into higher-order repeats (HORs) and human satellite sequences (HSat1–3). Existing mappability tools were developed for short-read sequencing and have not been validated on fully assembled centromeric sequences. Moreover, they do not capture the mappability characteristics of long reads, which are now central to efforts to resolve previously intractable genomic regions. Here, we describe a method for generating long-read centromere mappability tracks and evaluate the performance of two state-of-the-art aligners, Minimap2[10] and Winnowmap[11], for confidently mapping reads to these regions.

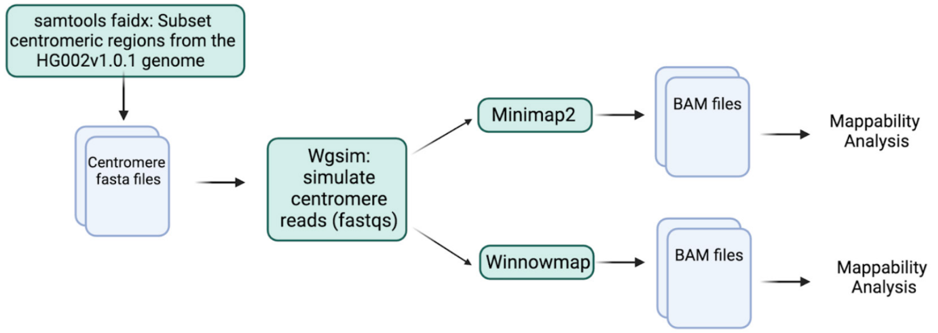


**SNote Fig 1:** Centromere Mapping Analysis Workflow

#### Generating Centromere Mappability Tracks

To establish a ground truth for read alignment performance, we simulated error-free reads from the centromeric regions of the T2T-HG002 (HG002v1.0.1) reference genome[12], a well-characterized human genome benchmark derived from the Ashkenazi cell line HG002. Centromeric regions were defined by the coordinates of the active higher-order repeat (HOR) array for each chromosome, expanded by 5 Mb on both sides to capture flanking satellite sequences and transition regions. Using the read simulator wgsim[13], we generated 10,000 error-free reads per chromosome at eleven read sizes ranging from 1 kb to 300 kb, as well as 30x coverage simulations for each read size. Reads were simulated with 0% error rate to eliminate sequencing noise as a confounding variable, enabling a clean assessment of how genomic sequence complexity alone affects alignment accuracy. Simulated reads were aligned to the full diploid HG002 reference genome using both Minimap2 v2.26 and Winnowmap v2.03 under ONT parameters (map-ont).

Each aligned read was classified into one of six categories: (1) uniquely mapped reads with a single, perfect alignment to the correct haplotype; (2) multi-mapping reads with one top alignment to the correct haplotype; (3) multi-mapping reads with two top alignments both to the correct haplotype; (4) multi-mapping reads with two top alignments to different haplotypes, or three or more top alignments; (5) reads mapping to an incorrect chromosome; and (6) unmapped reads. A "top alignment" was defined by a perfect CIGAR string, zero edit distance (NM:i:0), and an alignment score equal to twice the read length, strict criteria appropriate for error-free simulated reads.

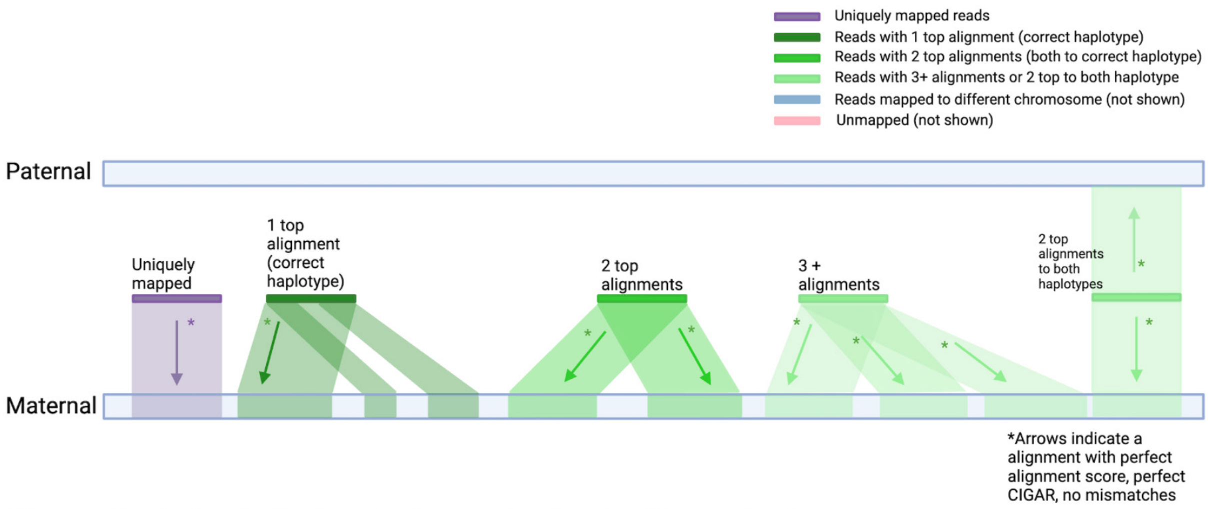


**SNote Fig 2**: Diagram of read characterization for uniquely mapped reads and multi-mapped reads

Mappability tracks were generated in BED format by recording genomic intervals covered by uniquely mapped reads at each read size. These tracks reveal that mappability increases substantially with read length. Active HOR arrays, which are the functionally critical regions of the centromere where CENP-A binds to facilitate kinetochore assembly, become fully mappable at read sizes of 40–60 kb for most chromosomes, indicating that these arrays are largely haplotype-specific. Diverged and monomeric HOR regions remain largely unmappable until read sizes of 200 kb or greater, consistent with their higher degree of sequence similarity across haplotypes and homologous chromosomes. Human satellite regions, including HSat1A and portions of HSat2, also become uniquely mappable at 60 kb, suggesting these sequences are sufficiently haplotype-specific to support confident alignment at ultra-long read lengths. Notably, non-centromeric flanking regions showed few mappable intervals despite their presumed lower repetitiveness, likely because reads originating from these regions align preferentially to other haplotypic copies as non-primary alignments. These mappability tracks provide a practical resource for filtering variant calls and assembly evaluations to regions of high alignment confidence.

#### Aligner Comparison: Minimap2 vs. Winnowmap

A key variable in centromere assembly accuracy is the choice of alignment tool. Minimap2 is a widely adopted, general-purpose long-read aligner that performs efficiently across diverse genomic contexts. Winnowmap, built on the Minimap2 codebase, introduces a weighted minimizer sampling algorithm designed to downweight frequently occurring k-mers during the seeding stage, a modification specifically intended to improve alignment accuracy in regions dominated by tandem repeats and other repetitive sequences. Winnowmap additionally uses minimal confidently alignable substrings to mitigate allelic bias, making it more sensitive to paralog-specific variants in complex genomic regions. Our simulations show that Winnowmap outperforms Minimap2 for aligning shorter centromeric reads. At 10 kb, Winnowmap uniquely maps more than 90% of simulated centromeric reads for most chromosomes, while Minimap2 achieves only approximately 50% unique mapping at the same size. Minimap2's unique alignment rate increases gradually following an approximately exponential curve, whereas Winnowmap shows a marked improvement in unique alignment beginning at 10 kb. This difference likely reflects Winnowmap's optimized handling of the high-frequency k-mers characteristic of centromeric satellite sequences. At larger read sizes (60 kb and above), both aligners converge on high unique mapping rates for the active HOR, with differences becoming less pronounced. This difference can be alleviated by the subsequent filtering of bam files for the alignment quality.

In terms of computational performance, Minimap2 completes alignments substantially faster than Winnowmap, consistent with Winnowmap's more intensive evaluation of candidate alignments. Chromosomes with the lowest and shortest mappable regions also required the longest alignment times, suggesting that the computational cost of Winnowmap scales with the complexity of the repetitive landscape.

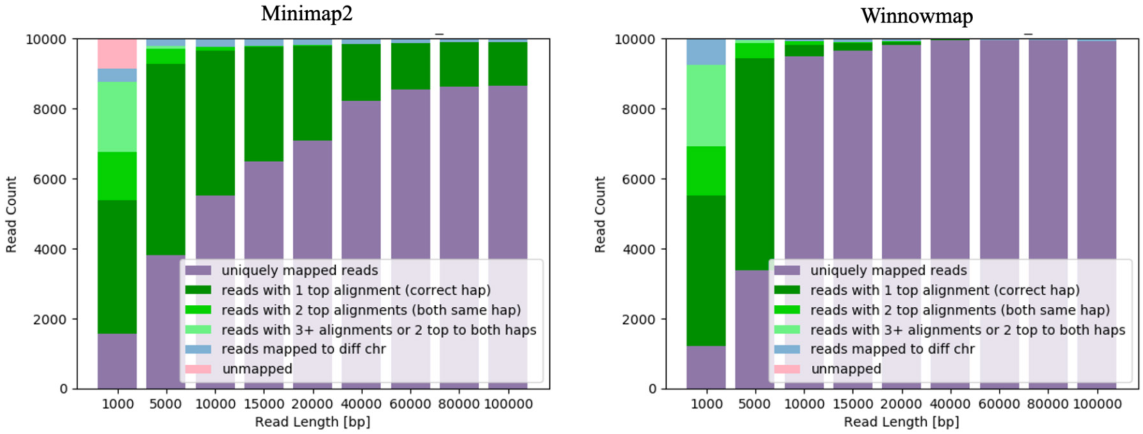


**SNote Fig 3**: Comparison of read breakdown for Minimap2 vs Winnowmap alignment results for chr1_MATERNAL centromere

Comparison of variant detection using experimental ONT reads with NucFreq[14] provides an additional lens for evaluating aligner performance. For the majority of chromosomes, Minimap2 identifies slightly fewer total variants than Winnowmap, with a high proportion of variants shared between the two aligners, indicating that most detected variants reflect consistent sequence differences rather than aligner-specific artifacts. On chromosomes 6 and 7, however, Minimap2 produces a higher variant count associated with coverage irregularities, including a prominent coverage peak at the start of the active HOR, suggestive of a read collapse in which reads from low-coverage regions mismap to a single locus. This observation prompted the development of the lr:hq preset that resolves this issue in minimap2. Winnowmap does not exhibit this behavior, indicating superior alignment fidelity in these cases.

Taken together, these results support the use of Winnowmap for centromere-focused assembly and quality control workflows, particularly when aligning reads at the 10–20 kb range typical of high-accuracy HiFi sequencing. For applications requiring faster turnaround and where reads exceed 60 kb, Minimap2 represents a reasonable alternative with comparable unique mapping rates in most centromeric regions.

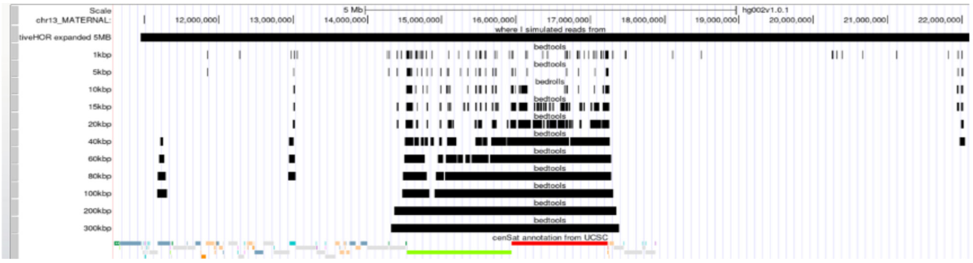


**SNote Fig 4**: The black tracks are mappability tracks of read sizes 1kb to 300kb for the regions where simulated reads uniquely map to for centromeric regions chr13 maternal, the bottom tracks are CenSat annotations (light green: HSat and red: alpha satellite)

### Supplemental Note 3: Quality Control and Characterization of hiPSC Lines

#### Overview

All cell lines used in this study were obtained from the Coriell Institute for Medical Research. The lymphoblastoid cell line (LCL) GM24385, induced pluripotent stem cell (hiPSC) line GM26105 (reprogrammed from GM24385), and hiPSC line GM27730 (reprogrammed from peripheral blood mononuclear cells, PBMCs) are derived from the same Personal Genome Project (PGP) participant, huAA53E0, also known as HG002 in the NIST Genome in a Bottle (GIAB) dataset. GM26105 was generated using an episomal reprogramming approach, while GM27730 was generated using the Sendai viral reprogramming. Initial genomic characterization of GM26105 and GM27730, including short-read whole-genome sequencing, karyotyping, pluripotency marker validation, and confirmation of high single-nucleotide variant (SNV) and small indel (<50 bp) concordance with the donor reference, was previously described in Scheinfeldt *et al.* (2025)[15]. Here we report additional quality control analyses performed specifically to support the centromere-focused studies described in the main text, including long-read nanopore sequencing, structural variant and genome integrity assessment, CDR calling, and long-read RNA expression profiling of CENP-A.

#### hiPSC Culture Conditions

Human iPSC lines were cultured as previously described[15], without antibiotics or antifungals in mTeSR1 medium (STEMCELL Technologies) and maintained at 37°C with 5% $CO_2$ and ambient oxygen levels on Matrigel-coated tissue culture vessels. Cells were passaged when reaching approximately 75–85% confluency at a split ratio of 1:3–1:6. Subcultivation was performed using standard dissociation reagents (Versene or ReLeSR), and cells were reseeded in fresh Matrigel-coated vessels with mTeSR1 medium. Medium was replaced daily to maintain optimal culture conditions. Cell density and morphology were monitored routinely to ensure healthy proliferation and the absence of contamination. For cryopreservation, cells were frozen in a medium composed of 90% KnockOut Serum Replacement and 10% DMSO.

#### Generation of Nanopore Long-Read Data

To enable assessment of structural variation and centromere-level genome integrity, we generated Oxford Nanopore Technology (ONT) ultra-long (UL) whole-genome sequencing (WGS) data from GM26105 and GM27730. DiMeLo-seq[6] was performed on the PromethION platform (PromethION, R10.4.1) using an ultra-long read protocol optimized for high-molecular-weight DNA isolation and library preparation, for both GM26105 and GM27730. In this experiment, we utilized a modified partial ultra-long method, where we added a step of shearing the DNA by pipetting with a standard P1000 pipette tip. This step was performed until no viscosity was observed in the samples. Furthermore, during the fragmentation mix step after the addition of fragmentase, we utilized the standard P1000 pipette tips instead of the widebore P1000 tips. Both experiments achieved read length N50s of 108,349 bp and 96,890 bp, respectively. Raw signal data were basecalled using Dorado v1.0.2 with modified base detection enabled for 5-methylcytosine (5mC) and N6-methyladenine (6mA) using dna_r10.4.1_e8.2_400bps_sup@v5.2.0. The coverage for GM26105 was 9.4812, and the average coverage for GM27730 was 16.9247.

#### Evaluation of Genome Integrity and Structural Variation

Prior QC of these lines[15] confirmed high concordance of SNVs and small indels relative to the GIAB HG002 reference callset; however, structural variants (SVs) and large-scale rearrangements, particularly within centromeric satellite arrays, were not systematically evaluated. These regions are prone to somatic copy number variation and rearrangement during reprogramming and extended culture, and instability in centromeric sequence could confound downstream analyses of centromere identity and chromatin organization. Furthermore, the use of the complete T2T-HG002 assembly as the alignment target introduces unique mapping considerations in centromeric regions not encountered with GRCh38.

To assess large-scale genome integrity, we applied Flagger[16], a reference-based read-depth analysis tool designed to flag regions of unexpected coverage in long-read assemblies and alignments. Flagger partitions the genome into coverage-state categories, including haploid, diploid, collapsed (over-represented), and erroneous (under-represented) windows, by modeling expected read depth distributions and identifying statistically anomalous regions. Critically, Flagger was developed and validated in the context of complete T2T assemblies, where coverage anomalies in centromeric and segmentally duplicated regions are particularly informative. We focused Flagger analysis on centromeric regions of all 23 chromosome pairs to identify any evidence of copy number change, deletion, or collapse in the two hiPSC lines relative to the donor LCL.

Using Flagger v1.1.0, we found that 99.3% of the evaluated censat sequence was classified as haploid, with minor fractions flagged as erroneous (0.6%), collapsed (0.1%), or duplicated (0.04%), indicating that centromeric architecture is broadly preserved across both hiPSC lines. Importantly, no assembly errors were reported overlapping with CDR intervals. However, it is possible that sites of decreased methylation (due to read misalignment) may have contributed to the limited false peak calls in the hiPSC that are outside of the primary CDR location (SNote 4).

***STable 1***. *Flagger coverage tracks for active arrays*

| Chromosome | Hap (%) | Err (%) | Col (%) | Dup (%) |
|---|---|---|---|---|
| chr1_MATERNAL | 99.9 | 0.0 | 0.0 | 0.1 |
| chr1_PATERNAL | 100.0 | 0.0 | 0.0 | 0.0 |
| chr2_MATERNAL | 99.9 | 0.0 | 0.1 | 0.0 |
| chr2_PATERNAL | 100.0 | 0.0 | 0.0 | 0.0 |
| chr3_MATERNAL | 99.9 | 0.0 | 0.1 | 0.0 |
| chr3_PATERNAL | 99.8 | 0.0 | 0.1 | 0.0 |
| chr4_MATERNAL | 99.6 | 0.0 | 0.4 | 0.0 |
| chr4_PATERNAL | 99.3 | 0.0 | 0.7 | 0.0 |
| chr5_MATERNAL | 100.0 | 0.0 | 0.0 | 0.0 |
| chr5_PATERNAL | 100.0 | 0.0 | 0.0 | 0.0 |
| chr6_MATERNAL | 100.0 | 0.0 | 0.0 | 0.0 |
| chr6_PATERNAL | 100.0 | 0.0 | 0.0 | 0.0 |
| chr7_MATERNAL | 100.0 | 0.0 | 0.0 | 0.0 |
| chr7_PATERNAL | 100.0 | 0.0 | 0.0 | 0.0 |
| chr8_MATERNAL | 100.0 | 0.0 | 0.0 | 0.0 |
| chr8_PATERNAL | 100.0 | 0.0 | 0.0 | 0.0 |
| chr9_MATERNAL | 100.0 | 0.0 | 0.0 | 0.0 |

| | | | | |
|---|---|---|---|---|
| *chr9_PATERNAL* | *100.0* | *0.0* | *0.0* | *0.0* |
| *chr10_MATERNAL* | *99.9* | *0.0* | *0.0* | *0.0* |
| *chr10_PATERNAL* | *98.7* | *1.3* | *0.0* | *0.1* |
| *chr11_MATERNAL* | *100.0* | *0.0* | *0.0* | *0.0* |
| *chr11_PATERNAL* | *99.9* | *0.0* | *0.0* | *0.1* |
| *chr12_MATERNAL* | *100.0* | *0.0* | *0.0* | *0.0* |
| *chr12_PATERNAL* | *100.0* | *0.0* | *0.0* | *0.0* |
| *chr13_MATERNAL* | *94.5* | *5.2* | *0.3* | *0.0* |
| *chr13_PATERNAL* | *99.7* | *0.0* | *0.1* | *0.2* |
| *chr14_MATERNAL* | *94.7* | *4.7* | *0.6* | *0.0* |
| *chr14_PATERNAL* | *98.5* | *1.4* | *0.1* | *0.0* |
| *chr15_MATERNAL* | *96.3* | *3.4* | *0.2* | *0.1* |
| *chr15_PATERNAL* | *96.9* | *2.9* | *0.3* | *0.0* |
| *chr16_MATERNAL* | *100.0* | *0.0* | *0.0* | *0.0* |
| *chr16_PATERNAL* | *100.0* | *0.0* | *0.0* | *0.0* |
| *chr17_MATERNAL* | *100.0* | *0.0* | *0.0* | *0.0* |
| *chr17_PATERNAL* | *99.8* | *0.0* | *0.0* | *0.2* |
| *chr18_MATERNAL* | *100.0* | *0.0* | *0.0* | *0.0* |
| *chr18_PATERNAL* | *100.0* | *0.0* | *0.0* | *0.0* |
| *chr19_MATERNAL* | *100.0* | *0.0* | *0.0* | *0.0* |
| *chr19_PATERNAL* | *100.0* | *0.0* | *0.0* | *0.0* |
| *chr20_MATERNAL* | *99.8* | *0.0* | *0.0* | *0.2* |
| *chr20_PATERNAL* | *99.1* | *0.0* | *0.2* | *0.7* |
| *chr21_MATERNAL* | *91.8* | *7.8* | *0.4* | *0.0* |
| *chr21_PATERNAL* | *94.0* | *5.2* | *0.8* | *0.1* |
| *chr22_MATERNAL* | *89.6* | *10.0* | *0.4* | *0.0* |
| *chr22_PATERNAL* | *96.4* | *2.2* | *0.4* | *1.1* |
| *chrX_MATERNAL* | *100.0* | *0.0* | *0.0* | *0.0* |
| *chrY_PATERNAL* | *98.9* | *0.0* | *1.1* | *0.0* |

### Supplemental Note 4: HG002 hiPSCs CDR predictions

#### Overview

Centromere Dip Regions (CDRs) mark sites that have reduced CpG methylation relative to the surrounding active αSat HOR array. The HG002 hiPSC active arrays have higher array methylation (average 87% vs ~60% for matched LCLs), and present a challenge to predict sub-CDR boundaries, as shown below for the chromosome 10 PATERNAL array.

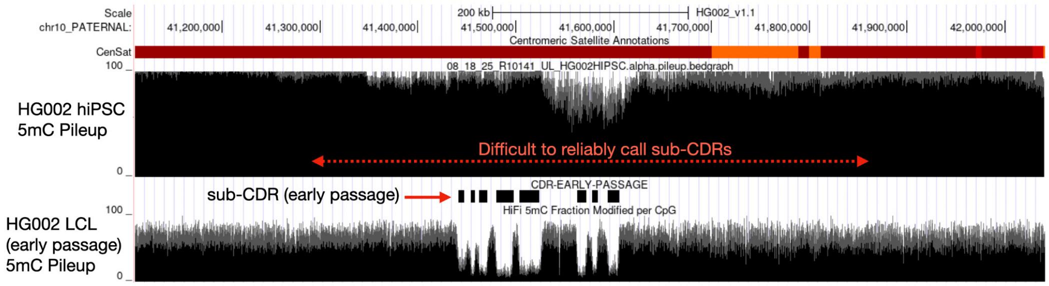


**SNote 4 Figure 1.** UCSC Genome Browser visualization of the CDR region in chr10_PATERNAL. Visualization of the difference in CDR detection between early passage (bottom) to the top hiPSC, which is shallow, and the edges of the subCDR are difficult to confidently call.

Therefore, in an effort to define sub-CDRs in the hiPSC 5mC dataset, we developed a sliding-window CDR caller that detects all contiguous regions falling below a per-array threshold, treats each as a distinct candidate domain, and assigns each domain a composite reliability score to distinguish well-supported calls from marginal detections at the detection boundary. CDR calling was performed on phased, per-CpG methylation bedgraph files derived from Oxford Nanopore long-read sequencing. Each file represents a single haplotype of a single alpha-satellite array, with columns: chromosome/haplotype name, CpG start position, CpG end position, and methylation percentage (0–100). Methylation values were generated using DiMeLo-seq or equivalent long-read methylation-aware base-calling pipelines. Each bedgraph covers the full span of the assembled alpha-satellite array for the corresponding haplotype.

#### CDR Detection Algorithm

Array-Wide Baseline and Detection Threshold

For each haplotype array, the mean CpG methylation was computed across all CpG sites in the input bedgraph. This array-wide mean serves as the baseline methylation level for the array. The CDR detection threshold was set at 10 percentage points below the array-wide (ie, CDR threshold = array mean methylation − 10%). This array-relative threshold accounts for the substantial variation in baseline methylation across centromeric arrays (ranging from approximately 77% on chrY to 96% on chr10) and avoids applying a fixed absolute threshold that would be inappropriately stringent for lower-methylation arrays or permissive for higher-methylation arrays.

Sliding Window Smoothing

A sliding window was applied across each array to smooth the per-CpG methylation signal and reduce the influence of individual CpG-level noise. Windows of 10 kb were stepped at 1 kb intervals across the array coordinates. For each window, the mean methylation of all CpG sites falling within the window boundaries was computed. Windows containing fewer than 3 CpGs were excluded from analysis to avoid unreliable mean estimates in sparsely covered regions.

Domain Calling

CDR domains were identified as contiguous runs of windows whose mean methylation fell below the CDR threshold. Each uninterrupted run of below-threshold windows was treated as a single candidate domain, with its genomic boundaries defined by the positions of the first and last windows in the run. This multi-domain approach explicitly allows for complex CDR architectures containing multiple distinct hypomethylated sub-domains separated by partial methylation recoveries, which are observed on a number of chromosomes. Candidate domains containing no individual CpG observations in the raw bedgraph data (degenerate calls arising from window boundary effects) were discarded.

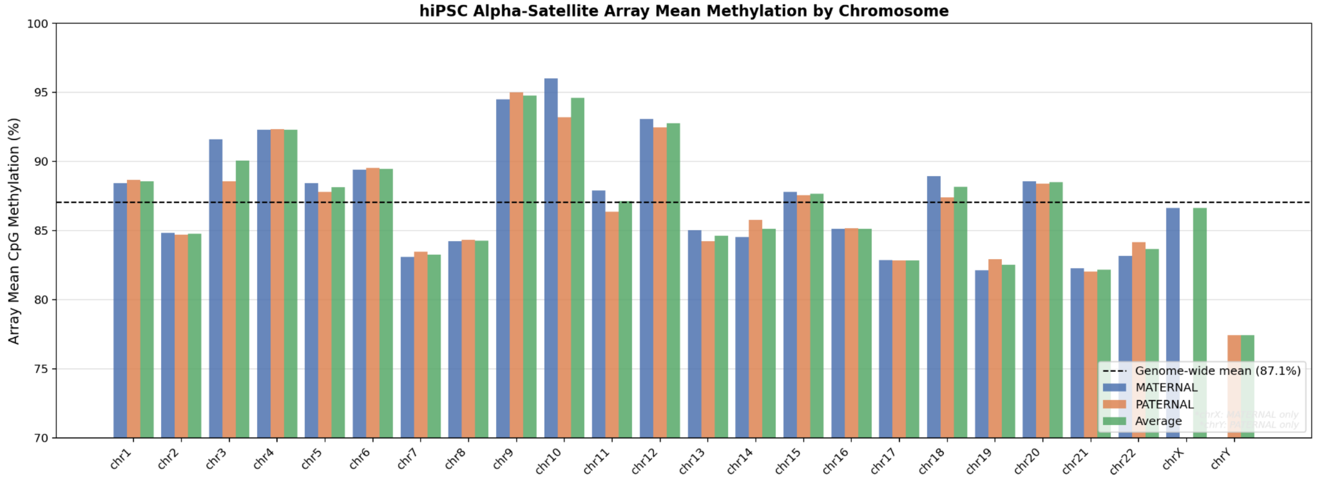


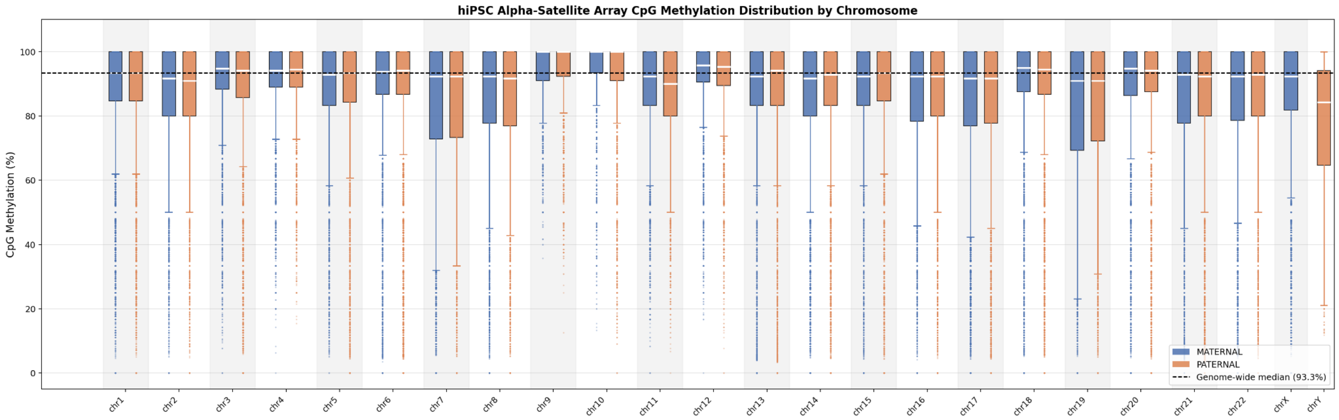


#### Domain Reliability Scoring

The sliding window approach can produce candidate domains that are small, shallow, and sit only marginally below the detection threshold. Such calls may reflect local noise in the methylation signal rather than a genuine hypomethylated sub-domain. To distinguish well-supported CDR domains from marginal calls, we assigned each candidate domain a composite reliability score (0–100) based on three orthogonal properties.

<u>Score Components</u>

The reliability score is computed as a weighted sum of three normalized component scores:

| Component | Weight | Description |
|---|---|---|
| Depth score | 50% | How far the domain mean methylation falls below the detection threshold, normalized to the offset of the threshold. Capped at 1.5× the offset to avoid extreme values dominating the score. Domains at or above the threshold score 0; domains 15+ percentage points below score 1.0. |

| | | |
|---|---|---|
| CpG count score | 30% | Number of individual CpG observations within the domain, capturing the statistical reliability of the mean methylation estimate. Saturates at 100 CpGs (score = 1.0). Domains supported by very few CpGs (<20) are penalised. |
| Span score | 20% | Domain size in kilobases, as a proxy for biological relevance. CDRs are typically tens to hundreds of kilobases in span; very small domains (<5 kb) are penalised. Saturates at 20 kb (score = 1.0). |

The composite score is computed as:
**Score = 0.50 × depth_score + 0.30 × cpg_score + 0.20 × span_score**

where:
**depth_score** = min((threshold − domain_mean) / threshold_offset, 1.5) / 1.5
**cpg_score** = min(nCpG / 100, 1.0)
**span_score** = min(span_kb / 20, 1.0)

The final score is multiplied by 100 to yield a 0–100 scale and clamped to zero to handle edge cases where domain mean methylation exceeds the threshold (e.g., degenerate calls).

Confidence Classification
Domains with a reliability score ≥ 50 are classified as HIGH_CONFIDENCE; domains with a score < 50 are classified as LOW_CONFIDENCE. The threshold of 50 was selected by inspection of the score distributions across all chromosomes: it reliably excluded domains with very small size (<5 kb), very few CpGs (<30), and methylation means within 2–3 percentage points of the threshold, while retaining domains with substantive hypomethylation and adequate CpG support.

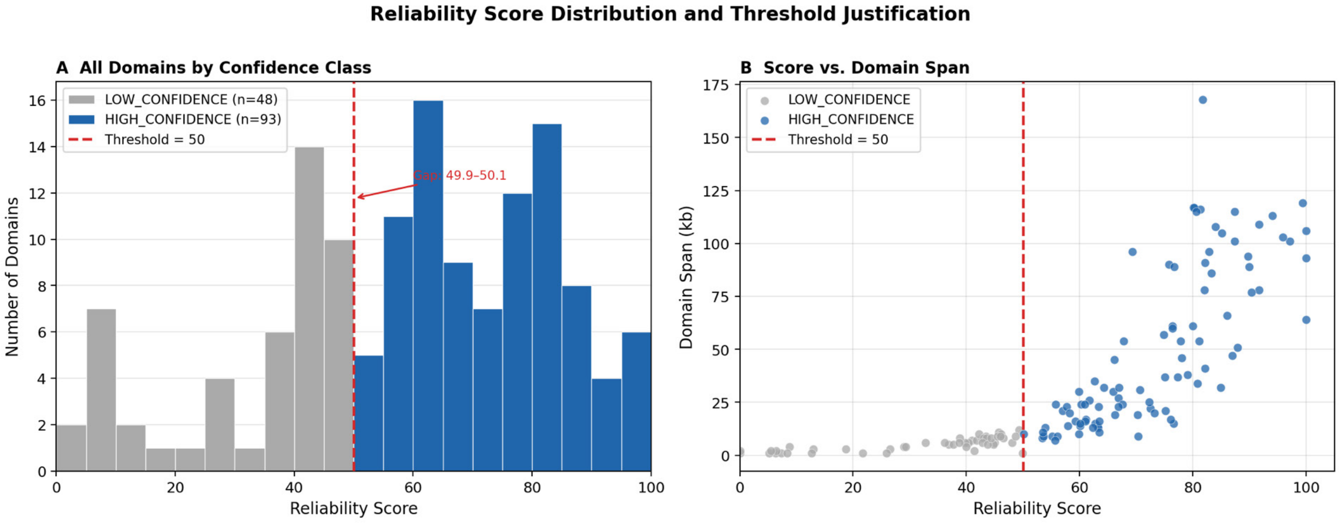


The data make a strong empirical case for a cutoff at 50. Across all 141 domains called genome-wide, there is a natural gap in the score distribution: the maximum LOW_CONFIDENCE score is 49.9 and the minimum

HIGH_CONFIDENCE score is 50.1 — the two classes do not overlap at all. This is not an artifact of the cutoff being imposed; the scoring components (depth, CpG count, span) jointly produce a bimodal distribution where marginal domains cluster in the 0–50 range and well-supported domains cluster from ~50 upward. Panel B reinforces this: LOW_CONFIDENCE domains (grey) are uniformly small in span (<20 kb), while HIGH_CONFIDENCE domains (blue) span a broad range reflecting genuine CDR structural diversity. The threshold of 50 sits cleanly in the gap between these two populations.

However, we noted that specific chromosomes, like chromosome 21 MATERNAL are much smaller than the other CDR estimates, and have several domains in the score category of 40-49.

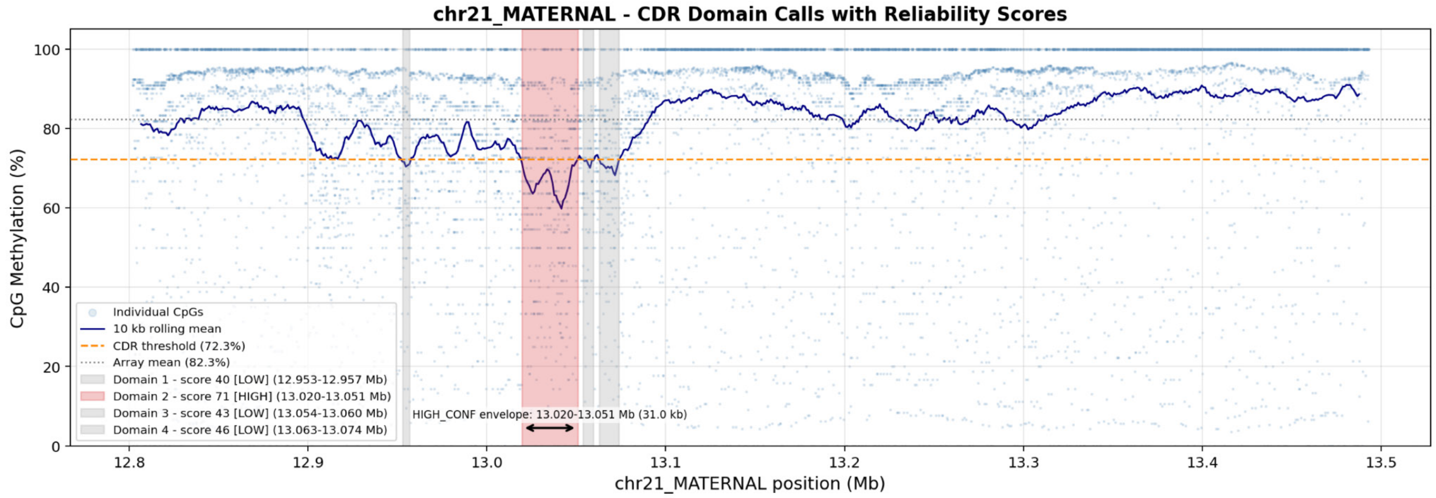


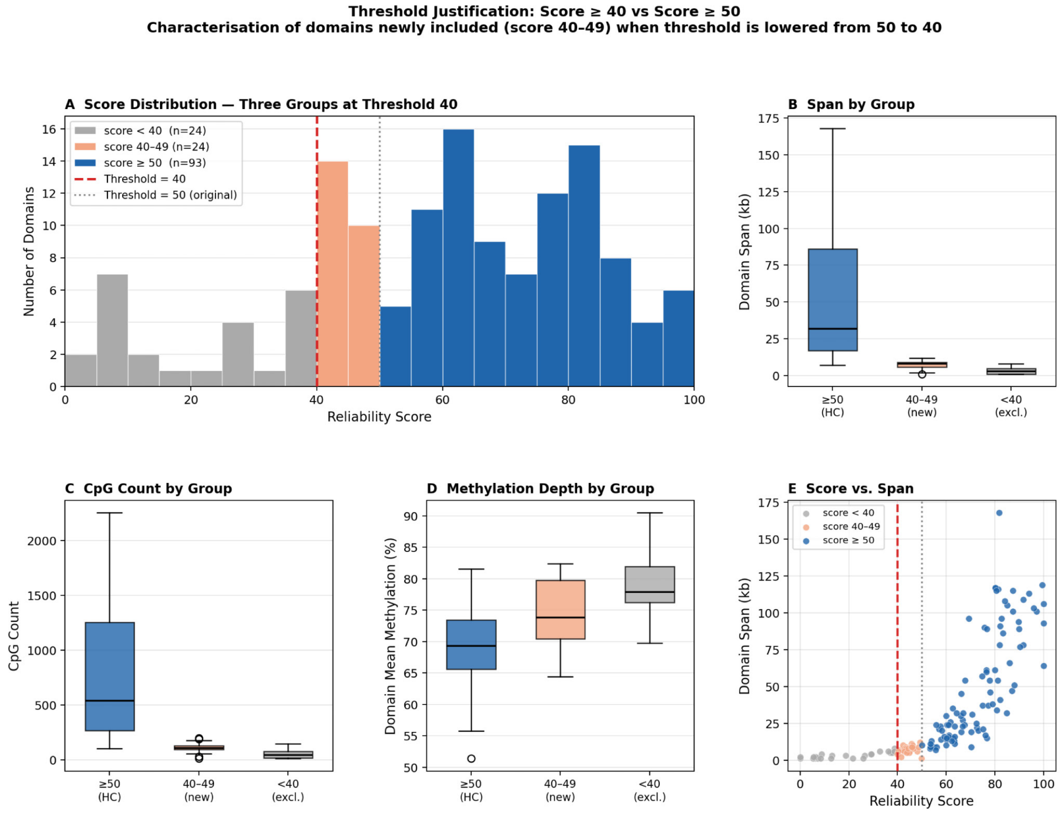

CDR Total Span Definition
The CDR total span is defined as the genomic span from the start of the first HIGH_CONFIDENCE domain to the end of the last HIGH_CONFIDENCE domain on each haplotype. This definition excludes LOW_CONFIDENCE domains at the periphery of the CDR region. The rationale for using the HIGH_CONFIDENCE total span is that peripheral LOW_CONFIDENCE domains are often separated from the main CDR cluster by substantial distances (in some cases >200 kb), inflating the apparent CDR extent by a factor of 2–3×. Restricting the total span to HIGH_CONFIDENCE domains yields a more conservative and biologically defensible estimate of the extent of each CDR.

**Software and Parameters**
CDR calling was performed using *cdr_caller_v2.py*, implemented in Python 3 using NumPy (v1.24+) and Matplotlib (v3.7+). Default parameters were used throughout: sliding window size 10 kb, step size 1 kb, minimum 3 CpGs per window, CDR threshold offset 10 percentage points, and reliability score cutoff 50. The script is available as Supplemental Code.

| **Parameter** | **Value** | **Description** |
|---|---|---|
| --window | 10,000 bp | Sliding window size |
| --step | 1,000 bp | Window step size |
| --min-cpg | 3 | Minimum CpGs required per window |
| --threshold | 10.0% | CDR threshold offset below array mean |
| --min-score | 50 | Reliability score cutoff for HIGH_CONFIDENCE classification |

The caller produces three output files per haplotype array:
- **BED file** (*_CDR_domains.bed): Tab-delimited, no header, compatible with UCSC Genome Browser custom track loading. Columns: chromosome, start, end, domain name, reliability score, strand (.), CpG count, mean methylation, minimum methylation, maximum methylation, confidence classification.
- **Statistics report** (*_CDR_stats.txt): Human-readable summary including run parameters, array-wide mean and threshold, total domain count, HIGH/LOW_CONFIDENCE counts, HIGH_CONFIDENCE and all-domain envelope coordinates, and a per-domain table with full statistics and reliability scores.
- **Visualisation** (*_CDR_plot.png): Per-array methylation plot showing individual CpG values, the 10 kb sliding window mean, the detection threshold, and shaded domain annotations coloured by confidence (HIGH_CONFIDENCE domains in distinct colours; LOW_CONFIDENCE domains in grey). A double-headed arrow annotation marks the HIGH_CONFIDENCE envelope span.

Below we provide the plotted version for all hiPSCs CDRs, sub-CDRs (Domains), and their respective scoring. Scores below 50 are in grey; however, for this paper, we include those scores 40 or higher, as that better represented chr21_MATERNAL (below). Notably, this included small sub-CDR domains, which could be erroneous for chromosome 11_PATERNAL.

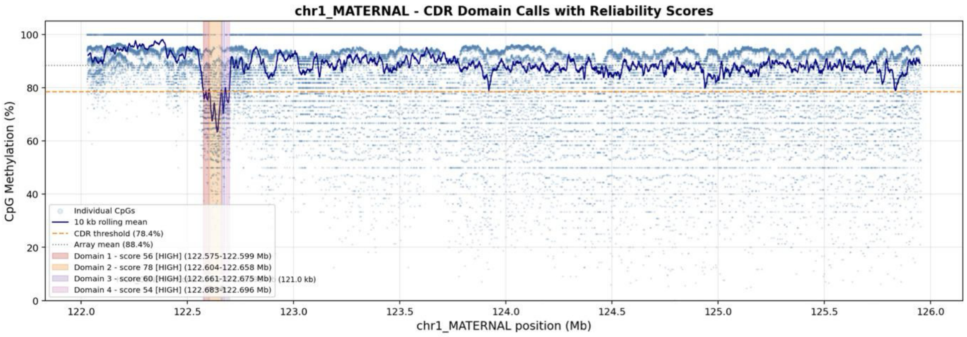
chr1_MATERNAL - CDR Domain Calls with Reliability Scores
CpG Methylation (%)
chr1_MATERNAL position (Mb)
Individual CpGs
10 kb rolling mean
CDR threshold (78.4%)
Array mean (88.4%)
Domain 1 - score 56 [HIGH] (122.575-122.599 Mb)
Domain 2 - score 78 [HIGH] (122.604-122.658 Mb)
Domain 3 - score 60 [HIGH] (122.661-122.675 Mb)
Domain 4 - score 54 [HIGH] (122.683-122.696 Mb)
(121.0 kb)
0
20
40
60
80
100
122.0
122.5
123.0
123.5
124.0
124.5
125.0
125.5
126.0

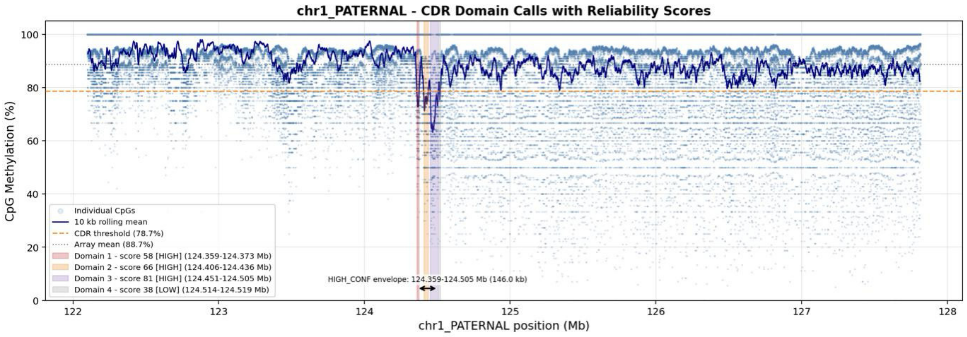
chr1_PATERNAL - CDR Domain Calls with Reliability Scores
CpG Methylation (%)
chr1_PATERNAL position (Mb)
Individual CpGs
10 kb rolling mean
CDR threshold (78.7%)
Array mean (88.7%)
Domain 1 - score 58 [HIGH] (124.359-124.373 Mb)
Domain 2 - score 66 [HIGH] (124.406-124.436 Mb)
Domain 3 - score 81 [HIGH] (124.451-124.505 Mb)
Domain 4 - score 38 [LOW] (124.514-124.519 Mb)
HIGH_CONF envelope: 124.359-124.505 Mb (146.0 kb)
0
20
40
60
80
100
122
123
124
125
126
127
128

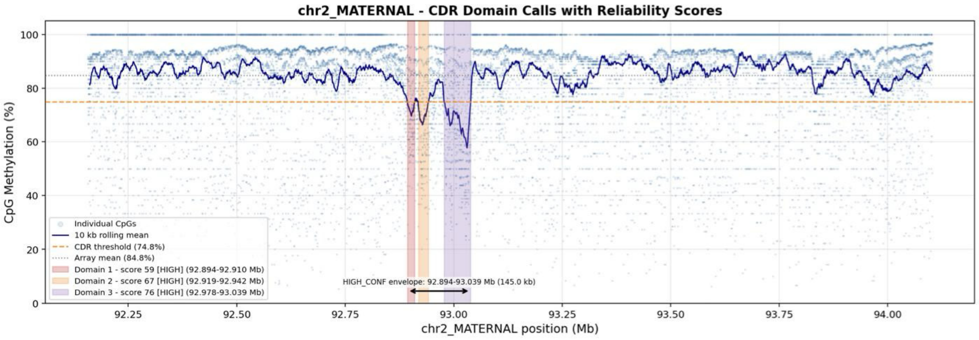
chr2_MATERNAL - CDR Domain Calls with Reliability Scores
CpG Methylation (%)
chr2_MATERNAL position (Mb)
Individual CpGs
10 kb rolling mean
CDR threshold (74.8%)
Array mean (84.8%)
Domain 1 - score 59 [HIGH] (92.894-92.910 Mb)
Domain 2 - score 67 [HIGH] (92.919-92.942 Mb)
Domain 3 - score 76 [HIGH] (92.978-93.039 Mb)
HIGH_CONF envelope: 92.894-93.039 Mb (145.0 kb)
0
20
40
60
80
100
92.25
92.50
92.75
93.00
93.25
93.50
93.75
94.00

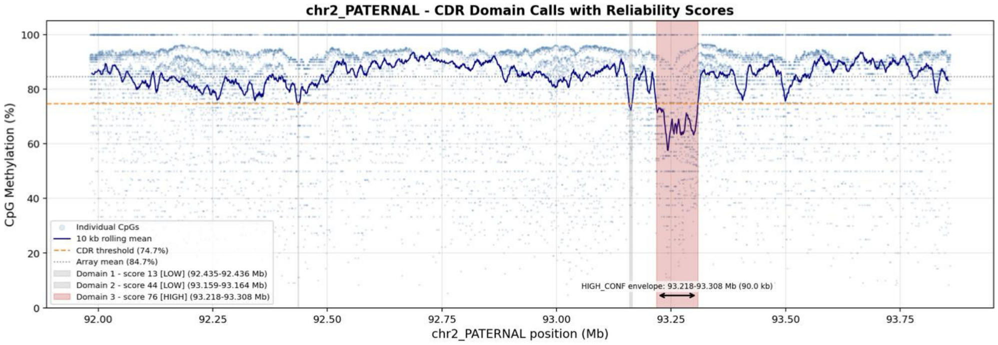
chr2_PATERNAL - CDR Domain Calls with Reliability Scores
CpG Methylation (%)
chr2_PATERNAL position (Mb)
Individual CpGs
10 kb rolling mean
CDR threshold (74.7%)
Array mean (84.7%)
Domain 1 - score 13 [LOW] (92.435-92.436 Mb)
Domain 2 - score 44 [LOW] (93.159-93.164 Mb)
Domain 3 - score 76 [HIGH] (93.218-93.308 Mb)
HIGH_CONF envelope: 93.218-93.308 Mb (90.0 kb)
0
20
40
60
80
100
92.00
92.25
92.50
92.75
93.00
93.25
93.50
93.75

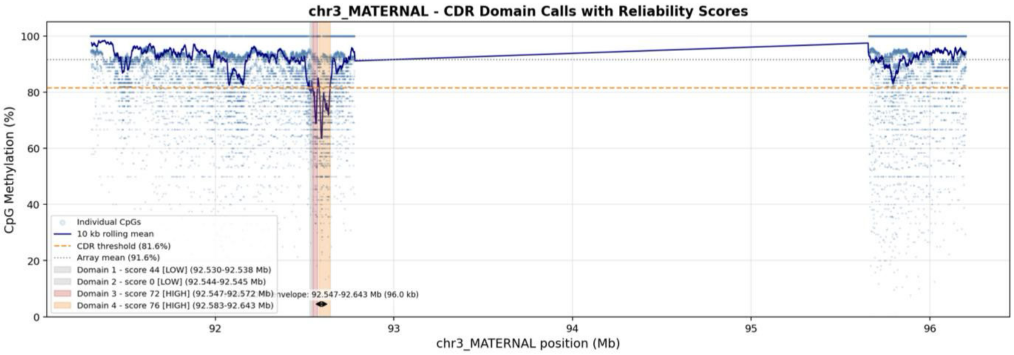
chr3_MATERNAL - CDR Domain Calls with Reliability Scores
CpG Methylation (%)
chr3_MATERNAL position (Mb)
Individual CpGs
10 kb rolling mean
CDR threshold (81.6%)
Array mean (91.6%)
Domain 1 - score 44 [LOW] (92.530-92.538 Mb)
Domain 2 - score 0 [LOW] (92.544-92.545 Mb)
Domain 3 - score 72 [HIGH] (92.547-92.572 Mb)
Domain 4 - score 76 [HIGH] (92.583-92.643 Mb)
nvelope: 92.547-92.643 Mb (96.0 kb)

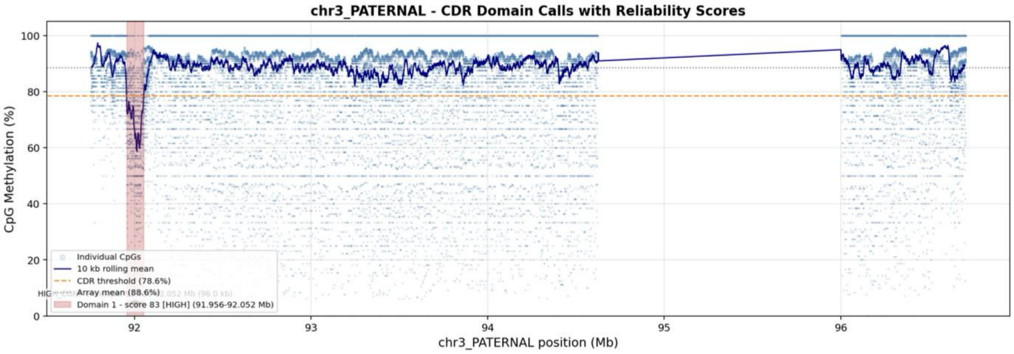
chr3_PATERNAL - CDR Domain Calls with Reliability Scores
CpG Methylation (%)
chr3_PATERNAL position (Mb)
Individual CpGs
10 kb rolling mean
CDR threshold (78.6%)
Array mean (88.6%)
Domain 1 - score 83 [HIGH] (91.956-92.052 Mb)

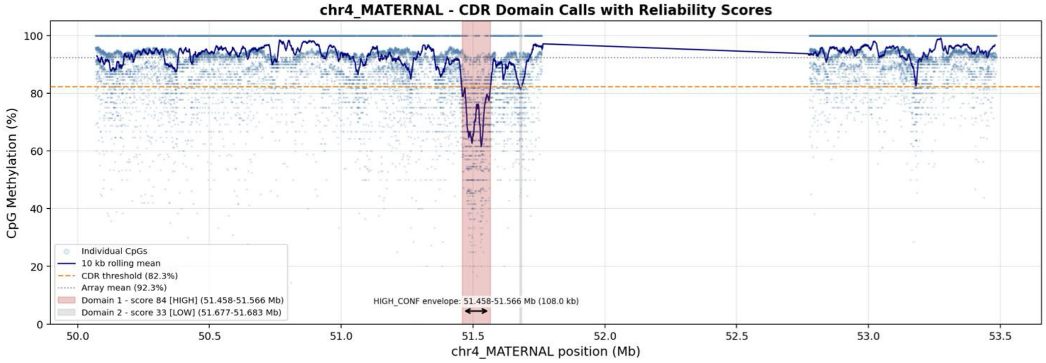
chr4_MATERNAL - CDR Domain Calls with Reliability Scores
CpG Methylation (%)
chr4_MATERNAL position (Mb)
Individual CpGs
10 kb rolling mean
CDR threshold (82.3%)
Array mean (92.3%)
Domain 1 - score 84 [HIGH] (51.458-51.566 Mb)
Domain 2 - score 33 [LOW] (51.677-51.683 Mb)
HIGH_CONF envelope: 51.458-51.566 Mb (108.0 kb)

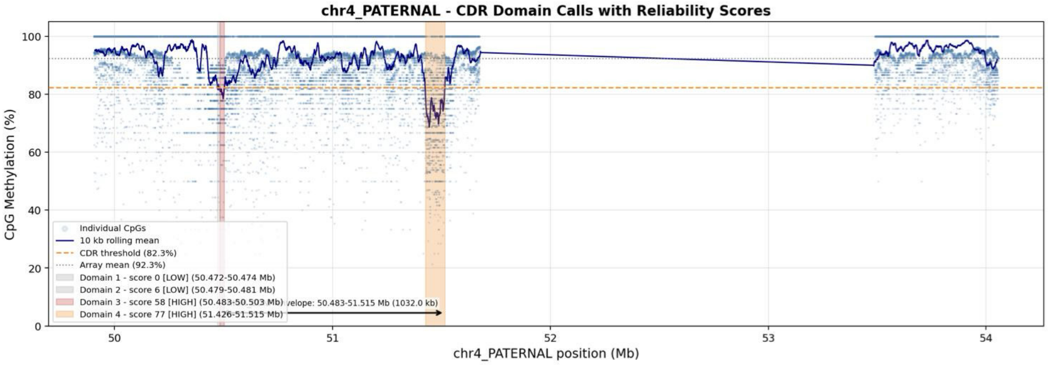
chr4_PATERNAL - CDR Domain Calls with Reliability Scores
CpG Methylation (%)
chr4_PATERNAL position (Mb)
Individual CpGs
10 kb rolling mean
CDR threshold (82.3%)
Array mean (92.3%)
Domain 1 - score 0 [LOW] (50.472-50.474 Mb)
Domain 2 - score 6 [LOW] (50.479-50.481 Mb)
Domain 3 - score 58 [HIGH] (50.483-50.503 Mb)
Domain 4 - score 77 [HIGH] (51.426-51.515 Mb)
velope: 50.483-51.515 Mb (1032.0 kb)

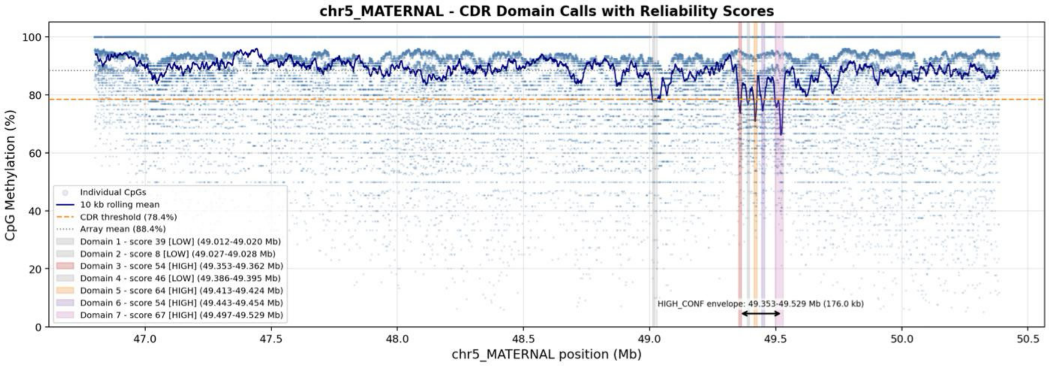
chr5_MATERNAL - CDR Domain Calls with Reliability Scores
CpG Methylation (%)
chr5_MATERNAL position (Mb)
Individual CpGs
10 kb rolling mean
CDR threshold (78.4%)
Array mean (88.4%)
Domain 1 - score 39 [LOW] (49.012-49.020 Mb)
Domain 2 - score 8 [LOW] (49.027-49.028 Mb)
Domain 3 - score 54 [HIGH] (49.353-49.362 Mb)
Domain 4 - score 46 [LOW] (49.386-49.395 Mb)
Domain 5 - score 64 [HIGH] (49.413-49.424 Mb)
Domain 6 - score 54 [HIGH] (49.443-49.454 Mb)
Domain 7 - score 67 [HIGH] (49.497-49.529 Mb)
HIGH_CONF envelope: 49.353-49.529 Mb (176.0 kb)

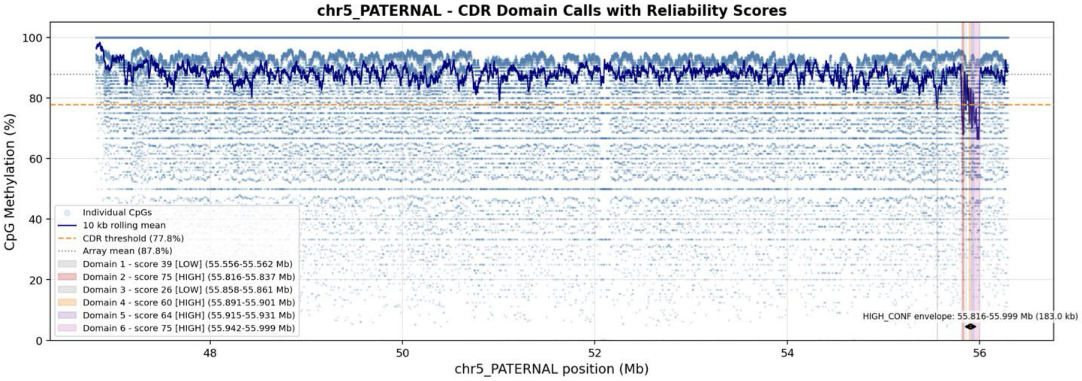
chr5_PATERNAL - CDR Domain Calls with Reliability Scores
CpG Methylation (%)
chr5_PATERNAL position (Mb)
Individual CpGs
10 kb rolling mean
CDR threshold (77.8%)
Array mean (87.8%)
Domain 1 - score 39 [LOW] (55.556-55.562 Mb)
Domain 2 - score 75 [HIGH] (55.816-55.837 Mb)
Domain 3 - score 26 [LOW] (55.858-55.861 Mb)
Domain 4 - score 60 [HIGH] (55.891-55.901 Mb)
Domain 5 - score 64 [HIGH] (55.915-55.931 Mb)
Domain 6 - score 75 [HIGH] (55.942-55.999 Mb)
HIGH_CONF envelope: 55.816-55.999 Mb (183.0 kb)

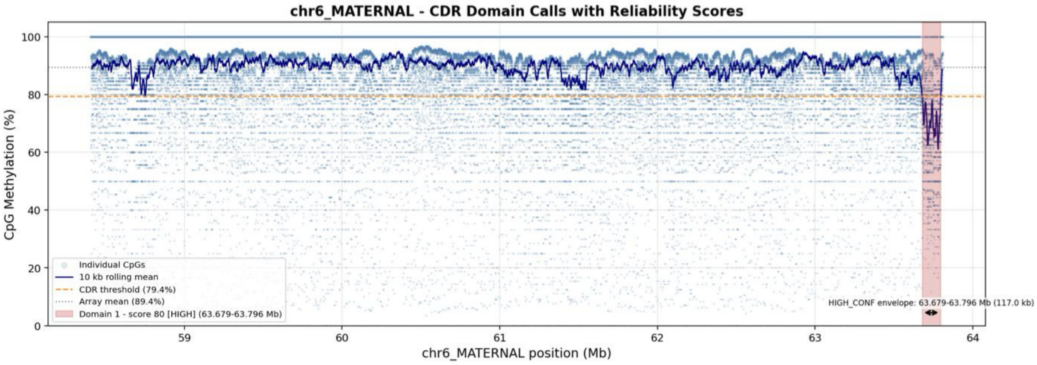
chr6_MATERNAL - CDR Domain Calls with Reliability Scores
CpG Methylation (%)
chr6_MATERNAL position (Mb)
Individual CpGs
10 kb rolling mean
CDR threshold (79.4%)
Array mean (89.4%)
Domain 1 - score 80 [HIGH] (63.679-63.796 Mb)
HIGH_CONF envelope: 63.679-63.796 Mb (117.0 kb)

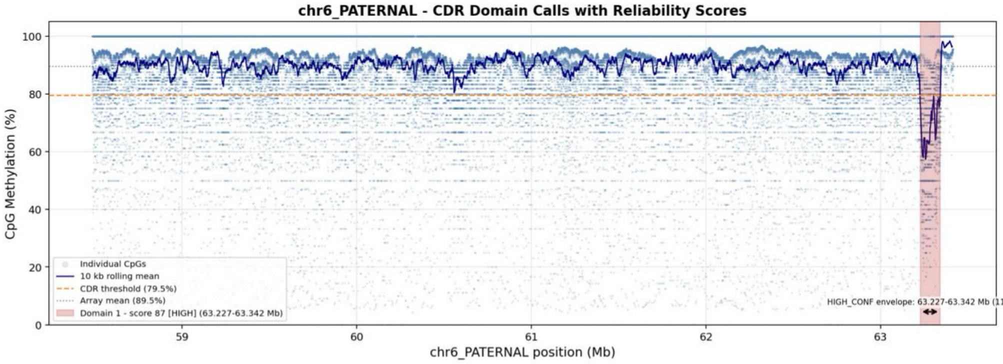
chr6_PATERNAL - CDR Domain Calls with Reliability Scores
CpG Methylation (%)
chr6_PATERNAL position (Mb)
Individual CpGs
10 kb rolling mean
CDR threshold (79.5%)
Array mean (89.5%)
Domain 1 - score 87 [HIGH] (63.227-63.342 Mb)
HIGH_CONF envelope: 63.227-63.342 Mb (115.0 kb)

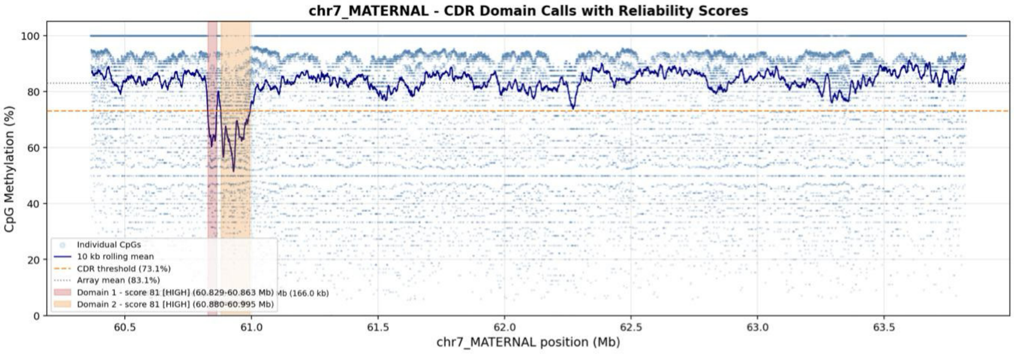
chr7_MATERNAL - CDR Domain Calls with Reliability Scores
CpG Methylation (%)
chr7_MATERNAL position (Mb)
Individual CpGs
10 kb rolling mean
CDR threshold (73.1%)
Array mean (83.1%)
Domain 1 - score 81 [HIGH] (60.829-60.863 Mb)
Domain 2 - score 81 [HIGH] (60.880-60.995 Mb)
Mb (166.0 kb)

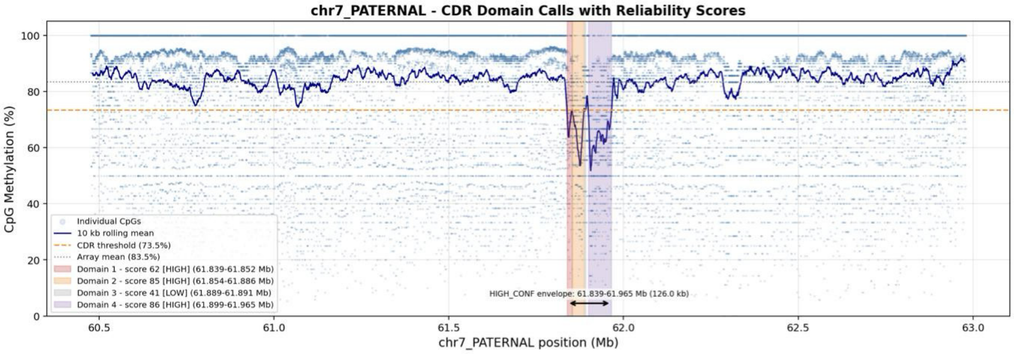
chr7_PATERNAL - CDR Domain Calls with Reliability Scores
CpG Methylation (%)
chr7_PATERNAL position (Mb)
Individual CpGs
10 kb rolling mean
CDR threshold (73.5%)
Array mean (83.5%)
Domain 1 - score 62 [HIGH] (61.839-61.852 Mb)
Domain 2 - score 85 [HIGH] (61.854-61.886 Mb)
Domain 3 - score 41 [LOW] (61.889-61.891 Mb)
Domain 4 - score 86 [HIGH] (61.899-61.965 Mb)
HIGH_CONF envelope: 61.839-61.965 Mb (126.0 kb)

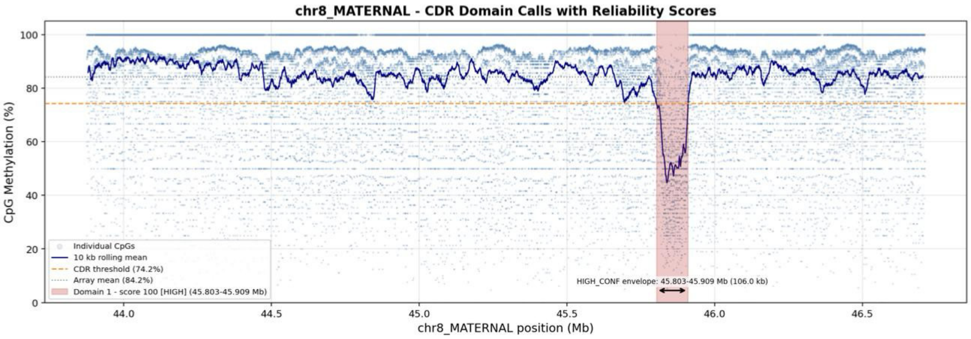
chr8_MATERNAL - CDR Domain Calls with Reliability Scores
CpG Methylation (%)
chr8_MATERNAL position (Mb)
Individual CpGs
10 kb rolling mean
CDR threshold (74.2%)
Array mean (84.2%)
Domain 1 - score 100 [HIGH] (45.803-45.909 Mb)
HIGH_CONF envelope: 45.803-45.909 Mb (106.0 kb)

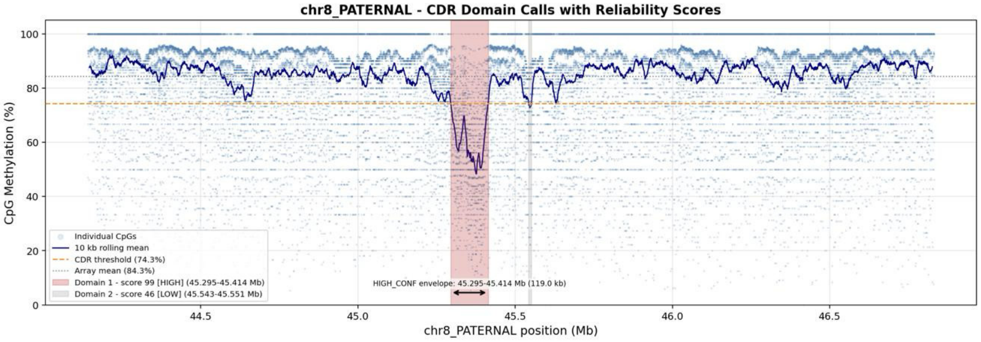
chr8_PATERNAL - CDR Domain Calls with Reliability Scores
CpG Methylation (%)
chr8_PATERNAL position (Mb)
Individual CpGs
10 kb rolling mean
CDR threshold (74.3%)
Array mean (84.3%)
Domain 1 - score 99 [HIGH] (45.295-45.414 Mb)
Domain 2 - score 46 [LOW] (45.543-45.551 Mb)
HIGH_CONF envelope: 45.295-45.414 Mb (119.0 kb)

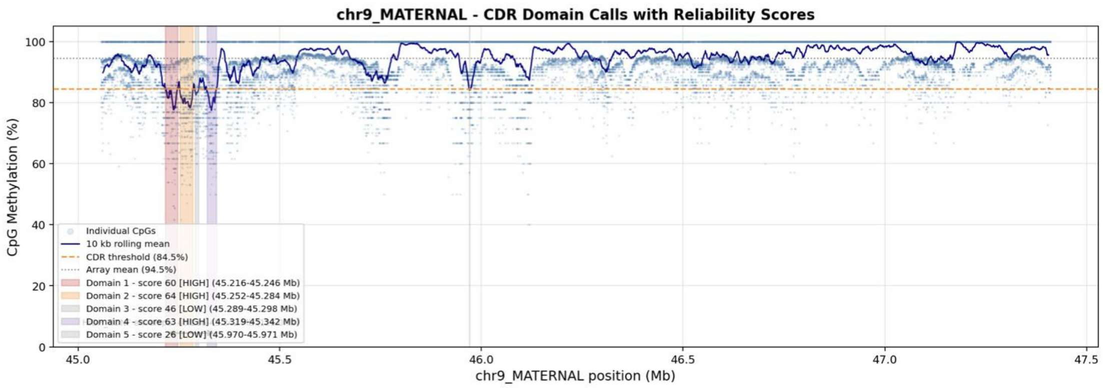
chr9_MATERNAL - CDR Domain Calls with Reliability Scores
CpG Methylation (%)
chr9_MATERNAL position (Mb)
Individual CpGs
10 kb rolling mean
CDR threshold (84.5%)
Array mean (94.5%)
Domain 1 - score 60 [HIGH] (45.216-45.246 Mb)
Domain 2 - score 64 [HIGH] (45.252-45.284 Mb)
Domain 3 - score 46 [LOW] (45.289-45.298 Mb)
Domain 4 - score 63 [HIGH] (45.319-45.342 Mb)
Domain 5 - score 26 [LOW] (45.970-45.971 Mb)

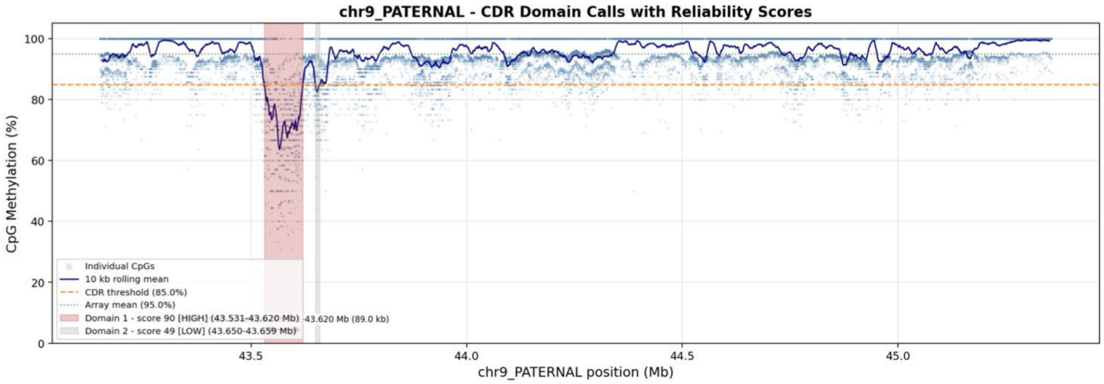
chr9_PATERNAL - CDR Domain Calls with Reliability Scores
CpG Methylation (%)
chr9_PATERNAL position (Mb)
Individual CpGs
10 kb rolling mean
CDR threshold (85.0%)
Array mean (95.0%)
Domain 1 - score 90 [HIGH] (43.531-43.620 Mb)
Domain 2 - score 49 [LOW] (43.650-43.659 Mb)
43.620 Mb (89.0 kb)

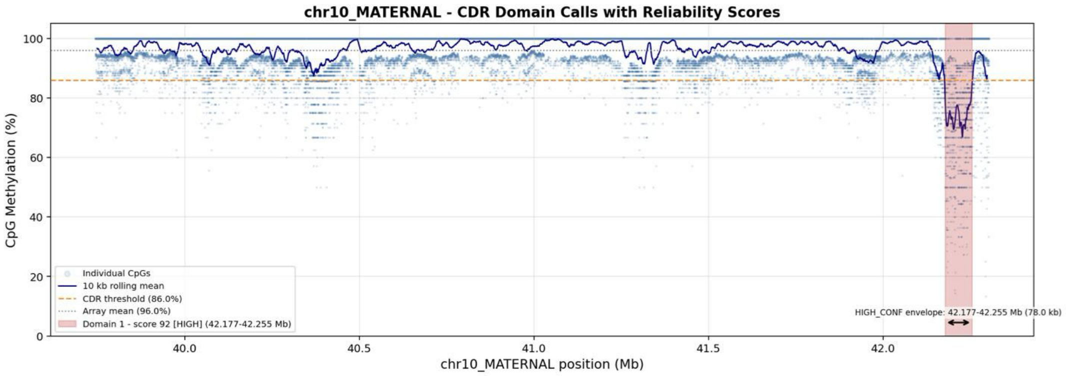
chr10_MATERNAL - CDR Domain Calls with Reliability Scores
CpG Methylation (%)
chr10_MATERNAL position (Mb)
Individual CpGs
10 kb rolling mean
CDR threshold (86.0%)
Array mean (96.0%)
Domain 1 - score 92 [HIGH] (42.177-42.255 Mb)
HIGH_CONF envelope: 42.177-42.255 Mb (78.0 kb)

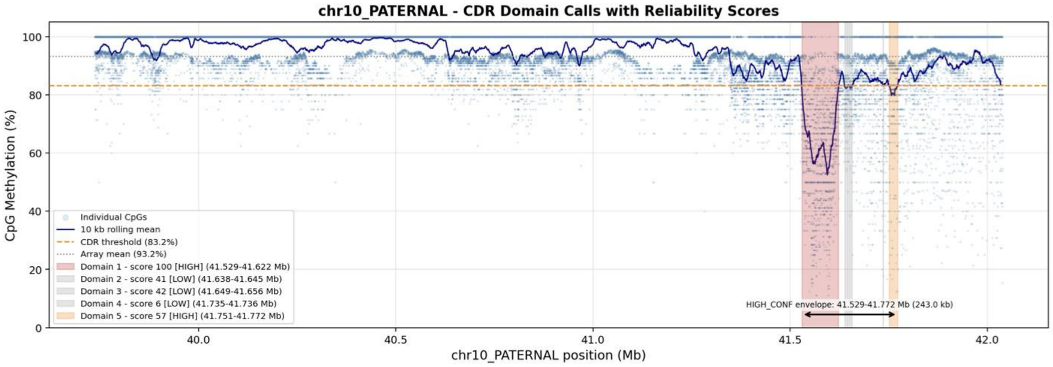
chr10_PATERNAL - CDR Domain Calls with Reliability Scores
CpG Methylation (%)
chr10_PATERNAL position (Mb)
Individual CpGs
10 kb rolling mean
CDR threshold (83.2%)
Array mean (93.2%)
Domain 1 - score 100 [HIGH] (41.529-41.622 Mb)
Domain 2 - score 41 [LOW] (41.638-41.645 Mb)
Domain 3 - score 42 [LOW] (41.649-41.656 Mb)
Domain 4 - score 6 [LOW] (41.735-41.736 Mb)
Domain 5 - score 57 [HIGH] (41.751-41.772 Mb)
HIGH_CONF envelope: 41.529-41.772 Mb (243.0 kb)

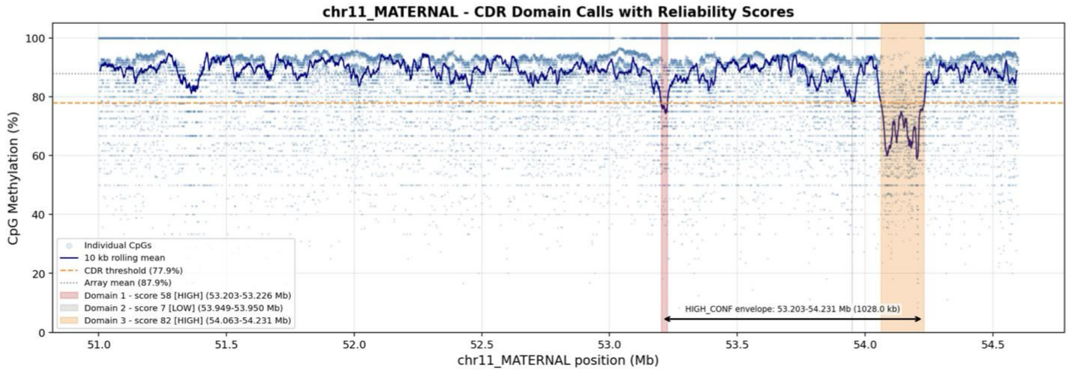
chr11_MATERNAL - CDR Domain Calls with Reliability Scores
CpG Methylation (%)
chr11_MATERNAL position (Mb)
Individual CpGs
10 kb rolling mean
CDR threshold (77.9%)
Array mean (87.9%)
Domain 1 - score 58 [HIGH] (53.203-53.226 Mb)
Domain 2 - score 7 [LOW] (53.949-53.950 Mb)
Domain 3 - score 82 [HIGH] (54.063-54.231 Mb)
HIGH_CONF envelope: 53.203-54.231 Mb (1028.0 kb)

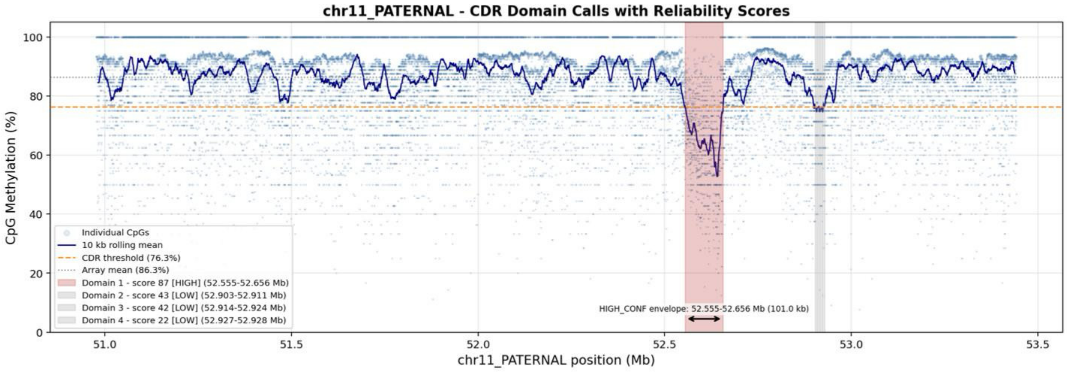
chr11_PATERNAL - CDR Domain Calls with Reliability Scores
CpG Methylation (%)
chr11_PATERNAL position (Mb)
Individual CpGs
10 kb rolling mean
CDR threshold (76.3%)
Array mean (86.3%)
Domain 1 - score 87 [HIGH] (52.555-52.656 Mb)
Domain 2 - score 43 [LOW] (52.903-52.911 Mb)
Domain 3 - score 42 [LOW] (52.914-52.924 Mb)
Domain 4 - score 22 [LOW] (52.927-52.928 Mb)
HIGH_CONF envelope: 52.555-52.656 Mb (101.0 kb)

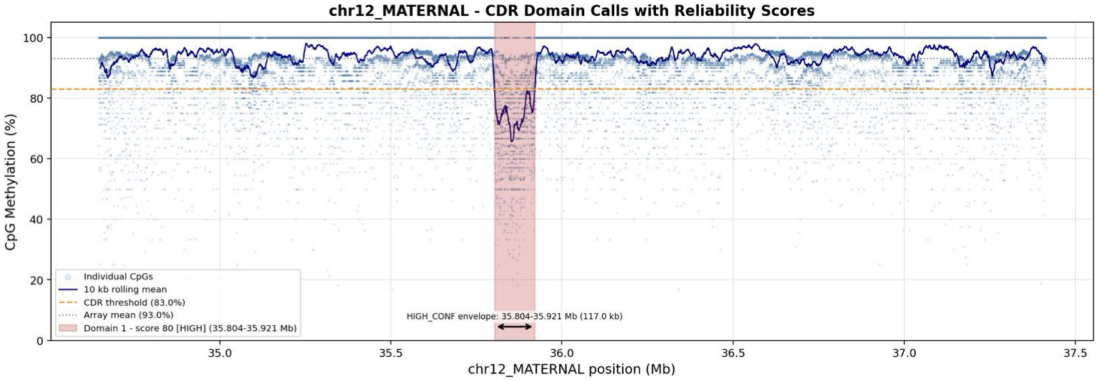
chr12_MATERNAL - CDR Domain Calls with Reliability Scores
CpG Methylation (%)
chr12_MATERNAL position (Mb)
Individual CpGs
10 kb rolling mean
CDR threshold (83.0%)
Array mean (93.0%)
Domain 1 - score 80 [HIGH] (35.804-35.921 Mb)
HIGH_CONF envelope: 35.804-35.921 Mb (117.0 kb)

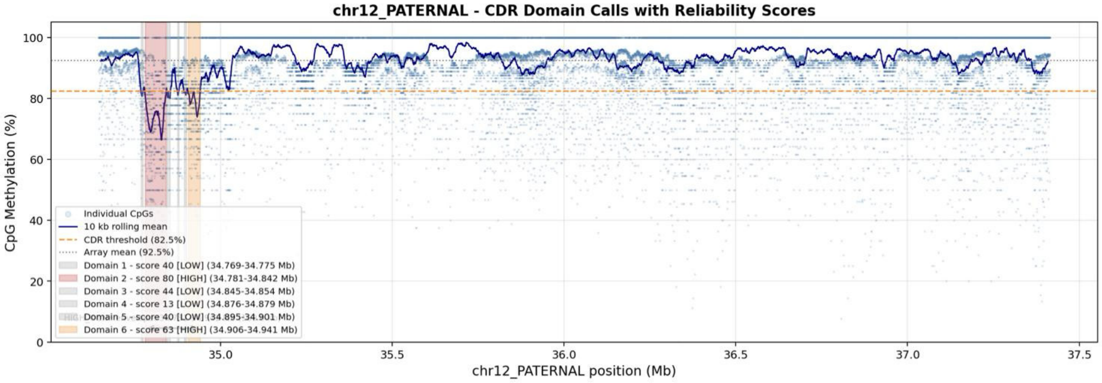
chr12_PATERNAL - CDR Domain Calls with Reliability Scores
CpG Methylation (%)
chr12_PATERNAL position (Mb)
Individual CpGs
10 kb rolling mean
CDR threshold (82.5%)
Array mean (92.5%)
Domain 1 - score 40 [LOW] (34.769-34.775 Mb)
Domain 2 - score 80 [HIGH] (34.781-34.842 Mb)
Domain 3 - score 44 [LOW] (34.845-34.854 Mb)
Domain 4 - score 13 [LOW] (34.876-34.879 Mb)
Domain 5 - score 40 [LOW] (34.895-34.901 Mb)
Domain 6 - score 63 [HIGH] (34.906-34.941 Mb)

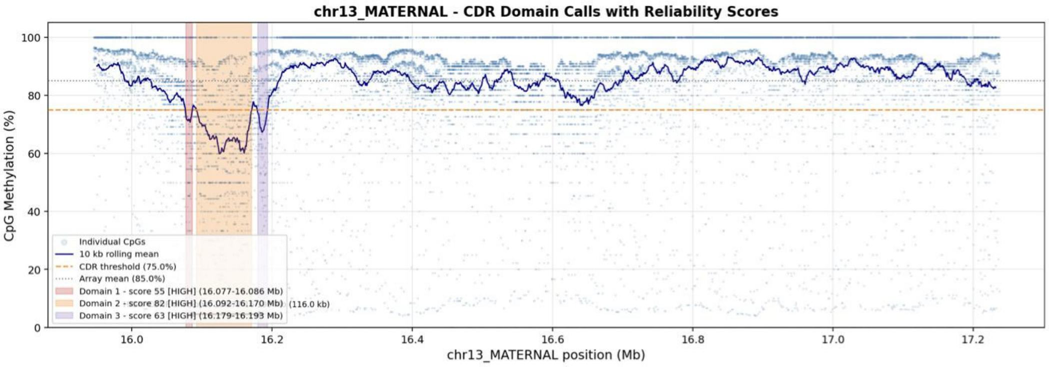
chr13_MATERNAL - CDR Domain Calls with Reliability Scores
CpG Methylation (%)
chr13_MATERNAL position (Mb)
Individual CpGs
10 kb rolling mean
CDR threshold (75.0%)
Array mean (85.0%)
Domain 1 - score 55 [HIGH] (16.077-16.086 Mb)
Domain 2 - score 82 [HIGH] (16.092-16.170 Mb)
(116.0 kb)
Domain 3 - score 63 [HIGH] (16.179-16.193 Mb)

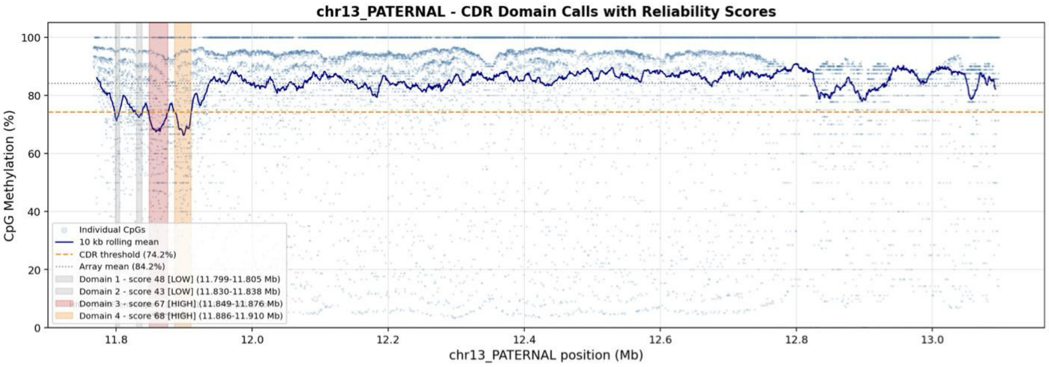
chr13_PATERNAL - CDR Domain Calls with Reliability Scores
CpG Methylation (%)
chr13_PATERNAL position (Mb)
Individual CpGs
10 kb rolling mean
CDR threshold (74.2%)
Array mean (84.2%)
Domain 1 - score 48 [LOW] (11.799-11.805 Mb)
Domain 2 - score 43 [LOW] (11.830-11.838 Mb)
Domain 3 - score 67 [HIGH] (11.849-11.876 Mb)
Domain 4 - score 68 [HIGH] (11.886-11.910 Mb)

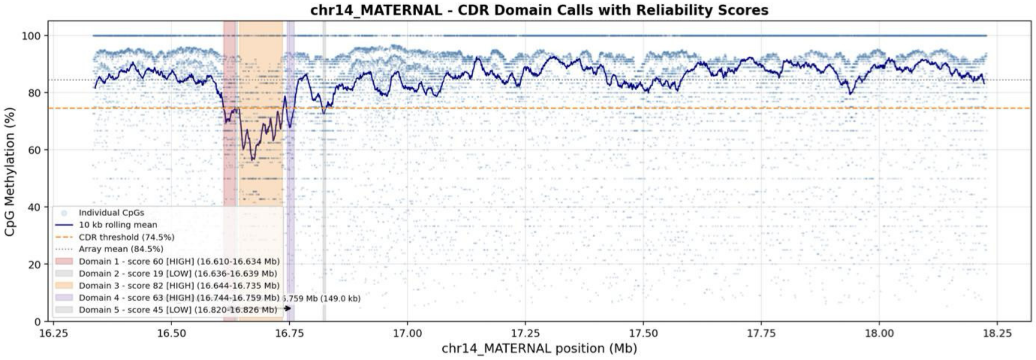
chr14_MATERNAL - CDR Domain Calls with Reliability Scores
CpG Methylation (%)
chr14_MATERNAL position (Mb)
Individual CpGs
10 kb rolling mean
CDR threshold (74.5%)
Array mean (84.5%)
Domain 1 - score 60 [HIGH] (16.610-16.634 Mb)
Domain 2 - score 19 [LOW] (16.636-16.639 Mb)
Domain 3 - score 82 [HIGH] (16.644-16.735 Mb)
Domain 4 - score 63 [HIGH] (16.744-16.759 Mb)
Domain 5 - score 45 [LOW] (16.820-16.826 Mb)
(149.0 kb)

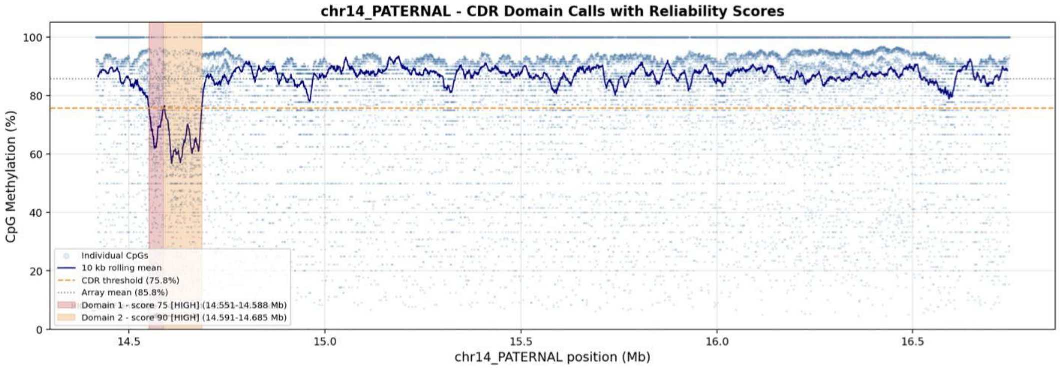
chr14_PATERNAL - CDR Domain Calls with Reliability Scores
CpG Methylation (%)
chr14_PATERNAL position (Mb)
Individual CpGs
10 kb rolling mean
CDR threshold (75.8%)
Array mean (85.8%)
Domain 1 - score 75 [HIGH] (14.551-14.588 Mb)
Domain 2 - score 90 [HIGH] (14.591-14.685 Mb)

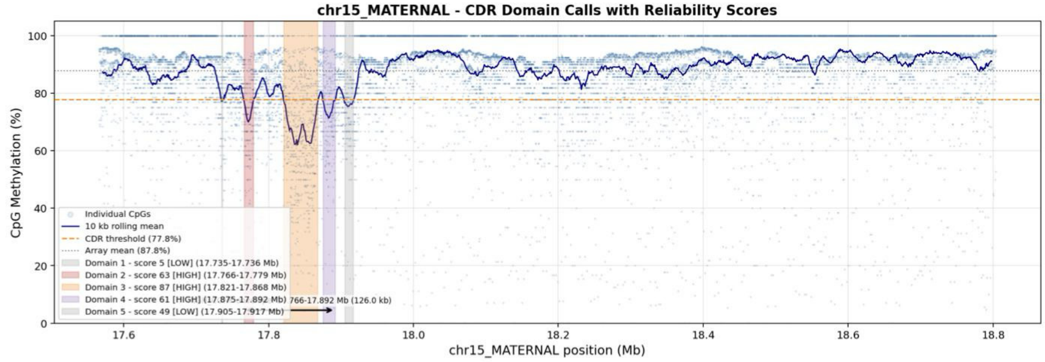
chr15_MATERNAL - CDR Domain Calls with Reliability Scores
CpG Methylation (%)
chr15_MATERNAL position (Mb)
Individual CpGs
10 kb rolling mean
CDR threshold (77.8%)
Array mean (87.8%)
Domain 1 - score 5 [LOW] (17.735-17.736 Mb)
Domain 2 - score 63 [HIGH] (17.766-17.779 Mb)
Domain 3 - score 87 [HIGH] (17.821-17.868 Mb)
Domain 4 - score 61 [HIGH] (17.875-17.892 Mb)
Domain 5 - score 49 [LOW] (17.905-17.917 Mb)
766-17.892 Mb (126.0 kb)

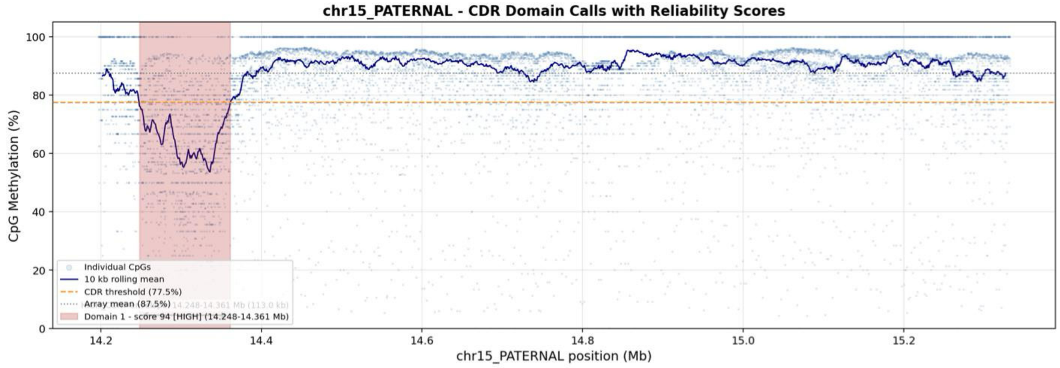
chr15_PATERNAL - CDR Domain Calls with Reliability Scores
CpG Methylation (%)
chr15_PATERNAL position (Mb)
Individual CpGs
10 kb rolling mean
CDR threshold (77.5%)
Array mean (87.5%)
Domain 1 - score 94 [HIGH] (14.248-14.361 Mb)

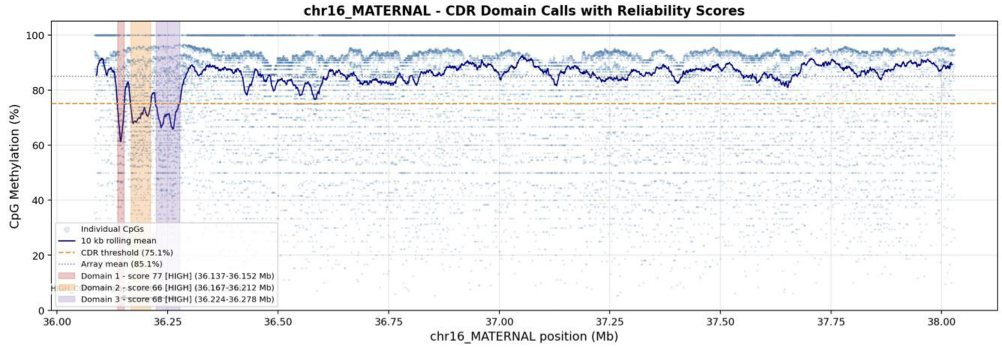
chr16_MATERNAL - CDR Domain Calls with Reliability Scores
CpG Methylation (%)
chr16_MATERNAL position (Mb)
Individual CpGs
10 kb rolling mean
CDR threshold (75.1%)
Array mean (85.1%)
Domain 1 - score 77 [HIGH] (36.137-36.152 Mb)
Domain 2 - score 66 [HIGH] (36.167-36.212 Mb)
Domain 3 - score 68 [HIGH] (36.224-36.278 Mb)

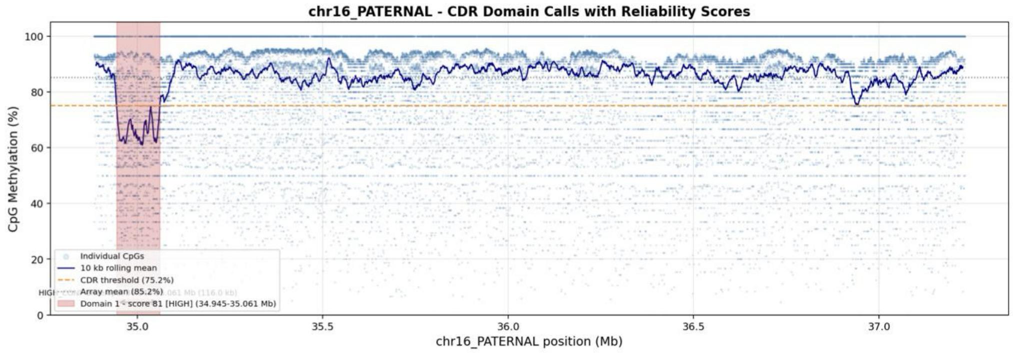
chr16_PATERNAL - CDR Domain Calls with Reliability Scores
CpG Methylation (%)
chr16_PATERNAL position (Mb)
Individual CpGs
10 kb rolling mean
CDR threshold (75.2%)
Array mean (85.2%)
Domain 1 - score 81 [HIGH] (34.945-35.061 Mb)

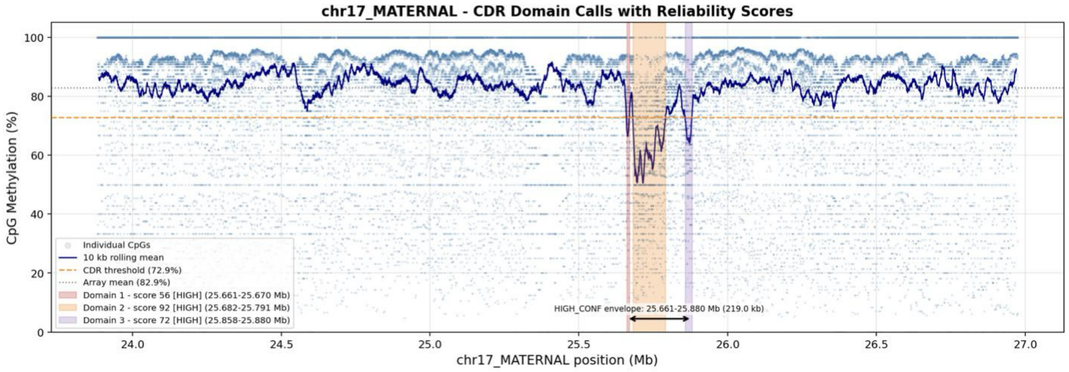
chr17_MATERNAL - CDR Domain Calls with Reliability Scores
CpG Methylation (%)
chr17_MATERNAL position (Mb)
Individual CpGs
10 kb rolling mean
CDR threshold (72.9%)
Array mean (82.9%)
Domain 1 - score 56 [HIGH] (25.661-25.670 Mb)
Domain 2 - score 92 [HIGH] (25.682-25.791 Mb)
Domain 3 - score 72 [HIGH] (25.858-25.880 Mb)
HIGH_CONF envelope: 25.661-25.880 Mb (219.0 kb)

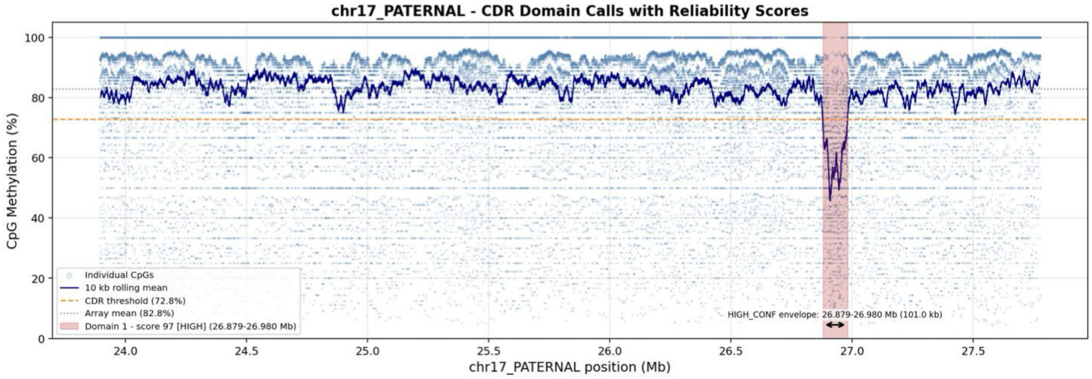
chr17_PATERNAL - CDR Domain Calls with Reliability Scores
CpG Methylation (%)
chr17_PATERNAL position (Mb)
Individual CpGs
10 kb rolling mean
CDR threshold (72.8%)
Array mean (82.8%)
Domain 1 - score 97 [HIGH] (26.879-26.980 Mb)
HIGH_CONF envelope: 26.879-26.980 Mb (101.0 kb)

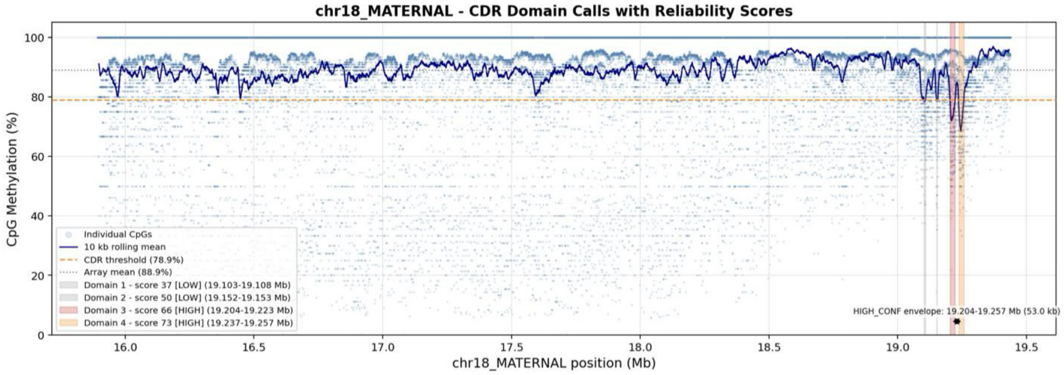
chr18_MATERNAL - CDR Domain Calls with Reliability Scores
CpG Methylation (%)
chr18_MATERNAL position (Mb)
Individual CpGs
10 kb rolling mean
CDR threshold (78.9%)
Array mean (88.9%)
Domain 1 - score 37 [LOW] (19.103-19.108 Mb)
Domain 2 - score 50 [LOW] (19.152-19.153 Mb)
Domain 3 - score 66 [HIGH] (19.204-19.223 Mb)
Domain 4 - score 73 [HIGH] (19.237-19.257 Mb)
HIGH_CONF envelope: 19.204-19.257 Mb (53.0 kb)

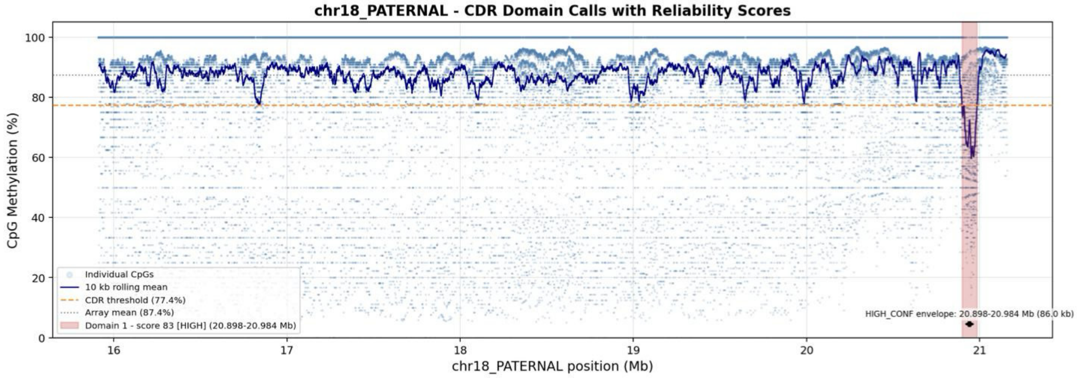
chr18_PATERNAL - CDR Domain Calls with Reliability Scores
CpG Methylation (%)
chr18_PATERNAL position (Mb)
Individual CpGs
10 kb rolling mean
CDR threshold (77.4%)
Array mean (87.4%)
Domain 1 - score 83 [HIGH] (20.898-20.984 Mb)
HIGH_CONF envelope: 20.898-20.984 Mb (86.0 kb)

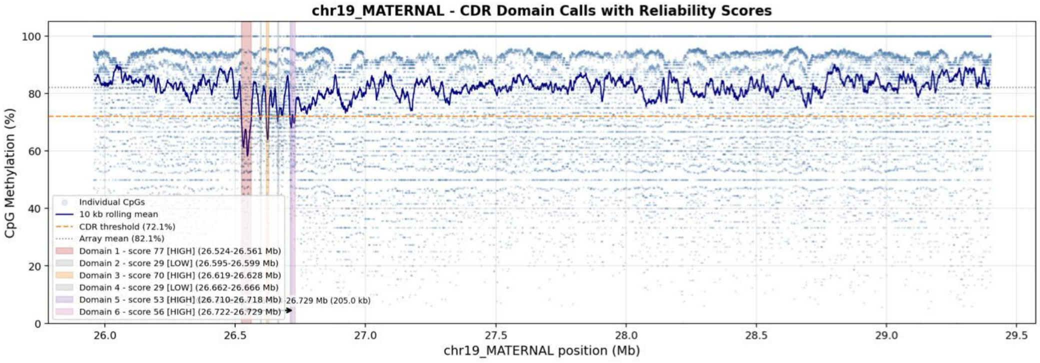
chr19_MATERNAL - CDR Domain Calls with Reliability Scores
CpG Methylation (%)
chr19_MATERNAL position (Mb)
Individual CpGs
10 kb rolling mean
CDR threshold (72.1%)
Array mean (82.1%)
Domain 1 - score 77 [HIGH] (26.524-26.561 Mb)
Domain 2 - score 29 [LOW] (26.595-26.599 Mb)
Domain 3 - score 70 [HIGH] (26.619-26.628 Mb)
Domain 4 - score 29 [LOW] (26.662-26.666 Mb)
Domain 5 - score 53 [HIGH] (26.710-26.718 Mb)
Domain 6 - score 56 [HIGH] (26.722-26.729 Mb)
26.729 Mb (205.0 kb)

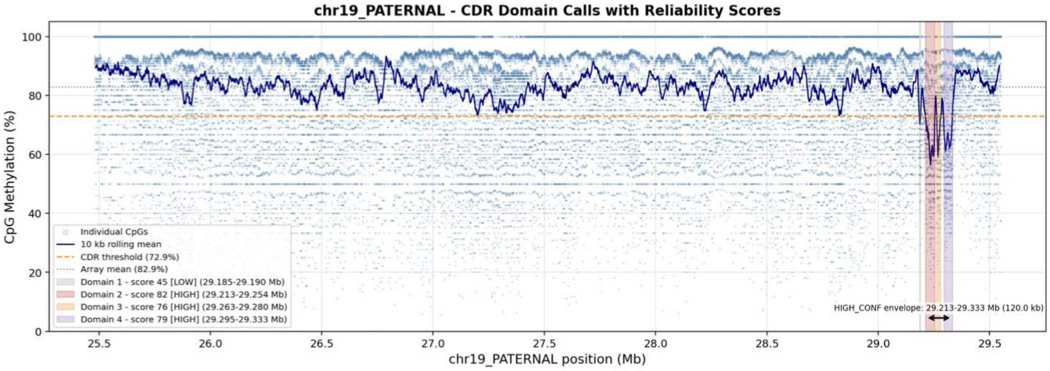
chr19_PATERNAL - CDR Domain Calls with Reliability Scores
CpG Methylation (%)
chr19_PATERNAL position (Mb)
Individual CpGs
10 kb rolling mean
CDR threshold (72.9%)
Array mean (82.9%)
Domain 1 - score 45 [LOW] (29.185-29.190 Mb)
Domain 2 - score 82 [HIGH] (29.213-29.254 Mb)
Domain 3 - score 76 [HIGH] (29.263-29.280 Mb)
Domain 4 - score 79 [HIGH] (29.295-29.333 Mb)
HIGH_CONF envelope: 29.213-29.333 Mb (120.0 kb)

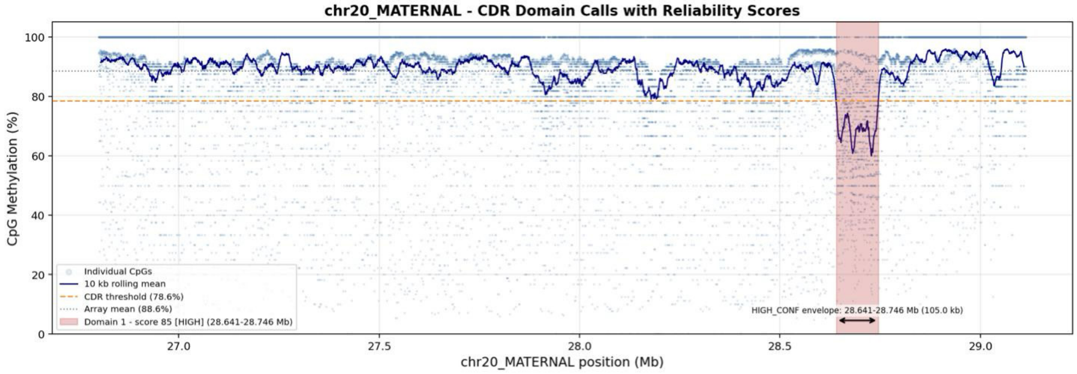
chr20_MATERNAL - CDR Domain Calls with Reliability Scores
CpG Methylation (%)
chr20_MATERNAL position (Mb)
Individual CpGs
10 kb rolling mean
CDR threshold (78.6%)
Array mean (88.6%)
Domain 1 - score 85 [HIGH] (28.641-28.746 Mb)
HIGH_CONF envelope: 28.641-28.746 Mb (105.0 kb)

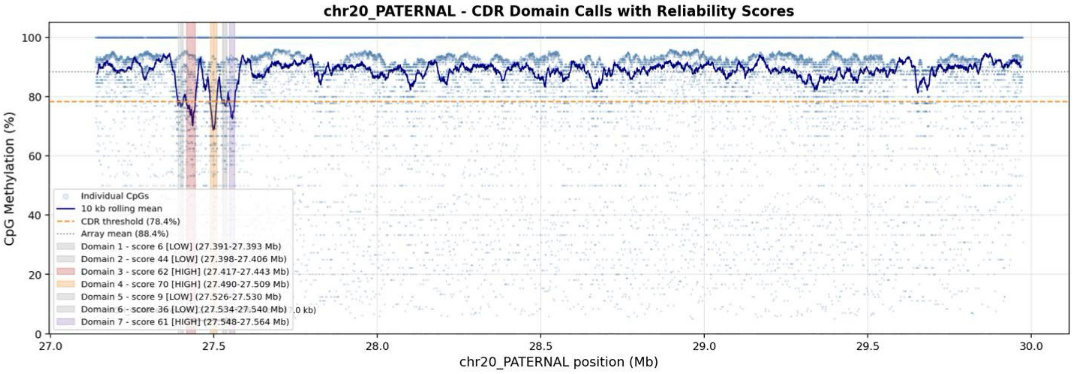
chr20_PATERNAL - CDR Domain Calls with Reliability Scores
CpG Methylation (%)
chr20_PATERNAL position (Mb)
Individual CpGs
10 kb rolling mean
CDR threshold (78.4%)
Array mean (88.4%)
Domain 1 - score 6 [LOW] (27.391-27.393 Mb)
Domain 2 - score 44 [LOW] (27.398-27.406 Mb)
Domain 3 - score 62 [HIGH] (27.417-27.443 Mb)
Domain 4 - score 70 [HIGH] (27.490-27.509 Mb)
Domain 5 - score 9 [LOW] (27.526-27.530 Mb)
Domain 6 - score 36 [LOW] (27.534-27.540 Mb)
Domain 7 - score 61 [HIGH] (27.548-27.564 Mb)

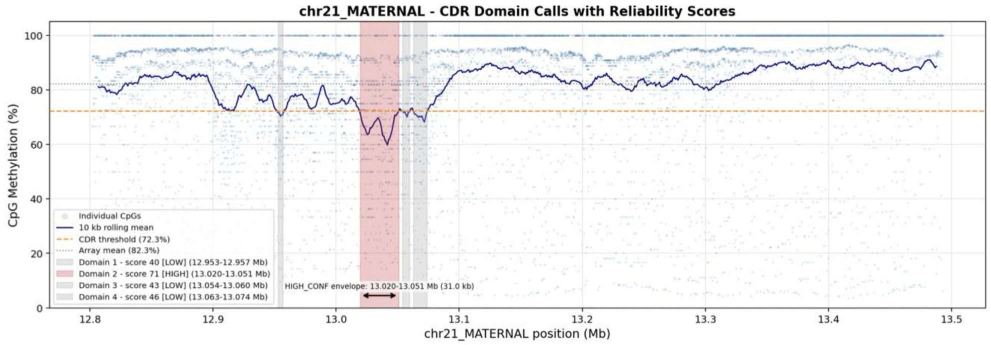
chr21_MATERNAL - CDR Domain Calls with Reliability Scores
CpG Methylation (%)
chr21_MATERNAL position (Mb)
Individual CpGs
10 kb rolling mean
CDR threshold (72.3%)
Array mean (82.3%)
Domain 1 - score 40 [LOW] (12.953-12.957 Mb)
Domain 2 - score 71 [HIGH] (13.020-13.051 Mb)
Domain 3 - score 43 [LOW] (13.054-13.060 Mb)
Domain 4 - score 46 [LOW] (13.063-13.074 Mb)
HIGH_CONF envelope: 13.020-13.051 Mb (31.0 kb)

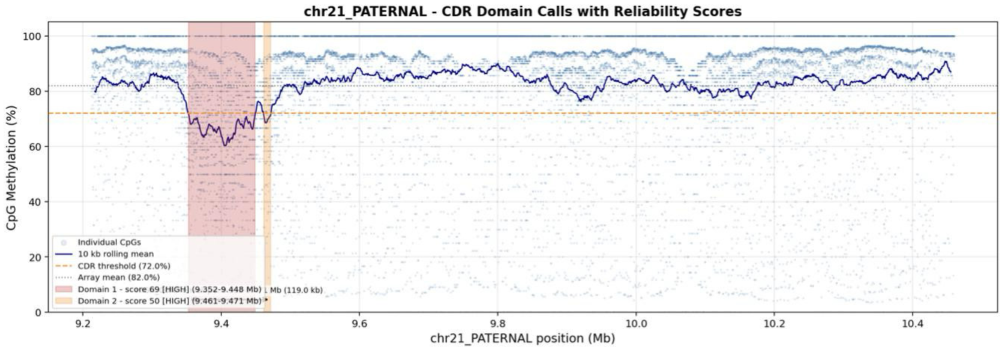
chr21_PATERNAL - CDR Domain Calls with Reliability Scores
CpG Methylation (%)
chr21_PATERNAL position (Mb)
Individual CpGs
10 kb rolling mean
CDR threshold (72.0%)
Array mean (82.0%)
Domain 1 - score 69 [HIGH] (9.352-9.448 Mb)
Domain 2 - score 50 [HIGH] (9.461-9.471 Mb)
Mb (119.0 kb)

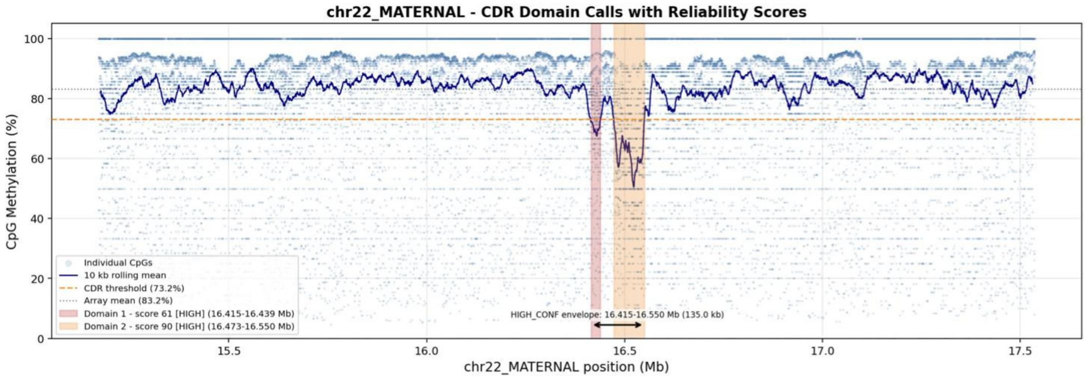
chr22_MATERNAL - CDR Domain Calls with Reliability Scores
CpG Methylation (%)
chr22_MATERNAL position (Mb)
Individual CpGs
10 kb rolling mean
CDR threshold (73.2%)
Array mean (83.2%)
Domain 1 - score 61 [HIGH] (16.415-16.439 Mb)
Domain 2 - score 90 [HIGH] (16.473-16.550 Mb)
HIGH_CONF envelope: 16.415-16.550 Mb (135.0 kb)

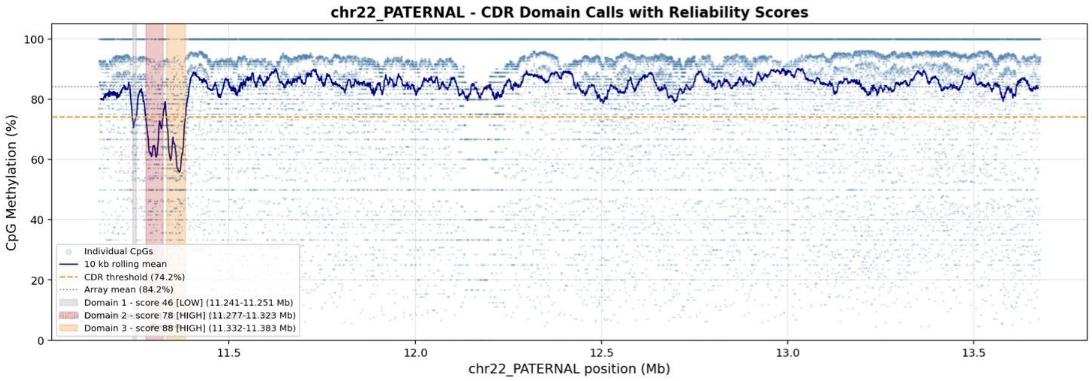
chr22_PATERNAL - CDR Domain Calls with Reliability Scores
CpG Methylation (%)
chr22_PATERNAL position (Mb)
Individual CpGs
10 kb rolling mean
CDR threshold (74.2%)
Array mean (84.2%)
Domain 1 - score 46 [LOW] (11.241-11.251 Mb)
Domain 2 - score 78 [HIGH] (11.277-11.323 Mb)
Domain 3 - score 88 [HIGH] (11.332-11.383 Mb)

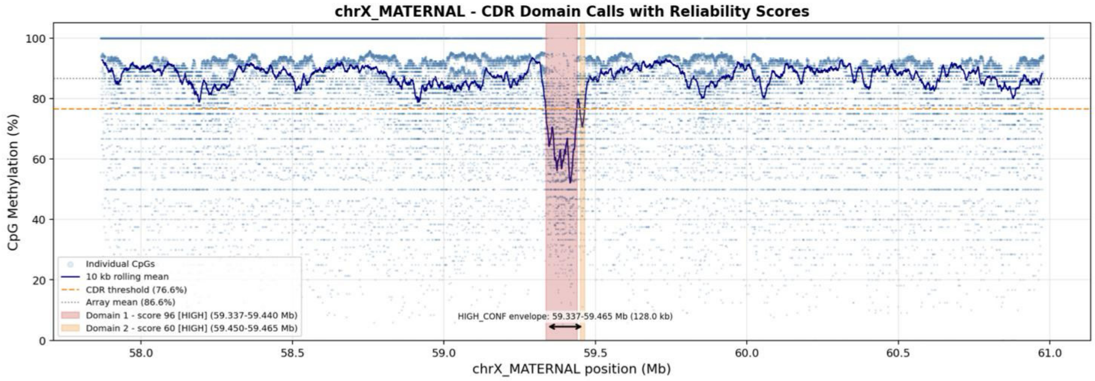
chrX_MATERNAL - CDR Domain Calls with Reliability Scores
CpG Methylation (%)
100
80
60
40
20
0
Individual CpGs
10 kb rolling mean
CDR threshold (76.6%)
Array mean (86.6%)
Domain 1 - score 96 [HIGH] (59.337-59.440 Mb)
Domain 2 - score 60 [HIGH] (59.450-59.465 Mb)
HIGH_CONF envelope: 59.337-59.465 Mb (128.0 kb)
58.0
58.5
59.0
59.5
60.0
60.5
61.0
chrX_MATERNAL position (Mb)

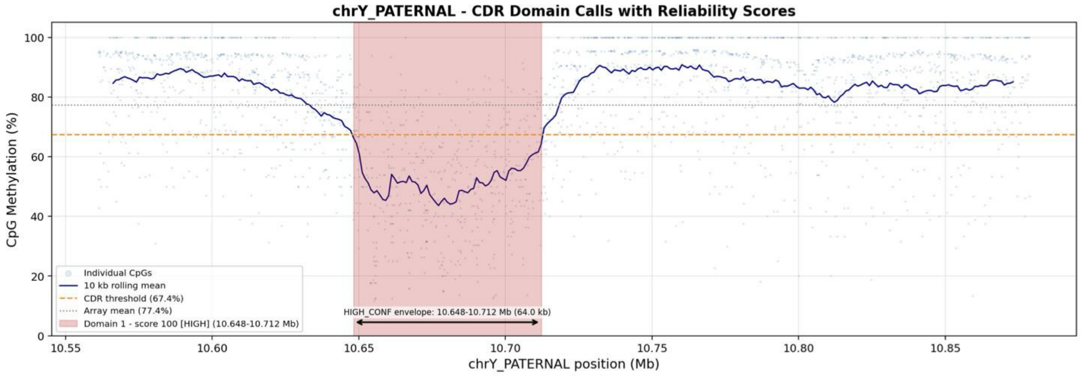
chrY_PATERNAL - CDR Domain Calls with Reliability Scores
CpG Methylation (%)
100
80
60
40
20
0
Individual CpGs
10 kb rolling mean
CDR threshold (67.4%)
Array mean (77.4%)
Domain 1 - score 100 [HIGH] (10.648-10.712 Mb)
HIGH_CONF envelope: 10.648-10.712 Mb (64.0 kb)
10.55
10.60
10.65
10.70
10.75
10.80
10.85
chrY_PATERNAL position (Mb)
